# Species delimitation using genome-wide estimates of *Dxy* and *Fst*

**DOI:** 10.64898/2025.12.08.692692

**Authors:** Menno J. de Jong, Christopher Wambach, Malambo Muloongo, Bruno Lopes da Silva Ferrette, Magnus Wolf, Axel Janke

## Abstract

Accurate and stable species delimitation is essential for effective nature conservation and management. Advances in genomics now offer the potential for consistent decision-making. We performed a meta-analysis of genomic data from hundreds of sister lineages of large mammals to test whether species boundaries can be identified using genome-wide sequence dissimilarity estimates. We find that a combined threshold of absolute (*Dxy* = 0.225% [0.20%-0.27%]) and relative genetic distance (*Fst* = 0.26 [0.19-0.54]) predicts current taxonomic status with over 90% accuracy. This predictive power indicates that mammalian taxonomists managed to maintain consistency across taxa despite having to rely on disparate morphological traits. Lineage pairs exhibiting Haldane’s Rule exceed far higher thresholds (*Dxy* = 0.4%; *Fst* = 0.55), implying that mammalian taxonomists adhere to species concepts which allow for interbreeding. Our findings challenge the significance of fast-track bottleneck speciation and suggest instead that deep divergences of up to 100,000 generations are usually required for mammalian sister lineages to remain genetically isolated upon secondary contact. We discuss taxonomic revisions to improve temporal banding, including three potential cases of cryptic speciation: southern versus eastern aardwolves, Asiatic black bears versus Japanese black bears, and European versus Asian wild boar. In summary, our meta-analysis provides simple rules for species delimitation and offers new insights into the taxonomy and speciation dynamics of large mammals.

## INTRODUCTION

A species is the basal level of taxonomy and the standard unit for quantifying biodiversity. Yet what exactly defines a species is the subject of an ongoing debate involving dozens of competing species concepts (Harrison 1998; Baker and Bradley 2006; Hey 2006; De Queiroz 2007; Zachos 2016). This broader species debate, which ultimately questions whether species truly exist as entities outside the human mind (Mayr 1996; Avise and Walker 1999; Hey et al. 2003; Hey and Pinho 2012), underlies endless discussions about the status of individual lineages. A shared concern among biologists is that the lack of agreement promotes taxonomic inconsistency and instability, thereby hindering the implementation and evaluation of effective nature conservation (Chaitra et al. 2004; Isaac et al. 2004; Garnett and Christidis 2017; Padial and De la Riva 2021; Dufresnes et al. 2023).

The species problem involves three central questions: (1) What is a species? (2) Why do species exist? and (3) How to delimit species? (Mayr 1996; De Queiroz 2007) Answering the third question is a routine yet crucial task for natural historians, who must frequently decide whether an unfamiliar organism represents merely a variety of a previously described species, or instead a new species altogether. Traditionally, such judgments relied on observable morphological features, but the rise of genetics has transformed taxonomy: the rank of known organisms is being re-examined, with decisions increasingly relying on genetic evidence (Wang et al. 1999; Roca et al. 2001; Ludt et al. 2004; Buckley-Beason et al. 2006; Rannala and Yang 2020; Costa et al. 2022; Morin et al. 2024; Muneza et al. 2025).

We set out to evaluate the renewed promise of one of the oldest genetic methods for species delimitation, applied to modern datasets of complete nuclear genomes. Namely, we tested the hypothesis that a pair of lineages may be diagnosed as species if their genetic distance falls within the range typically observed for sister species (Avise et al. 1998; Avise and Walker 1999; Bradley and Baker 2001; Hey and Pinho 2012; Allio et al. 2021; Weber et al. 2021; Lebedev et al. 2025).

Previous attempts to delimit species using a genetic distance threshold inferred from mtDNA data proved challenging owing to the non-recombining nature of the mitochondrial genome (Avise et al. 1998; Avise and Walker 1999; Bradley and Baker 2001; Curnoe and Thorne 2003; Funk and Omland 2003; Groves 2004; Hebert et al. 2004; Moritz and Cicero 2004; Morin et al. 2023). In contrast, nuclear genomes consist of thousands of independently segregating loci, which together rule out stochastic discordant phylogenetic signals that complicate single-locus inferences.

Species delimitation based on a genetic distance threshold is historically associated with the genetic species concept, which posits that two sister lineages are separate species if interbreeding occurs too infrequently to homogenise gene pools even when geographic ranges overlap (Baker and Bradley 2006) (Table 1). For allopatric lineages, in which such genetic isolation cannot be directly assessed, the genetic species concept provides a convenient work-around solution. For a pair of lineages to be considered separate species, it must be demonstrated, using genetic distance estimates, that their divergence is similar to values typically observed for sympatric lineages that are genetically isolated. In practice, this means that genetic distances between lineages ought to be indicative of sufficiently deep divergence times required to establish some form of isolating mechanism (Mallet and Mullen 2022; Dufresnes et al. 2023). The genetic species concept therefore overlaps with both the biological species concept (Mayr 1996; Ayala and Fitch 1997; Baker and Bradley 2006; Lebedev et al. 2025) and the genotypic cluster concept (Mallet 1995; Mallet 2020) (Table 1), while overcoming the major hurdle of poor diagnosability when species ranges do not overlap (Groves 2004; Groves 2013).

**Table 1.** Species concepts. Overview of an arbitrary selection of species concepts and their associated species delimitation methods, discussed in this manuscript.

| Species concept | What is a species? | Interbreeding in sympatry? | How to delimit species? |
| --- | --- | --- | --- |
| Biological (BSC), *original* (Dobzhansky, 1935) | a meta-population that is *reproductively* isolated from other meta-populations | not possible | Test for reproductive isolation, i.e. inability to produce viable and fertile hybrids (*ex-situ cross-breeding or in-vitro fertilisation experiments*) |
| Biological (BSC), *revised* (Mayr, 1970) | a meta-population that is *genetically* isolated from other meta-populations | possible, but restricted (sympatric gene pools do not homogenise) | Test for genetic isolation where ranges overlap: how frequent are F1 hybrids and backcrosses? (*genetic hybrid detection*) |
| Genotypic cluster (GCSC) (Mallet, 1995) | idem | idem | Test for genetic isolation where lineages co-occur: do sympatric lineages form distinct clusters? (*PCA, Admixture, Tree topology*) |
| Genetic (GSC) (Bradley and Baker, 2001) | idem | idem | Test if genetic distance is in the range typically observed for genetically isolated sister lineages (*Distance matrix, Tree branch lengths, convergence of PSMC-curves*) |
| Phylogenetic (PSC) (Baum, 1992) | a meta-population in which all individuals share one or more fixed derived phenotypic traits or alleles | possible, no explicitly stated restrictions | Test for reciprocal monophyly (*e.g., single-locus analyses, such as mtDNA-based inferences*) |
| Morphological (MSC) (Cronquist, 1978) | a meta-population in which all individuals share a set of distinct phenotypic traits | possible, no explicitly stated restrictions | Test for a discontinuous distribution of physical traits |

In order to construct the required reference database and to identify potential threshold values, we calculated genetic distances for a large number of closely related lineages. We included only taxonomic clades with roughly comparable life history traits, namely large placental mammals with generation times of over four years. Data for over 150 lineages belonging to approximately 100 species and 50 genera (Fig. S1A, Table S1) provided sufficient statistical power to draw general conclusions.

To ensure comparability, all datasets were analysed using a uniform protocol for short read filtering, short read mapping, and genotype calling. Unlike the exome-focused approach of Lebedev et al (2025), we analysed entire genomes, with the assumption that functional regions constitute a relatively small portion of the genome such that the effect of direct selection is negligible. Admittedly, linked selection affects much larger proportions of genomes, but only alters genetic diversity within lineages, not absolute genetic distances among lineages (Birky and Walsh 1988; Charlesworth 1998; Menno J. de Jong et al. 2024).

For each pair of closely related lineages, we calculated two measures of population-level differentiation, namely absolute distance (*Dxy*) and relative distance (*Fst*) (Fig. 1) (Cruickshank and Hahn 2014). The former, which can be considered a measure of gross distance, simply denotes the mean sequence dissimilarity between a pair of lineages (Fig. 1). The latter, also known as the fixation index, normalises net divergence (*Da,* approximated by *Nei’s D,* Fig. S1B). We calculated *Fst*-estimates using the formula of Hudson et al. (1992) (Fig. 1), which is particularly well-suited for small sample sizes (M. J. de Jong et al. 2024).

**Figure 1.**
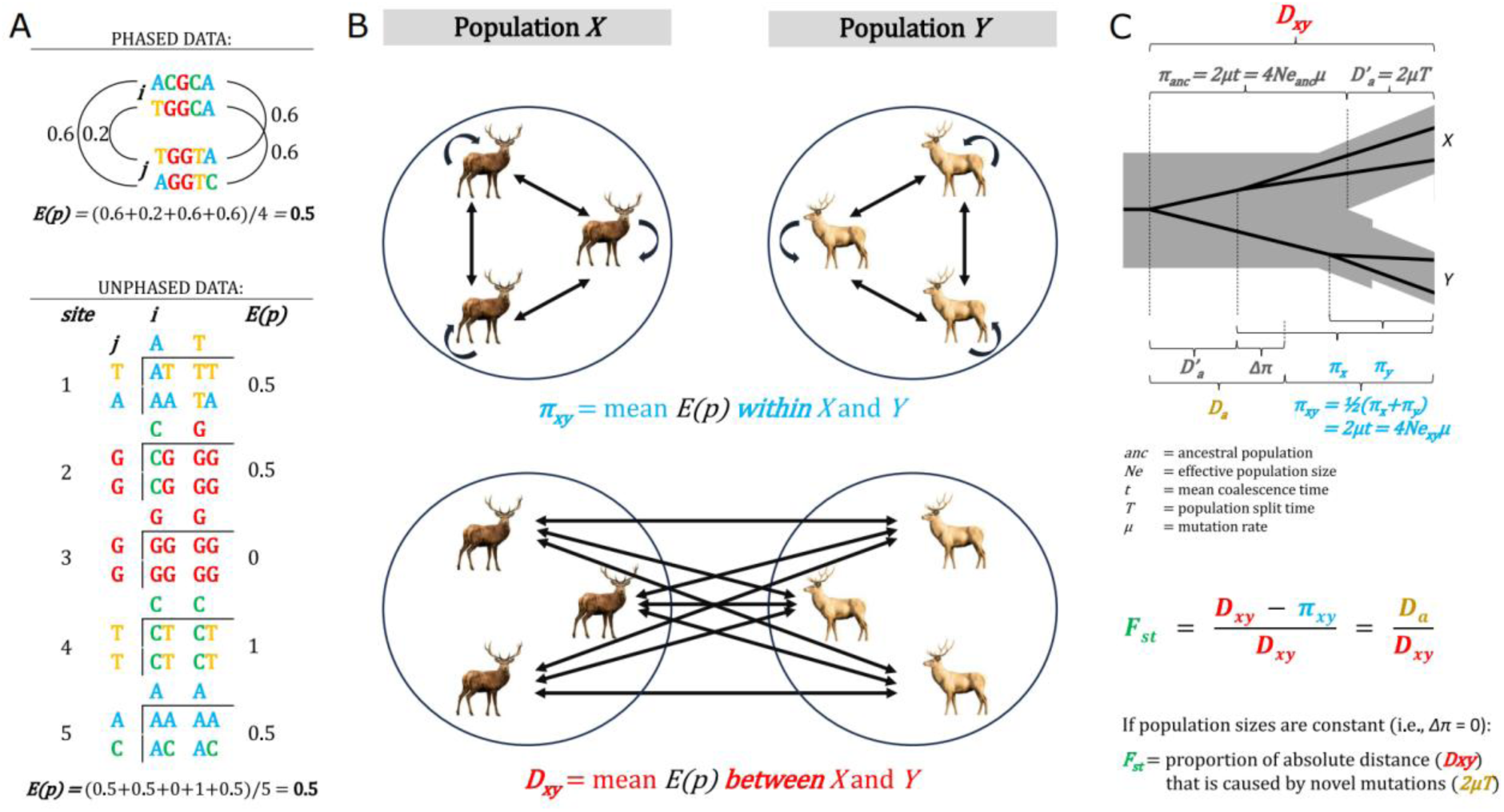
Genetic distance calculations. ***A***. Example calculation of expected sequence dissimilarity, *E(p)*, between a pair of diploid individuals, *i* and *j*, using either phased or unphased data. ***B***. Nucleotide diversity (*π_xy_*) denotes the mean *E(p)* for all possible pairs of individuals <u>within</u> populations *X* and *Y*, including self-comparisons (heterozygosity). Absolute/gross distance (*Dxy*) denotes the mean *E(p)* for all possible pairs of individuals <u>between</u> populations *X* and *Y*. ***C***. Assuming mutation rate constancy and the absence of parallel and back mutations (i.e., the infinite sites model), *π_xy_* and *Dxy* are proportional to mean coalescence times (*t*). When effective population sizes (*Ne*) vary, e.g. through reduced *Ne* in population *Y*, net nucleotide divergence (*Da*) is the sum of novel mutations (*2μT*) and the change in nucleotide diversity following a population split (*Δπ*). When *Ne* is constant (*Δπ* = 0), the fixation index (*Fst*) denotes the proportion of *Dxy* caused by novel mutations after the population split (*2μT*) rather than by ancestral nucleotide diversity (*4Ne_anc_μ*).

Measures of absolute and relative genetic distances are needed in combination because, on their own, they do not suffice for species delimitation. The reason is that both measures are confounded by effective population size (*Ne*), either before (*Dxy*) or after (*Fst*) a population split. High *Dxy*-values may indicate deep population split times (high *T*), but also recent divergence from a genetically diverse ancestral population (high *π_anc_*). Similarly, high *Fst*-values may indicate a deep population split (high *T*), or alternatively, signal a recent population bottleneck (low *π_xy_*) (Charlesworth 1998; Frankham et al. 2012; Hey and Pinho 2012). According to many species concepts, the first scenario (a deep population split) justifies species recognition, while the second does not.

Fortunately, a combination of high *Dxy-* and high *Fst*-estimates is theoretically more typical of deep population splits only, and hence a better diagnostic of the prolonged independent evolutionary histories needed to build up species-level differentiation. We tested this prediction by examining how well independent and combined *Dxy*- and *Fst*-thresholds can differentiate species pairs from conspecific pairs.

## RESULTS & DISCUSSION

### Taxonomic level versus hybrid fertility

We used two approaches to categorise lineage pairs as either species pairs or conspecific pairs. First, we followed current classifications of the Mammal Diversity Database (https://www.mammaldiversity.org/) (Burgin et al. 2025), distinguishing between three categories: intraspecific, interspecific, and intergeneric (Table S2). Given their recent split times, pairs of domesticated breeds and their wild relatives were considered conspecific, even if listed in the Mammal Diversity Database as separate species.

As expected from the confounding influence of population size differences, and consistent with previous studies (Hey and Pinho 2012; Lebedev et al. 2025), substantial overlap in *Dxy-* and *Fst*-estimates was observed between the category of interspecific lineage pairs and the other two categories (Fig. 2, S2A). Even so, a Kruskal-Wallis test and subsequent posthoc tests (i.e., pairwise Wilcoxon rank sum tests), suitable for non-parametric data, indicated that the *Dxy-* and *Fst*-estimates differ significantly among all categories, including intraspecific and interspecific comparisons (Fig. 2), with *p*-values below 10^−15^.

**Figure 2.**
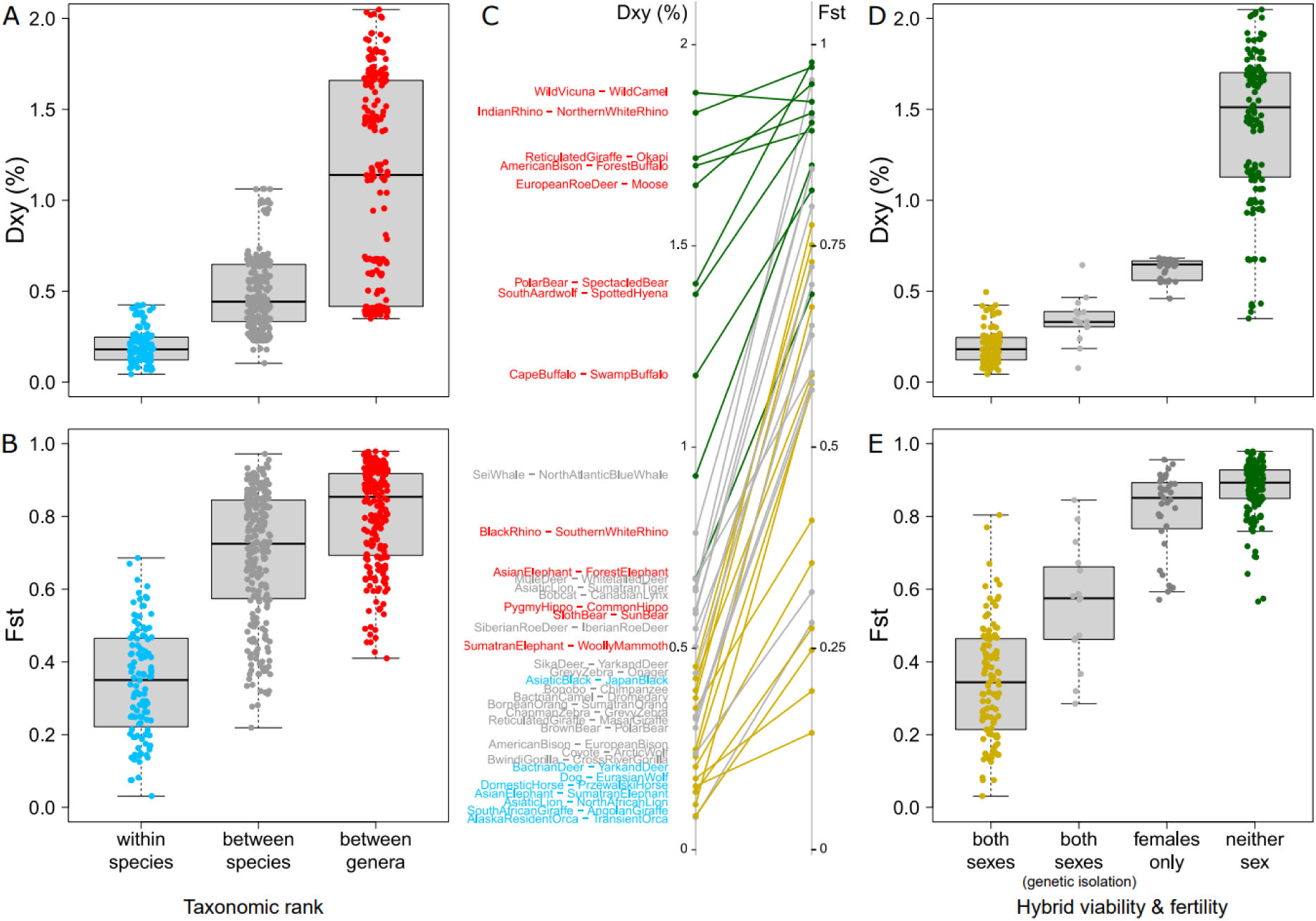
Genetic distances across taxonomic level and hybrid viability/fertility. **A**. Boxplots depicting absolute genetic distances (*Dxy*, in percentages) between three types of mammalian lineage pairs – intraspecific (lightblue), interspecific (grey) and intergeneric (red) – based on current taxonomic status in the Mammal Diversity Database. **B.** Same as A, but for relative genetic distances (Hudson *Fst* estimates). **C**. Line chart depicting genetic distances for an arbitrary selection of lineage pairs. Colour coding is the same as in A-B, and D-E. **D**-**E**. Same as A and B, but with lineage pairs categorised based on hybrid viability and fertility, with either both sexes fertile (gold, or light grey when evidence exists for genetic isolation), only females fertile (dark grey) or neither sex fertile or viable (green). Lineage pairs for which only female hybrids are fertile exhibit Haldane’s Rule. Lineage pairs with missing information are omitted, except for intraspecific lineage pairs, which were assigned to the first category.

As a second approach to categorise lineage pairs as either within-species pairs or species pairs, we assessed their capacity to interbreed and produce fertile offspring (Table S2). This criterion is inherently challenging to apply because reproductive compatibility does not naturally fall in discrete classes but rather spans a continuum of reduced hybrid viability and fertility. Moreover, available information is often indirect or circumstantial, making it difficult to discern whether reported exceptions prove or disprove a rule (Skidmore et al. 1999; Robinson et al. 2005).

Nevertheless, based on published literature (Benirschke 1967; Gray 1971; Yadav et al. 2019), we attempted to assign each lineage pair to one of three categories depending on the fertility of the F1 hybrids: (i) both sexes fertile with only slightly reduced fitness; (ii) only females fertile, consistent with Haldane’s Rule (Haldane 1922; Allen et al. 2020); or (iii) neither sex fertile or even viable (Table S2). For the first category, we also recognised a subcategory of lineage pairs for which evidence exists of genetic isolation despite overlapping geographical ranges. The lineage pairs in our meta-analysis in the second category (exhibiting Haldane’s Rule) comprised species pairs of big cats (*Panthera*), bovines (*Bos/Bison*), roe deer (*Capreolus*) and New World deer (*Odocoileus*) (Douglas et al. 2011; Yadav et al. 2019). *Dxy*- and *Fst*-estimates differed significantly between this category and the other two, with *p*-values below 10^−15^, despite substantial overlap (Fig. 2D-E).

While associated (Fisher exact test: *p*-value < 10^−4^), the two explanatory variables (i.e., taxonomic rank and hybrid fertility) are not identical. For the majority of lineage pairs classified by the Mammal Diversity Database as sister species, F1-hybrids are both viable and fertile regardless of the sex (Fig. 2C, S2B). In other words, for most lineage pairs that are separate species according to the scientific consensus, no hard physiological barriers are known that preclude hybridization (Mallet 2005). A few illustrative examples of such species pairs are sika deer and red deer (Biedrzycka et al. 2012), Grevy zebra and plains zebra (Cordingley et al. 2009; Ito et al. 2015), blue whale and fin whale (Pampoulie et al. 2020), chimpanzee and bonobo (Vervaecke and Elsacker 1992), gelada and common baboons (Jolly et al. 1997), and Bornean and Sumatran orangutan (Cocks 2007). These examples highlight that mammalian taxonomists do not adhere strictly to Dobzhansky’s original biological species concept, which defines species as lineages that are “*physiologically incapable of interbreeding*” (Ayala and Fitch 1997).

According to the original biological species concept, within-species pairs correspond to the category “*both sexes fertile*” while species pairs correspond to “*only females fertile*” or “*neither sex fertile*”. In contrast, the revised biological species concept recognises that even lineage pairs in the category *“both sexes fertile”* may represent distinct species (Mayr 1970; Mayr 1996). This also holds true for the genotypic cluster and genetic species concepts (Table 1), which allow for occasional gene flow events that are insufficient to homogenise gene pools (Baker and Bradley 2006; Mallet 2020; Lebedev et al. 2025). According to these species concepts, the maintenance of separate gene pools upon secondary contact (i.e., genetic isolation) arises not through physiological reproductive barriers, but rather through isolating mechanisms such as habitat selection, assortative mating or reduced hybrid fitness (Cocks 2007; Senn and Pemberton 2009; Bohling et al. 2016; Dioli 2020; Mallet and Mullen 2022; Markviriya et al. 2022; Miller et al. 2024; Pineda et al. 2024).

### Predicting taxonomic level

In order to evaluate the extent to which genetic distance thresholds can predict the taxonomic level of lineage pairs, we calculated the accuracy of a range of *Dxy*- and *Fst*-thresholds in distinguishing between the 111 intraspecific and 257 interspecific pairs in our dataset. Following standard conventions, we defined accuracy as the sum of the proportion of true positives (TP) and the proportion of true negatives (TN), which in this context corresponds to lineage pairs that were correctly identified as either interspecific or intraspecific.

The best performing *Dxy*-threshold of 0.225% scored an accuracy of 0.89, while the best performing *Fst*-thresholds, which ranged between 0.36 and 0.39, scored an accuracy of 0.83 (Fig. 3A). A combined *Dxy*- and *Fst*-threshold of 0.225% and 0.26 increased the accuracy to 0.92 (Fig 3A, S3A). More specifically, these combined thresholds allowed us to correctly classify 252 out of 257 (98%) interspecific pairs, compared to 88 out of 111 intraspecific pairs (79%). This indicates high sensitivity but only moderate specificity (Fig. 3A), with the model classifying as species pairs a fifth of lineage pairs widely considered to be conspecific.

**Figure 3.**
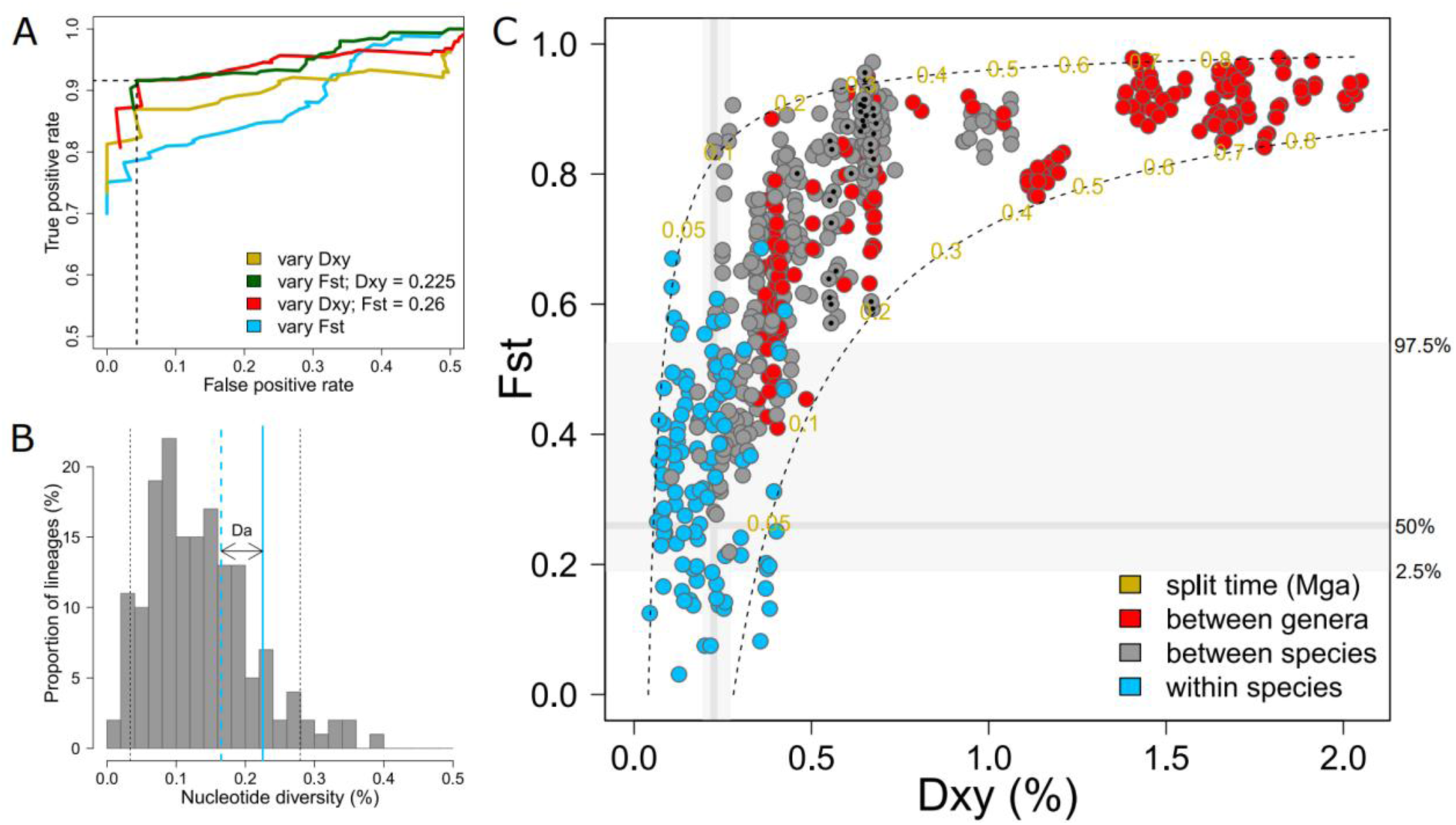
Species delimitation based on a combined *Dxy-* and *Fst*-threshold. **A**. Receiver operating characteristic (ROC) curve, depicting the accuracy of distinguishing between intraspecific and interspecific lineage pairs of large mammals based on genome-wide *Dxy* and Hudson *Fst* thresholds. The highest accuracy (dashed lines) is obtained with *Dxy* = 0.225% and *Fst* = 0.26. **B**. Histogram depicting sequence dissimilarity within lineages (nucleotide diversity, *π*, in percentages). The black dashed lines indicate the 5^th^ and 95^th^ percentiles of 0.03% and 0.28%. The solid and dashed blue lines denote *Dxy* = 0.225% and an ancestral nucleotide diversity (*π_anc_*) of 0.165%, respectively, resulting in *Fst* = 0.26 and a net divergence (*Da*) of 0.06%. **C**. Scatterplot depicting genome-wide *Dxy-*and *Fst*-estimates per lineage pair, with colour coding denoting current taxonomic rank. The dark grey lines indicate the here inferred *Dxy* and *Fst-*thresholds, with 95% bootstrap confidence intervals in light grey. Dashed lines denote predicted curves based on a mutation rate of 10^−8^ per site per generation, assuming constant effective population sizes (*Ne*) of 10,000 and 70,000 individuals, with the numbers denoting split time in millions of generations (Mga). Black dots mark lineage pairs that exhibit Haldane’s rule.

In order to evaluate sampling bias, we performed additional bootstrap analyses to determine a 95% confidence interval for the best performing *Dxy*- and *Fst*-threshold. We identified best performing *Dxy* and *Fst*-thresholds for a random selection of lineage pairs, by repeatedly sampling genera with replacement, a procedure that was repeated 10,000 times. We defined 95% confidence intervals as the 2.5% and 97.5% percentiles of the resulting *Dxy* and *Fst*-thresholds. The resulting confidence interval was relatively narrow for the *Dxy*-threshold [95% CI: 0.195 – 0.27] but much wider for the *Fst*-threshold [95% CI: 0.19 – 0.54] (Fig. S3B).

That genetic distance thresholds can predict the current taxonomic rank of large mammalian lineage pairs with more than 90% accuracy indicates that mammalian taxonomists have managed to maintain an admirable level of consistency in species demarcation across genera. Most species of large mammals were first recognised by modern science long before genetic data became available, largely by explicit or implicit adherence to the morphological species concept. This concept holds that members of a species share a particular set of distinct physical traits that distinguish them from other species (Cronquist 1978), but without specifying the degree of morphological differentiation indicative of species boundaries. Major inconsistencies across genera caused by subjective taxonomic judgment appear, nevertheless, to be rare.

The ability to predict taxonomic rank based on distance thresholds should not be considered support for the genetic species concept. It only demonstrates that two lineages can be classified as separate species when their absolute and relative genetic distances fall within the range characteristic of lineages widely considered to be distinct species. This differs from the operational criterion of the genetic species concept, as such lineage pairs are not necessarily genetically isolated.

### The *Dxy*-vs-*Fst* plane

The distribution of genetic distances across lineage pairs can be depicted by plotting *Fst*-estimates on the y-axis against *Dxy*-estimates on the x-axis (Fig. 3C). Assuming population size constancy, this plot depicts the mean sequence dissimilarity between lineages (x-axis), and the proportion of this dissimilarity that is caused by novel mutations after the population split (y-axis) (Fig. 1C, S4A-B).

Because *Fst*-values rapidly converge toward one while *Dxy*-estimates increase more or less proportionally for millions of generations, any ‘*Fst* vs *Dxy*’-plane is expected to reveal an asymptotic curve (Fig. 3C) (Duranton et al. 2018). Assuming population size constancy, a global mutation rate (*μ*) of 10^−8^ mutations per site per generation (Bergeron et al. 2023; Zhang et al. 2023), and adhering to Hudson’s *Fst* formula (*Fst* = *Dxy* – *π_xy_*)/*Dxy*) (Hudson et al. 1992), the left-hand and right-hand boundaries are roughly predicted by *Ne* = 10000 and *Ne* = 70000 (Fig. 3C, S4C). Given *π* = 4*Neμ*, these *Ne*-values translate to π-values of 0.04% and 0.28%, which correspond to the 5^th^ and 95^th^ percentiles of observed *π*-values (0.03% and 0.28%) (Fig. 3B).

Using the inferred *Fst*- and *Dxy*-thresholds, we can divide the ‘*Fst* vs *Dxy*’-plane into four sections (Fig. 3C, S4A-B). In clockwise direction, and starting from the top-left, these sections contain:

1. lineage pairs with low *Dxy* but high *Fst* (top-left), suggestive of *a recent population split with a subsequent population bottleneck*, such as Asiatic and North African lions (*Dxy* = 0.11%, *Fst* = 0.58);
2. lineage pairs with high *Dxy* and high *Fst* (top-right), suggestive of *a deep population split*, such as chimpanzees and bonobos (*Dxy* = 0.40%, *Fst* = 0.78);
3. lineage pairs with high *Dxy* but low *Fst*, (bottom-right), suggestive of *a recent population split from an ancestral population with high genetic diversity*, such as Iberian and Caspian red deer (*Dxy* = 0.26%, *Fst* = 0.21);
4. lineage pairs with low *Dxy* and low *Fst* (bottom-left), suggestive of a *recent population split*, such as Arctic and Eurasian grey wolves (*Dxy* = 0.18%, *Fst* = 0.18).

The near-absence of interspecific pairs in the top-left section of the *‘Fst* vs *Dxy*’-plane (Fig. 3C), fails to support the relevance of bottleneck speciation in large-bodied mammals (Nei 2005; Mallet 2020). Thus, incipient mammalian species appear to arise through accumulating novel mutations rather than through novel allele combinations caused by random genetic drift (i.e., genetic revolutions). Most intraspecific pairs in the top-right section of the ‘*Fst*-vs *Dxy*’ have genetic distances just above the here inferred thresholds, and hence are not strong examples of cryptic speciation, where lack of morphological differentiation belies deep population split times (Baker and Bradley 2006) (Fig. 3C).

Rather than revealing cryptic species, the increased reliance of species delimitation on genetic data appears to have lowered the bar. Many lineage pairs of recently recognised species – such as western and eastern gorilla (*Dxy* = 0.23%, *Fst* = 0.50), Sumatran and Tapanuli orangutan (*Dxy* = 0.27%, *Fst* = 0.22), Tianshan wapiti and Yarkand deer (*Dxy* = 0.24%, *Fst* = 0.32), onager and kiang (*Dxy* = 0.27%, *Fst* = 0.5), reticulated and Nubian giraffe (*Dxy* = 0.24%, *Fst* = 0.39), and Tamanend’s and common bottlenose dolphin (*Dxy* = 0.18%, *Fst* = 0.36) – have at least one type of genetic distance just above, at or below the here inferred thresholds. Two exceptions are the lineage pairs of common and Indo-Pacific bottlenose dolphins (*Dxy* = 0.34%, *Fst* = 0.58) (Wang et al. 1999) and forest and savanna elephants (*Dxy* = 0.34%, *Fst* = 0.58) (Roca et al. 2001).

### Speciation duration

The *Dxy*-threshold of 0.225% does not exceed observed levels of nucleotide diversity within the study populations, which although on average approximately 0.12%, reach values as high as 0.4% (Fig. 3B). Paired with the deduced *Fst*-threshold of 0.26, and assuming *μ* = 10^−8^, the *Dxy*-threshold of 0.225% corresponds to a demographic scenario in which two sister populations with a constant *Ne* split 30,000 generations ago from an ancestral population with a nucleotide diversity of 0.165%, implying a net divergence (*Da*) of 0.06% (Fig. 3B). If assuming population size constancy, a *Fst*-value of 0.26 corresponds to a population split time of 0.35 coalescent units (M. J. de Jong et al. 2024). Under this scenario, over 50% of haplotypes are expected to coalesce more recently than the population split, corresponding to a genealogical divergence index (*gdi*) of 0.3 (Table S3) (Jackson et al. 2017; Rannala and Yang 2020).

As mentioned above, current species boundaries reflect scientific consensus, not necessarily the ability or inability of lineages to interbreed. Indeed, lineage pairs in our dataset that exhibit Haldane’s Rule (the strictest definition of reproductive isolation) exceed thresholds of *Dxy* = 0.004 and *Fst* = 0.55, well above the deduced species boundaries (Fig. 3C). If we adhere to the revised biological species concept (Table 1), which overlaps with the genetic species concept, the correct threshold for species delineation should be inferred from lineages that exhibit genetic isolation (Fig. 2D-E) (Bradley and Baker 2001; Baker and Bradley 2006; Mallet 2020; Lebedev et al. 2025).

For lineages diverging in allopatry, the criterion of genetic isolation through assortative mating can only be evaluated for lineage pairs that are partially sympatric, either naturally or by human-caused translocations. Low percentages of early-generation hybrids have been found in contact zones of wolves and coyotes (<4%) (Bohling et al. 2016), polar bears and brown bears (<1%) (Miller et al. 2024), bobcat and Canadian lynx (<0.25%) (Koen et al. 2014), mountain zebra and plains zebra (0%) (Hrabar and Kerley 2013), red deer and sika deer (<2%) (Senn and Pemberton 2009), red deer and wapiti (<0.25%) (Smith et al. 2018), black and blue wildebeest (<3%) (Grobler et al. 2018), and Masai giraffes and reticulated giraffes (0%) (Coimbra et al. 2023).

Such a near-absence of hybrids either indicates reduced fitness of hybrids, as in the case of camels and dromedaries (Dioli 2020), or alternatively assortative mating – as in the case of plains and Grevy zebras (Cordingley et al. 2009), swamp-type and river-type buffalos (Pineda et al. 2024), transient and resident killer whales (Hoelzel and Dover 1991), guanaco and vicuna (Lucherini 1996), or common, Tamanend’s and Indo-Pacific bottlenose dolphins (Gridley et al. 2018; Costa et al. 2022).

This number of case studies is unfortunately too limited to draw strongly supported conclusions. Based on the available data included in our study, genetic isolation can occur among lineage pairs with a net divergence (*Da*) as low as 0.03%, as observed for transient and resident killer whales (Hoelzel and Dover 1991; Morin et al. 2024), but can also be absent in lineage pairs with a *Da* as high as 0.3%, as observed for forest elephants and savanna elephants (Roca et al. 2001; Mondol et al. 2015; Bonnald et al. 2024). Overall, however, we encountered very few examples of freely hybridising lineage pairs with a *Da* above 0.2% (Fig. S5). Assuming *μ* = 10^−8^ (Bergeron et al. 2023; Zhang et al. 2023), this *Da*-value of 0.2% suggests a rough upper boundary for a speciation duration of approximately 100,000 generations, which translates to 0.5–1.0 My given a generation time of 5 or 10 years (Lister 2004; Hedges et al. 2015).

### The grey zone of speciation

The grey zone of speciation denotes the range where limited gene flow persists such that taxonomic decisions are inherently controversial (Roux et al. 2016). If we may equate the grey zone of speciation to the transitional phase of genetic isolation (which separates unlimited genetic exchange from hard physiological reproductive barriers, as demonstrated by Haldane’s Rule), then our study findings suggest that the grey zone of speciation in large mammals is associated with a net divergence of up to 0.2%.

Our findings contrast with earlier claims, which suggest grey zones of speciation which are both deeper in time and wider. Roux et al (2016) concluded, by testing for ongoing gene flow using demographic modelling, that “the intermediate ‘grey zone’ of speciation, in which taxonomy is often controversial, spans from 0.5% to 2% of net synonymous divergence”. Galtier (2019) defined the grey zone as extending from 0.15% (genetic distances within humans) to 1.2% (chimp–human divergence). Jackson et al. (2017) reported a genealogical divergence index range of 0.2–0.7, which for deep population splits roughly corresponds to a similar range of *Fst*-values (Table S3).

The upper limit of 2% suggested by Roux et al. (2016) exceeds, according to our calculations, net distances between lineage pairs as divergent as giraffes and okapis, one-horned and two-horned rhinos, roe deer and moose, or cattle and cape buffalos, which are all incapable of producing viable let alone fertile offspring (Owiny et al. 2009; Prins et al. 2023). The suggested lower limit of 0.5% still exceeds net distances between lineage pairs as divergent as Asian elephants and mammoths, lions and leopards, zebras and wild asses, dromedaries and camels, or brown bears and American black bears. Species-level boundaries are not controversial for any of these lineage pairs, nor does evidence exist for ongoing gene flow where the geographical ranges of extant species overlap.

The discrepancy in study findings cannot be attributed to differences in underlying data types (i.e., whole genomes versus synonymous sites), as codon usage bias causes only slight deviations from neutral expectations (Li et al. 2015). Instead, the major cause for the discrepancy is likely the limited overlap between the two sets of study species. Whereas our study considers data of large mammals only, Roux et al. (2016) included data of 61 lineage pairs from across the animal kingdom, of which only twenty pairs were vertebrates, among them five mammals, and only one large mammal (i.e., western gorilla versus eastern gorilla). The discrepancy in study findings might therefore imply that the grey zone of speciation is best inferred in a taxon-specific manner (Dufresnes et al. 2023).

Lebedev et al. (2025), who focused on mammals specifically, reported a *Dxy*-threshold for intraspecific versus interspecific lineage pairs of mammals of approximately 0.15%, which is below the *Dxy*-threshold of 0.225% inferred here. For overlapping species pairs, *Dxy*-estimates of Lebedev et al. (2025) are approximately half the values reported here (Fig. S6). This discrepancy likely reflects different selection pressures, as Lebedev et al. (2025) studied the exome, from which many new mutations are removed by purifying selection.

### Dealing with the speciation continuum in time and space

Even when the grey zone of speciation is narrower than previously assumed, drawing a sharp boundary in a continuum of evolutionary divergence will never be fully satisfactory and will invariably yield controversial borderline cases (Coates et al. 2018; Stankowski and Ravinet 2021). Species delimitation becomes particularly challenging when the continuum of divergence has a spatial as well as temporal dimension (Mayr 1996; Coates et al. 2018; Stankowski and Ravinet 2021; Dufresnes et al. 2023; Stankowski et al. 2024).

Isolation-by-distance trends, a hallmark of reticulated evolution, highlight these difficulties. For instance, the genetic distance between Amur leopards (*Panthera pardus orientalis*) and African leopards (*P. p. pardus*) is well above the here inferred thresholds (*Dxy* = 0.31%, *Fst* = 0.53), but these lineages are connected by geographically and genetically intermediate populations (Mochales-Riaño et al. 2023). Similarly, while genetic distance between the cape buffalo (*Syncerus caffer caffer*) and forest buffalo (*S. c. nanus*) is indicative of species-level differentiation (i.e., *Dxy* = 0.39%, *Fst* = 0.31), the distances with intermediate lineages of the Nile buffalo (*S. c. brachyceros*) and Sudan buffalo (*S. c. aequinoctialis*) are not.

A potential pragmatic solution is an exemption rule. Lineages may be considered and treated as different species if their level of genetic differentiation is above the here observed thresholds, unless extant intermediate lineages exist with result in genetic distances below the specified thresholds. However, according to our calculations this rule would have profound implications, such as that grey wolves (*Canis lupus*) and coyotes (*C. latrans*) ought to be considered intraspecific varieties of a single polytopic species. While the genetic distances between unadmixed populations of these two lineages meet the here inferred thresholds for species delimitation (*Dxy* = 0.24% and *Fst* = 0.31), those between admixed varieties, such as the Great Lakes wolf, eastern wolf, red wolf and eastern coyote, do not.

### Reproducibility of distance estimates

A potential weakness of species delimitation based on genetic distance estimates is that these estimates are sensitive to the bioinformatic pipelines used for mapping and genotype calling. For comparison, while our calculations suggest *Dxy*-estimates of 0.10%, 0.23%, 0.26%, 0.42%, 0.6% and 0.67% for the lineage pairs of bighorn sheep vs thinhorn sheep, western vs eastern gorilla, taurine vs zebu cattle, vicuna vs lama, bison vs cattle, and African vs Asian elephants, previously published estimates are 0.09%, 0.20%, 0.26%, 0.47%, ∼0.65% and 0.84-0.9%, respectively (Roux et al. 2016; Palkopoulou et al. 2018; Wang et al. 2018; Santos et al. 2021; Zefu Wang et al. 2025).

Sensitivity analyses suggest that the obtained sequence dissimilarity estimates are largely robust to potential confounders such as mean sample depth, data thinning parameters and the reference genome, with the precision of the estimates being approximately 0.02% (Fig. S7). In contrast, filters on site quality and the proportions of missing data per site were found to introduce substantial bias. These filters were therefore omitted from our analyses (Fig. S7) and replaced with a site depth filter.

Given the inherent differences between mapping and genotype calling approaches, we acknowledge that the inferred species-level thresholds should be considered rough estimates at best. Including a set of benchmark lineage pairs that fall in the grey zone may help to standardise outcomes across research groups, regardless of the mapping and genotype-calling pipelines employed.

### Potential misclassifications

A key rationale of our study is that high relative and absolute genetic distances are together characteristic of a deep population split. This assumption can be violated by two evolutionary scenarios. When sister lineages split from an ancestral population with exceptionally low genetic diversity, *Dxy-*estimates will be low despite deep divergence times, resulting in false negatives. At the opposite extreme, bottlenecked sister lineages that split from an ancestral population with exceptionally high genetic diversity may have high *Dxy-* and high *Fst*-estimates despite shallow divergence times, resulting in false positives. Caution should therefore be exercised when drawing conclusions regarding outlier pairs.

With the risk of circular reasoning, interspecific lineage pairs with low *Dxy-* and/or low *Fst*-estimates raise suspicion of taxonomic inflation (Baker and Bradley 2006). Among the lineage pairs included in our meta-analysis, this applies in particular to thinhorn sheep and bighorn sheep (*O. dalli* vs *O. canadensis*, *Dxy* = 0.10%, *Fst* = 0.33) and to pairs of domesticated animals and their wild ancestors. For instance, and consistent with their shallow divergence times, the absolute genetic distance between Bactrian camels and wild camels (*Dxy* = 0.12%), lamas and guanacos (*Dxy* = 0.18%), domestic horses and Przewalski horses (*Dxy* = 0.16%), and dogs and grey wolves (*Dxy* = 0.18%) are all well below the here inferred thresholds.

Conversely, intraspecific lineage pairs with high *Dxy-* as well as high *Fst*-estimates raise suspicion of taxonomic deflation. Potential candidates for elevation from subspecies to the species level are the Japanese and continental Asiatic black bear (*Ursus thibetanus japonicus* vs *Ursus thibetanus sspp.*, *Dxy* = 0.43%, *Fst* = 0.59) (Zou et al. 2022), the southern and eastern aardwolf (*Proteles cristatus cristatus* vs *P. c. septentrionalis*, *Dxy* = 0.36%, *Fst* = 0.69) (Allio et al. 2021), and European and Asian wild boar (*Dxy* > 0.4%, *Fst* > 0.45). However, for the latter species pair, admixture at a contact zone in central Asia refutes genetic isolation (Zishuai Wang et al. 2025).

Our analyses furthermore support suggestions to raise swamp-type water buffalo to the species level (priority name: *B. carabanesis*), separate from wild and domesticated river-type buffalo (priority name: *B. bubalis*). High genetic distances between the two lineages (*Dxy* = 0.33%, *Fst* = 0.46) indicate that the swamp buffalo was domesticated from a now extinct wild ancestor, which split from wild water buffalo long before domestication commenced (Curaudeau et al. 2021; Cailipan et al. 2023; Pineda et al. 2024; Si et al. 2025). Similarly, zebu cattle differ markedly from taurine cattle and their aurochs ancestors (*Dxy* = 0.26%, *Fst* = 0.52).

### Temporal banding

Shifting the focus from species boundaries to genus boundaries, relatively low absolute genetic distances were observed among dolphin genera within the subfamily Delphininae (Dxy < 0.4%) as well as among the genera of brown hyenas and striped hyenas (*Parahyaena* vs *Hyaena*, *Dxy* < 0.4%), woolly mammoths and Asian elephants (*Mammuthus* vs *Elephas*, *Dxy* < 0.52%), straight-tusked elephants and African elephants (*Palaeoloxodon* vs *Loxodonta*, *Dxy* < 0.5%), bisons and remaining *Bos* (*Bison* vs *Bos*, *Dxy* < 0.56%), common hippos and pygmy hippos (*Hippopotamus* vs *Choeropsis*, *Dxy* < 0.6%) and the bear genera of *Ursus*, *Melursus* and *Helarctos* (*Dxy* < 0.7%). Conversely, unusual high genetic distances were observed within the genus *Equus* (*Dxy* > 0.95%), potentially supporting the recognition of a genus of asses and zebras separate from that of caballine horses (S4A-B).

Merging and/or splitting of the aforementioned genera would improve temporal banding (Van Gelder 1977), placing the boundary between intra- and intergeneric lineage pairs at *Dxy* ≈ 0.7-0.9%. Assuming *μ* = 10^−8^, a *Dxy*-value of 0.9% corresponds to approximately 450,000 generations (≈ 4.5 My, given a generation time of 10 years), which is broadly consistent with previous estimates of generic boundaries (Groves 2001).

The question is, however, to what extent such revisions would create a biologically meaningful boundary (Van Gelder 1977). A common *Dxy*-threshold for genus boundaries may, in fact, decrease the correspondence between genus boundaries and physiological reproductive isolation (Fig. 2D). For instance, strongly reduced hybrid fertility has been reported for crosses between the relatively closely related zebras and wild asses (*Hippotrigris* vs *Asinus*, *Dxy* < 0.5%) (Gray 1971). On the opposite end of the divergence spectrum, repeated attempts to crossbreed the highly divergent new world and old world camelids (*Lama* vs *Camelus*, Dxy > 1.9%) have occasionally produced viable progeny (Skidmore et al. 1999). These examples underscore that the accumulation of mutations that underlie reproductive isolation is a stochastic rather than deterministic process, making genetic distance an imperfect predictor of reproductive isolation.

### Concluding remarks

Species concepts are akin to axioms: unproven statements accepted as true, that serve as a basis from which one can derive other statements, such as species delimitation rules. If a species is considered a population with one or more diagnostic traits, as stated by the phylogenetic species concept (Baum 1992; Groves 2013), it follows logically that young monophyletic lineages such as Sumatran tigers and Corsican red deer should be elevated to the species rank (Cracraft et al. 1998; Pitra et al. 2004). If instead a species is defined as a lineage with an independent evolutionary history of at minimum several 10^5^ years, it follows logically that their subspecies rank ought to be maintained.

In the debate surrounding the phylogenetic species concept (Baum 1992; Isaac et al. 2004; Meiri and Mace 2007; Frankham et al. 2012; Groves 2013; Heller et al. 2013; Zachos et al. 2013; Zachos and Lovari 2013; Zachos 2015), it is frequently argued that the value of a species concept lies in its diagnosability: its ability to generate testable and therefore falsifiable species boundaries. Species concepts that employ genetic distance thresholds partially fulfil this requirement. They provide objectivity in terms of a common set of rules that enable independent researchers to draw repeatable conclusions (Hey and Pinho 2012). Here, we demonstrate that distance thresholds can, moreover, be defined in a non-arbitrary manner, and distinguish, with reasonable accuracy, two different types of real natural entities (Hey and Pinho 2012): lineage pairs which are genetically isolated, and lineage pairs which are not. In addition, we provide evidence that the grey zone can be narrowed when distance thresholds are defined in a taxon-specific manner, leaving less room for ambiguity. Nevertheless, we also acknowledge that the suggested upper boundary of *Da* = 0.2% for genetic isolation retains a considerable margin of uncertainty, which may faithfully reflect the gradual and stochastic nature of speciation, but does not aid effective binary decision-making.

One might argue that reducing species delimitation to measures of genetic distance is overly reductionistic and risks oversimplifying natural history. However, this perceived limitation is in fact a strength: genomic data provides a standardised basis for straight-forward and consistent decision-making using a measure that is proportional to split time and that, aside from variation in mutation rates, does not rely on lineage-specific idiosyncrasies (Groves 2001).

In terms of practical implications, the here suggested *Dxy-* and *Fst*-thresholds serve to delimit species only, and not to identify conservation units, aka evolutionary significant units (Taylor et al. 2017; Coates et al. 2018; Dufresnes et al. 2023; Hoelzel 2023). Conservation agencies increasingly aim to conserve species as well as the breadth of intraspecific variation, as illustrated by CITES split-listings, IUCN subspecies risk assessments, and EAZA subspecies breeding programs. A conservation unit is therefore an umbrella term which includes species but also subspecific lineages. By recognising this distinction between species and conservation units, species delimitation is a matter of purely fundamental science, free from personal, political or economic motivations (Cracraft et al. 1998; Hey et al. 2003; Chaitra et al. 2004; Isaac et al. 2004; Padial and De la Riva 2021; Dufresnes et al. 2023).

## ACKNOWLEDGEMENTS

We thank Frank Zachos for providing comments on an earlier draft of the manuscript. We thank Tunca Deniz Yazici, Verena Hartmann, Marcel Nebenfuehr, Shahriar Nur, David Prochotta, and Saurav Sharma for help with the data analyses.

## DATA AND CODE AVAILABILITY

No new sequencing data has been generated for this study. Scripts for mapping and genotype calling: https://github.com/mennodejong1986/WGS_data_analyses. Scripts for genetic distance calculations: https://github.com/mennodejong1986/DIST.

## AUTHOR CONTRIBUTIONS

**MJ**: conceptualization (lead), data analysis (equal), interpretation of study finding (equal), manuscript writing (lead). **CW**: data analysis (equal), interpretation of study findings (equal), manuscript writing (supporting). **MM**, **BF**, **MW**: data analysis (supporting), review and editing (supporting). **AJ**: interpretation of study findings (supporting), review and editing (supporting), funding acquisition (lead).

## METHODS

### Read mapping and genotype calling

We obtained publicly available short-read sequencing data as well as reference genomes either from Dryad or the short read archive (SRA) repository of NCBI/ENA, targeting at minimum 10x depth per individual (Table S1A-B). We strived to select multiple samples per lineage (i.e., per panmictic population).

Reads were filtered using the software fastp (Chen et al. 2018) using default options and subsequently mapped using the software BWA mem (Li and Durbin 2009) to a reference genome of a closely related species. Samtools (Li et al. 2009) was used to remove reads with a mapping quality below 20 and/or alignment scores below 100 (assuming a read length of 150bp), as well as reads that mapped discordantly or to multiple locations in the genome, the latter by applying a minimum mapping quality threshold of 20. Read duplicates were removed using the software Picard (DePristo et al. 2011).

Genotype likelihoods and calls were generated using the bcftools mpileup and call pipeline (Li 2011). Indels were normalised and realigned using ‘bcftools norm’. When calling genotypes from genotype likelihoods (bcftools call), we used the ‘group-samples’ option to assign each individual to its unique group (i.e., we disabled the option of influencing genotype calls based on information from other samples). As such, our approach does not make a priori assumptions about population assignment, meaning the results are reproducible regardless of the sample set.

The bcftools filter command was used to mask sites with a read depth below five, after establishing experimentally that this provided a balance between disposal of useful data and incorrect heterozygosity estimation (de Jong et al. 2023). Sites were removed based on a global depth, after visual examining of the depth distribution across sites (Fig. S5). To avoid biases, we did not filter on site quality (Fig. S6) nor on proportion of missing data per site (Fig. S7). We validated that allelic ratios for heterozygous sites were approximately 0.5 (Fig. S8).

### Genetic distance calculations

Expected sequence dissimilarity estimates, *E(p)* (Fig. 1A), for each pair of individuals were estimated using a custom-built Unix script (Jong and Janke 2025), using the following rules: *E(p)* = 0 for AA:AA; *E(p)* = 0.5 for AT:TT and AT:AT; *E(p)* = 0.75 for AT:CT; and *E(p)* = 1 for AA:TT and AT:CG (Mountain and Ramakrishnan 2005). Sites with missing data for one or both individuals involved in the pairwise comparison were excluded. Genetic distances for all pairs of individuals were superimposed on UPGMA dendrograms, generated using the ‘upgma’ function of the R package ‘phangorn’ (Schliep 2011), in order to evaluate adherence to the theoretical expectation of equidistance in the absence of gene flow (Fig S9).

For each population, nucleotide diversity (*π*) values were derived from the *E(p)*-estimates, namely as the mean of all possible pairwise sample comparisons <u>within</u> a population (Fig. 1B). Similarly, population pairwise *Dxy* values were derived as the mean of all possible pairwise sample comparisons <u>between</u> two populations (Fig. 1B). Relative distances (i.e., normalised net divergence) between populations were estimated using the formula: *F_ST_ = (Dxy–π_xy_)/Dxy* (Hudson et al. 1992) (Fig. 1B). These calculations were performed using the ‘runDIST’ function of the R package SambaR (de Jong et al. 2021; M. J. de Jong et al. 2024; Jong and Janke 2025).

We calculated distances across all autosomes, without discriminating between functional and non-functional regions. While natural selection on functional regions may cause deviations from neutral expectations, this confounding effect is small given the low functional content of the genomes of our study species. Linked selection, in particular background selection, does affect a much larger part of the genome, but its effects are equivalent to those of genetic drift and limited to lowering effective population sizes (*Ne*) (Birky and Walsh 1988; Charlesworth 1998; Menno J. de Jong et al. 2024).

### Statistical analyses

Taxonomic rank was assigned to lineage pairs following classifications of the Mammal Diversity Database (https://www.mammaldiversity.org/) (Burgin et al. 2025). In order to find the optimal genetic distance thresholds for predicting the taxonomic level of lineage pairs, we calculated the accuracy of a range of *Dxy*- and *Fst*-thresholds in distinguishing between intraspecific and interspecific pairs. Following standard conventions, we defined accuracy as the sum of the proportion of true positives (TP) and the proportion of true negatives (TN), which in this context corresponds to lineage pairs that were correctly identified as either interspecific or intraspecific. Bootstrap confidence intervals were estimated by rerunning this procedure 10,000 times using different input datasets. The bootstrapped input datasets were generated by sampling with replacement from the list of input genera, and by excluding lineage belonging to genera not drawn. All analyses were performed using custom-built R scripts (see captions of figures 3A and 3B).

