## Supplementary tables and figures for "Species delimitation using genome-wide estimates of *Dxy* and *Fst*"

**Tables S1A.** List of lineages included in the analyses.

| Common_name | genus | species | subspecies | latin | dataset |
|---|---|---|---|---|---|
| Hartebeest | Alcelaphus | buselaphus | NA | Alcelaphus_buselaphus | gnous |
| Moose | Alces | Alces | NA | Alces_Alces | capreolini |
| SeiWhale | Balaenoptera | borealis | NA | Balaenoptera_borealis | bluewhales |
| BlueWhale | Balaenoptera | musculus | NA | Balaenoptera_musculus | finwhales |
| NorthAtlanticBlueWhale | Balaenoptera | musculus | musculus | Balaenoptera_musculus | bluewhales |
| PacificBlueWhale | Balaenoptera | musculus | sulfureus | Balaenoptera_musculus | bluewhales |
| PygmyBlueWhale | Balaenoptera | musculus | brevicauda | Balaenoptera_musculus | bluewhales |
| FinWhale | Balaenoptera | physalus | NA | Balaenoptera_physalus | finwhales |
| AmericanBison | Bos | bison | NA | Bos_bison | bovini |
| EuropeanBison | Bos | bonasus | NA | Bos_bonasus | bovini |
| Gaur | Bos | gaurus | NA | Bos_gaurus | bovini |
| Yak | Bos | grunniens | NA | Bos_grunniens | bovini |
| Zebu | Bos | indicus | NA | Bos_indicus | bovini |
| Cattle | Bos | taurus | NA | Bos_taurus | bovini |
| RiverBuffalo | Bubalus | bubalis | NA | Bubalus_bubalis | bovini |
| LowlandAnoa | Bubalus | depressicornis | NA | Bubalus_depressicornis | bovini |
| SwampBuffalo | Bubalus | kerabau | NA | Bubalus_kerabau | bovini |
| BactrianCamel | Camelus | bactrianus | NA | Camelus_bactrianus | camelids |
| Dromedary | Camelus | dromedarius | NA | Camelus_dromedarius | camelids |
| WildCamel | Camelus | ferus | NA | Camelus_ferus | camelids |
| GoldenJackal | Canis | aureus | NA | Canis_aureus | wolves |
| Coyote | Canis | latrans | NA | Canis_latrans | wolves |
| EasternCoyote | Canis | latrans | var | Canis_latrans | wolves |
| AfricanWolf | Canis | lupaster | NA | Canis_lupaster | wolves |
| ArcticWolf | Canis | lupus | arctos | Canis_lupus | wolves |
| Dog | Canis | lupus | familiaris | Canis_lupus | wolves |
| EurasianWolf | Canis | lupus | lupus | Canis_lupus | wolves |
| EuropeanRoeDeer | Capreolus | capreolus | NA | Capreolus_capreolus | capreolini |
| IberianRoeDeer | Capreolus | capreolus | NA | Capreolus_capreolus | capreolini |
| SiberianRoeDeer | Capreolus | pygargus | pygargus | Capreolus_pygargus | capreolini |
| TianshanRoeDeer | Capreolus | pygargus | tianschanicus | Capreolus_pygargus | capreolini |
| NorthernWhiteRhino | Ceratotherium | simum | cottoni | Ceratotherium_simum | rhinos |
| SouthernWhiteRhino | Ceratotherium | simum | simum | Ceratotherium_simum | rhinos |
| ThoroldsDeer | Cervus | albirostris | NA | Cervus_albirostris | cervus |
| KansuRedDeer | Cervus | canadensis | kansuensis | Cervus_canadensis | cervus |
| NorthAmericanWapiti | Cervus | canadensis | manitobensis | Cervus_canadensis | cervus |
| TianshanWapiti | Cervus | canadensis | songaricus | Cervus_canadensis | cervus |
| CaspianRedDeer | Cervus | elaphus | maral | Cervus_elaphus | cervus |
| CentralEuropeanRedDeer | Cervus | elaphus | hippelaphus | Cervus_elaphus | cervus |
| IberianRedDeer | Cervus | elaphus | hispanicus | Cervus_elaphus | cervus |
| BactrianDeer | Cervus | hanglu | bactrianus | Cervus_hanglu | cervus |
| YarkandDeer | Cervus | hanglu | yarkandensis | Cervus_hanglu | cervus |
| SikaDeer | Cervus | nippon | NA | Cervus_nippon | cervus |
| PygmyHippo | Choeropsis | liberiensis | NA | Choeropsis_liberiensis | hippos |
| BlackWildebeest | Connochaetes | gnou | NA | Connochaetes_gnou | gnous |
| EastBlueWildebeest | Connochaetes | taurinus | albojubatus | Connochaetes_taurinus | gnous |
| SouthBlueWildebeest | Connochaetes | taurinus | taurinus | Connochaetes_taurinus | gnous |
| WestBlueWildebeest | Connochaetes | taurinus | mattosi | Connochaetes_taurinus | gnous |
| SpottedHyena | Crocuta | crocuta | NA | Crocuta_crocuta | hyenas |
| CommonDolphin | Delphinus | delphis | NA | Delphinus_delphis | dolphins |
| SumatranRhino | Dicerorhinus | sumatrensis | NA | Dicerorhinus_sumatrensis | rhinos |
| BlackRhino | Diceros | bicornis | NA | Diceros_bicornis | rhinos |
| AsianElephant | Elephas | maximus | NA | Elephas_maximus | elephants |
| BorneanElephant | Elephas | maximus | borneesis | Elephas_maximus | elephants |
| SriLankanElephant | Elephas | maximus | maximus | Elephas_maximus | elephants |

| | | | | | |
|---|---|---|---|---|---|
| SumatranElephant | Elephas | maximus | sumatranus | Elephas_maximus | elephants |
| AfricanWildass | Equus | africanus | NA | Equus_africanus | equus |
| DomesticDonkey | Equus | africanus | asinus | Equus_africanus | equus |
| DomesticHorse | Equus | ferus | caballus | Equus_ferus | equus |
| PrzewalskiHorse | Equus | ferus | przewalskii | Equus_ferus | equus |
| GrevyZebra | Equus | grevyi | NA | Equus_grevyi | equus |
| Kiang | Equus | hemionus | kiang | Equus_hemionus | equus |
| Onager | Equus | hemionus | onager | Equus_hemionus | equus |
| ChapmanZebra | Equus | quagga | chapmani | Equus_quagga | equus |
| KordofanGiraffe | Giraffa | camelopardalis | antiquorum | Giraffa_camelopardalis | giraffes |
| NubianGiraffe | Giraffa | camelopardalis | camelopardalis | Giraffa_camelopardalis | giraffes |
| WestAfricanGiraffe | Giraffa | camelopardalis | peralta | Giraffa_camelopardalis | giraffes |
| AngolanGiraffe | Giraffa | giraffa | angolensis | Giraffa_giraffa | giraffes |
| SouthAfricanGiraffe | Giraffa | giraffa | giraffa | Giraffa_giraffa | giraffes |
| ReticulatedGiraffe | Giraffa | reticulata | NA | Giraffa_reticulata | giraffes |
| LuangwaGiraffe | Giraffa | tippelskirchi | thornicrofti | Giraffa_tippelskirchi | giraffes |
| MasaiGiraffe | Giraffa | tippelskirchi | tippelskirchi | Giraffa_tippelskirchi | giraffes |
| BwindiGorilla | Gorilla | beringei | beringei | Gorilla_beringei | gorillas |
| EasternLowlandGorilla | Gorilla | beringei | graueri | Gorilla_beringei | gorillas |
| VirungaGorilla | Gorilla | beringei | beringei | Gorilla_beringei | gorillas |
| CrossRiverGorilla | Gorilla | gorilla | diehli | Gorilla_gorilla | gorillas |
| WesternLowlandGorilla | Gorilla | gorilla | gorilla | Gorilla_gorilla | gorillas |
| SunBear | Helarctos | malayanus | NA | Helarctos_malayanus | ursids |
| CommonHippo | Hippopotamus | amphibius | NA | Hippopotamus_amphibius | hippos |
| StripedHyena | Hyaena | hyaena | NA | Hyaena_hyaena | hyenas |
| Waterdeer | Hydropotes | inermis | NA | Hydropotes_inermis | capreolini |
| FraserDolphin | Lagenodelphis | hosei | NA | Lagenodelphis_hosei | dolphins |
| WildGuanaco | Lama | guanicoe | NA | Lama_guanicoe | camelids |
| Lama | Lama | lama | NA | Lama_lama | camelids |
| Alpaca | Lama | pacos | NA | Lama_pacos | camelids |
| WildVicuna | Lama | vicugna | NA | Lama_vicugna | camelids |
| ForestElephant | Loxodonta | africana | NA | Loxodonta_africana | elephants |
| SavannaElephant | Loxodonta | cyclotis | NA | Loxodonta_cyclotis | elephants |
| CanadianLynx | Lynx | canadensis | NA | Lynx_canadensis | lynx |
| NorthernLynx | Lynx | lynx | lynx | Lynx_lynx | lynx |
| SiberianLynx | Lynx | lynx | wrangeli | Lynx_lynx | lynx |
| IberianLynx | Lynx | pardinus | NA | Lynx_pardinus | lynx |
| Bobcat | Lynx | rufus | NA | Lynx_rufus | lynx |
| WoollyMammoth | Mammuthus | primigenius | NA | Mammuthus_primigenius | elephants |
| SlothBear | Melursus | ursinus | NA | Melursus_ursinus | ursids |
| MuleDeer | Odocoileus | hemionus | hemionus | Odoceileus_hemionus | odocoileus |
| SitkaDeer | Odocoileus | hemionus | sitkensis | Odoceileus_hemionus | odocoileus |
| WhitetailedDeer | Odocoileus | virginianus | NA | Odoceileus_virginianus | odocoileus |
| Okapi | Okapia | johnstoni | NA | Okapia_johnstoni | giraffes |
| AlaskaResidentOrca | Orcinus | orca | ater | Orcinus_orca | killerwhales |
| AntarcticOrca | Orcinus | orca | orca | Orcinus_orca | killerwhales |
| NorthAtlanticOrca | Orcinus | orca | orca | Orcinus_orca | killerwhales |
| PacificOffshoreOrca | Orcinus | orca | orca | Orcinus_orca | killerwhales |
| SouthernResidentOrca | Orcinus | orca | ater | Orcinus_orca | killerwhales |
| TransientOrca | Orcinus | orca | rectipinnus | Orcinus_orca | killerwhales |
| BighornSheep | Ovis | canadensis | NA | Ovis_canadensis | sheep |
| ThinhornSheep | Ovis | dalli | NA | Ovis_dalli | sheep |
| StraightTuskedElephant | Palaeoloxodon | antiquus | NA | Palaeoloxodon_antiquus | elephants |
| Bonobo | Pan | paniscus | NA | Pan_paniscus | chimpanzees |
| Chimpanzee | Pan | troglodytes | NA | Pan_troglodytes | chimpanzees |
| AsiaticLion | Panthera | leo | leo | Panthera_leo | panthera |
| EasternLion | Panthera | leo | melanochaita | Panthera_leo | panthera |
| NorthAfricanLion | Panthera | leo | leo | Panthera_leo | panthera |
| SouthernLion | Panthera | leo | melanochaita | Panthera_leo | panthera |
| WesternLion | Panthera | leo | leo | Panthera_leo | panthera |
| Jaguar | Panthera | onca | NA | Panthera_onca | panthera |

| | | | | | |
|---|---|---|---|---|---|
| AmurLeopard | Panthera | pardus | orientalis | Panthera_pardus | panthera |
| SouthafricanLeopard | Panthera | pardus | pardus | Panthera_pardus | panthera |
| AmurTiger | Panthera | tigris | altaica | Panthera_tigris | panthera |
| BengalTiger | Panthera | tigris | tigris | Panthera_tigris | panthera |
| SouthChinaTiger | Panthera | tigris | tigris | Panthera_tigris | panthera |
| SumatranTiger | Panthera | tigris | sondaica | Panthera_tigris | panthera |
| SnowLeopard | Panthera | uncia | NA | Panthera_uncia | panthera |
| BrownHyena | Parahyaena | brunnea | NA | Parahyaena_brunnea | hyenas |
| SumatranOrang | Pongo | abelii | NA | Pongo_abelii | orangutan |
| BorneanOrang | Pongo | pygmaeus | NA | Pongo_pygmaeus | orangutan |
| TapanuliOrang | Pongo | tapanuliensis | NA | Pongo_tapanuliensis | orangutan |
| EastAardwolf | Proteles | cristatus | septentrionalis | Proteles_cristatus | hyenas |
| SouthAardwolf | Proteles | cristatus | cristatus | Proteles_cristatus | hyenas |
| JavanRhino | Rhinoceros | sondaicus | NA | Rhinoceros_sondaicus | rhinos |
| IndianRhino | Rhinoceros | unicornis | NA | Rhinoceros_unicornis | rhinos |
| BridledDolphin | Stenella | attenuata | NA | Stenella_attenuata | dolphins |
| StripedDolphin | Stenella | coeruleoalba | NA | Stenella_coeruleoalba | dolphins |
| AtlanticSpottedDolphin | Stenella | frontalis | NA | Stenella_frontalis | dolphins |
| SpinnerDolphin | Stenella | longirostris | NA | Stenella_longirostris | dolphins |
| AnatolianBoar | Sus | scrofa | libycus | Sus_scrofa | wildboar |
| CentralEuropeanBoar | Sus | scrofa | scrofa | Sus_scrofa | wildboar |
| NorthChinaBoar | Sus | scrofa | moupinensis | Sus_scrofa | wildboar |
| SouthChinaBoar | Sus | scrofa | moupinensis | Sus_scrofa | wildboar |
| JavaWartyPig | Sus | verrucosus | NA | Sus_verrucosus | wildboar |
| CapeBuffalo | Syncerus | caffer | caffer | Syncerus_caffer | bovini |
| ForestBuffalo | Syncerus | caffer | nanus | Syncerus_caffer | bovini |
| NileBuffalo | Syncerus | caffer | aequinoctialis | Syncerus_caffer | bovini |
| SudanBuffalo | Syncerus | caffer | brachyceros | Syncerus_caffer | bovini |
| SpectacledBear | Tremarctos | ornatus | NA | Tremarctos_ornatus | ursids |
| AustralasiaBottlenose | Tursiops | aduncus | aduncus | Tursiops_aduncus | dolphins |
| BurrunanBottlenose | Tursiops | aduncus | australis | Tursiops_aduncus | dolphins |
| IndianOceanBottlenose | Tursiops | aduncus | aduncus | Tursiops_aduncus | dolphins |
| TamanendBottlenose | Tursiops | erebennus | NA | Tursiops_erebennus | dolphins |
| BlackSeaBottlenose | Tursiops | truncatus | ponticus | Tursiops_truncatus | dolphins |
| MediterraneanBottlenose | Tursiops | truncatus | truncatus | Tursiops_truncatus | dolphins |
| OffshoreBottlenose | Tursiops | truncatus | truncatus | Tursiops_truncatus | dolphins |
| AmericanBlack | Ursus | americanus | NA | Ursus_americanus | ursids |
| BrownBear | Ursus | arctos | NA | Ursus_arctos | ursids |
| EurasianBear | Ursus | arctos | arctos | Ursus_arctos | brownbears |
| GrizzlyBear | Ursus | arctos | horribilis | Ursus_arctos | brownbears |
| HokkaidoBear | Ursus | arctos | lasiotus | Ursus_arctos | brownbears |
| KamtchatkaBear | Ursus | arctos | beringianus | Ursus_arctos | brownbears |
| KodiakBear | Ursus | arctos | middendorffi | Ursus_arctos | brownbears |
| SyrianBear | Ursus | arctos | syriacus | Ursus_arctos | brownbears |
| PolarBear | Ursus | maritimus | NA | Ursus_maritimus | ursids |
| CaveBear | Ursus | spelaeus | NA | Ursus_spelaeus | ursids |
| AsiaticBlack | Ursus | thibetanus | NA | Ursus_thibetanus | ursids |
| JapanBlack | Ursus | thibetanus | japonicus | Ursus_thibetanus | ursids |

**Tables S1B.** Accession codes of samples.

| sample | ID | dataset |
|---|---|---|
| Bison1 | SRR14765467 | bovini |
| Bison2 | SRR14765461 | bovini |
| Bonasus1 | SRR3178073 | bovini |
| Bonasus2 | SRR3178074 | bovini |
| Gaurus1 | ERR3305589 | bovini |
| Gaurus2 | ERR7198375 | bovini |
| Indicus1 | SRR26321663 | bovini |
| Indicus2 | SRR26321667 | bovini |
| Javanicus1 | SRR28144104 | bovini |
| Javanicus2 | SRR14765489 | bovini |
| Mutus1 | SRR14685147 | bovini |
| Mutus2 | SRR9003424 | bovini |
| Primigenius1 | ERR13302290 | bovini |
| Primigenius2 | ERR13302594 | bovini |
| Sauveli1 | SRR16018328 | bovini |
| Sauveli2 | SRR16018330 | bovini |
| Taurus1 | ERR12373333 | bovini |
| Taurus2 | ERR12373334 | bovini |
| Asia1 | ERR4495084 | bovini |
| Asia2 | ERR4414062 | bovini |
| Cape1 | ERR11802505 | bovini |
| Cape2 | ERR11785784 | bovini |
| Euro1 | ERR4495068 | bovini |
| Euro2 | ERR4495071 | bovini |
| Forest1 | ERR11802525 | bovini |
| Forest2 | ERR11802524 | bovini |
| LowAnoa1 | SRR21016826 | bovini |
| Nile1 | ERR11867688 | bovini |
| Nile2 | ERR11867689 | bovini |
| Sudan1 | ERR11867708 | bovini |
| Sudan2 | ERR11867709 | bovini |
| Swamp1 | SRR12915633 | bovini |
| Swamp2 | SRR22557457 | bovini |
| CMB001 | SAMN06759210 | camelids |
| CMB001 | SAMN06759210 | camelids |
| CMB002 | SAMN06759208 | camelids |
| CMB002 | SAMN06759208 | camelids |
| CMD001 | SAMN06759213 | camelids |
| CMD001 | SAMN06759213 | camelids |
| CMD002 | SAMN06759211 | camelids |
| CMD002 | SAMN06759211 | camelids |
| CMF001 | SAMN06759129 | camelids |
| CMF001 | SAMN06759129 | camelids |
| CMF002 | SAMN06759136 | camelids |
| CMF002 | SAMN06759136 | camelids |
| LMC001 | SAMN38341426 | camelids |
| LMC001 | SAMN38341426 | camelids |
| LMC002 | SAMN38341426 | camelids |
| LMC002 | SAMN38341426 | camelids |
| LMG001 | SAMN14360337 | camelids |
| LMG001 | SAMN14360337 | camelids |

| | | |
|---|---|---|
| LMG002 | SAMN14360341 | camelids |
| LMG002 | SAMN14360341 | camelids |
| VIP001 | SAMN02996813 | camelids |
| VIP001 | SAMN02996813 | camelids |
| VIP002 | SAMN02996813 | camelids |
| VIP002 | SAMN02996813 | camelids |
| VIV001 | SAMN14360350 | camelids |
| VIV001 | SAMN14360350 | camelids |
| VIV002 | SAMN14360349 | camelids |
| VIV002 | SAMN14360349 | camelids |
| Dromedary | SAMEA117485871 | camelids |
| Dromedary | SAMEA117485873 | camelids |
| AtlanticCoastal1 | SRR10839565 | dolphins |
| AtlanticCoastal2 | SRR10839564 | dolphins |
| AtlanticCoastal3 | SRR10839563 | dolphins |
| AtlanticCoastal4 | SRR10839562 | dolphins |
| AtlanticCoastal5 | SRR10839561 | dolphins |
| AtlanticCoastal6 | SRR10839559 | dolphins |
| AtlanticSpotted1 | SRR10839582 | dolphins |
| Australasia1 | SRR10839583 | dolphins |
| Australasia2 | SRR10839581 | dolphins |
| Australasia3 | SRR10839580 | dolphins |
| Australasia4 | SRR10839579 | dolphins |
| Australasia5 | SRR10839578 | dolphins |
| Australasia6 | SRR10839577 | dolphins |
| Australasia7 | SRR10839576 | dolphins |
| Australasia8 | SRR10839575 | dolphins |
| BlackSea10 | SRR10839537 | dolphins |
| BlackSea1 | SRR10839539 | dolphins |
| BlackSea2 | SRR10839532 | dolphins |
| BlackSea3 | SRR10839531 | dolphins |
| BlackSea4 | SRR10839533 | dolphins |
| BlackSea5 | SRR10839535 | dolphins |
| BlackSea7 | SRR10839536 | dolphins |
| BlackSea8 | SRR10839534 | dolphins |
| BridledDolphin1 | SRR10839571 | dolphins |
| Burrunan1 | SRR10839574 | dolphins |
| Burrunan2 | SRR10839573 | dolphins |
| Burrunan3 | SRR10839572 | dolphins |
| Burrunan4 | SRR10839570 | dolphins |
| Burrunan5 | SRR10839569 | dolphins |
| Burrunan6 | SRR10839568 | dolphins |
| Burrunan7 | SRR10839567 | dolphins |
| Burrunan8 | SRR10839566 | dolphins |
| CommonDolphin1 | SRR10839560 | dolphins |
| FraserDolphin1 | SRR10839549 | dolphins |
| FraserDolphin2 | SRR10839538 | dolphins |
| IndianOcean1 | SRR10839589 | dolphins |
| IndianOcean2 | SRR10839588 | dolphins |
| IndianOcean3 | SRR10839587 | dolphins |
| IndianOcean4 | SRR10839586 | dolphins |
| IndianOcean5 | SRR10839585 | dolphins |
| IndianOcean6 | SRR10839584 | dolphins |
| Mediterranean1 | SRR10839548 | dolphins |

| | | |
|---|---|---|
| Mediterranean2 | SRR10839547 | dolphins |
| Mediterranean3 | SRR10839546 | dolphins |
| Mediterranean4 | SRR10839545 | dolphins |
| Mediterranean5 | SRR10839544 | dolphins |
| Mediterranean6 | SRR10839543 | dolphins |
| Mediterranean7 | SRR10839542 | dolphins |
| Mediterranean8 | SRR10839541 | dolphins |
| Mediterranean9 | SRR10839540 | dolphins |
| NAO1 | SRR10839558 | dolphins |
| NAO2 | SRR10839557 | dolphins |
| NAO3 | SRR10839556 | dolphins |
| NAO4 | SRR10839555 | dolphins |
| NAO5 | SRR10839554 | dolphins |
| NAO6 | SRR10839553 | dolphins |
| Oman1 | SRR10839590 | dolphins |
| Pakistan1 | SRR10839591 | dolphins |
| RoughToothed2 | SRR10839594 | dolphins |
| SpinnerDolphin1 | SRR10839529 | dolphins |
| SpinnerDolphin2 | SRR10839528 | dolphins |
| StripedDolphin1 | SRR10839527 | dolphins |
| StripedDolphin2 | SRR10839592 | dolphins |
| Taiwan1 | SRR10839552 | dolphins |
| Taiwan2 | SRR10839551 | dolphins |
| BorneanElephant | Manari (Kappelhof et al. 2025) | elephants |
| BorneanElephant | Sayang (Kappelhof et al. 2025) | elephants |
| SumatranElephant | Valentino (Kappelhof et al. 2025) | elephants |
| SumatranElephant | Cynthia (Kappelhof et al. 2025) | elephants |
| SriLankanElephant | Jarnitha (Kappelhof et al. 2025) | elephants |
| SriLankanElephant | CeylaHimali (Kappelhof et al. | elephants |
| VietnamElephant | Douanita (Kappelhof et al. 2025) | elephants |
| VietnamElephant | Delhi (Kappelhof et al. 2025) | elephants |
| ForestElephant | SAMEA115942074 | elephants |
| ForestElephant | SAMEA115942079 | elephants |
| ForestElephant | SAMEA115942097 | elephants |
| StraightTuskedElephant | SAMEA104469187 | elephants |
| AsianElephant3 | SRR25983393 | elephants |
| AsianElephant4 | SRR25983393 | elephants |
| ForestElephant3 | ERR14017657 | elephants |
| ForestElephant4 | ERR14017679 | elephants |
| ForestElephant5 | ERR14017762 | elephants |
| WoollyMammoth1 | ERR10173417 | elephants |
| WoollyMammoth2 | ERR10173468 | elephants |
| Equus przewalski | SAMN16229270 | equus |
| Equus przewalski | SAMN16229271 | equus |
| E. hemionus hemippus | SAMEA9991232 | equus |
| Equus caballus | SAMN36387955 | equus |
| Equus caballus | SAMN36387966 | equus |
| Equus asinus | SAMN31557344 | equus |
| Equus asinus | SAMN13284259 | equus |
| Equus quagga chapmani | SAMN17036589 | equus |
| Equus quagga chapmani | SAMN17036593 | equus |
| Equus zebra | SAMN15801456 | equus |
| Equus grevyi | SAMEA110663650 | equus |
| Equus hemionus | SAMEA110663661 | equus |

| | | |
|---|---|---|
| Equus kiang | SAMEA110663659 | equus |
| Equus kiang | SAMEA110663660 | equus |
| Equus asinus somalicus | SAMEA110663663 | equus |
| MasaiGiraffe | LVNP804 | giraffes |
| MasaiGiraffe | LVNP808 | giraffes |
| MasaiGiraffe | LVNP8-09 | giraffes |
| MasaiGiraffe | LVNP8-10 | giraffes |
| MasaiGiraffe | LVNP8-12 | giraffes |
| MasaiGiraffe | GF246 | giraffes |
| MasaiGiraffe | GF248 | giraffes |
| MasaiGiraffe | GF249 | giraffes |
| MasaiGiraffe | GF250 | giraffes |
| MasaiGiraffe | GF253 | giraffes |
| NorthernGiraffe | GNP01 | giraffes |
| NorthernGiraffe | GNP04 | giraffes |
| NorthernGiraffe | GNP05 | giraffes |
| NorthernGiraffe | WA733 | giraffes |
| NorthernGiraffe | WA746 | giraffes |
| NorthernGiraffe | WA806 | giraffes |
| NorthernGiraffe | GF261 | giraffes |
| NorthernGiraffe | GF262 | giraffes |
| NorthernGiraffe | GF263 | giraffes |
| NorthernGiraffe | GF264 | giraffes |
| Okapi | WOAK | giraffes |
| ReticulatedGiraffe | GF227 | giraffes |
| ReticulatedGiraffe | GF228 | giraffes |
| ReticulatedGiraffe | GF229 | giraffes |
| ReticulatedGiraffe | GF230 | giraffes |
| ReticulatedGiraffe | GF231 | giraffes |
| ReticulatedGiraffe | GF232 | giraffes |
| ReticulatedGiraffe | GF233 | giraffes |
| ReticulatedGiraffe | GF234 | giraffes |
| ReticulatedGiraffe | GF235 | giraffes |
| ReticulatedGiraffe | GF236 | giraffes |
| SouthernGiraffe | KKR01 | giraffes |
| SouthernGiraffe | KKR02 | giraffes |
| SouthernGiraffe | KKR03 | giraffes |
| SouthernGiraffe | KKR04 | giraffes |
| SouthernGiraffe | HSBM053 | giraffes |
| SouthernGiraffe | HSBM075 | giraffes |
| SouthernGiraffe | HSBM062 | giraffes |
| SouthernGiraffe | BVC10 | giraffes |
| SouthernGiraffe | HNBF035 | giraffes |
| SouthernGiraffe | HNBF037 | giraffes |
| Pongo tapanuliensis | SAMN00007170 | gnous |
| Pongo pygmaeus | SAMN10521809 | gnous |
| Pongo pygmaeus | SAMEA112483015 | gnous |
| Pongo abelii | SAMN01920544 | gnous |
| Pongo abelii | SAMN10521808 | gnous |
| Pan troglodytes | SAMN29543813 | gnous |
| Pan troglodytes | SAMN29543773 | gnous |
| Pan troglodytes verus | SAMEA4374765 | gnous |
| Pan troglodytes verus | SAMEA4374763 | gnous |
| Pan paniscus | SAMN35877942 | gnous |

| | | |
|---|---|---|
| Pan paniscus | SAMN35877943 | gnous |
| Connochaetes taurinus | SAMN39917060 | gnous |
| Connochaetes taurinus | SAMN39917061 | gnous |
| Connochaetes taurinus | SAMN39917031 | gnous |
| Connochaetes taurinus | SAMN39917109 | gnous |
| Connochaetes taurinus | SAMN39917108 | gnous |
| Connochaetes taurinus | SAMN39917057 | gnous |
| Connochaetes taurinus | SAMN39916989 | gnous |
| Connochaetes taurinus | SAMN39917122 | gnous |
| Connochaetes taurinus | SAMN39917131 | gnous |
| Connochaetes taurinus | SAMN39917130 | gnous |
| Connochaetes gnou | SAMN39916989 | gnous |
| Connochaetes gnou | SAMN39917122 | gnous |
| Alcelaphus buselaphus | SAMN39917131 | gnous |
| Alcelaphus buselaphus | SAMN39917130 | gnous |
| WesternLowland1 | SAMN01920503 | gorillas |
| WesternLowland2 | SAMN01920497 | gorillas |
| WesternLowland3 | SAMN01920501 | gorillas |
| CrossRiver1 | SAMN01920476 | gorillas |
| EasternLowland1 | SAMN01920473 | gorillas |
| EasternLowland2 | SAMN01920474 | gorillas |
| Virunga1 | SAMEA1692350 | gorillas |
| Virunga2 | SAMEA1692353 | gorillas |
| Virunga3 | SAMEA2697038 | gorillas |
| Bwindi1 | SAMEA3939557 | gorillas |
| Bwindi2 | SAMEA3939561 | gorillas |
| Brown1 | SRR5886633 | hyenas |
| Brown2 | SRR5886638 | hyenas |
| Cave1 | SRR9914660 | hyenas |
| Cave2 | SRR9914657 | hyenas |
| Cave3 | SRR9914655 | hyenas |
| Eastaard1 | SRR13177417 | hyenas |
| Southaard1 | SRR13177419 | hyenas |
| Southaard2 | SRR13177420 | hyenas |
| Spotted1 | SRR9914668 | hyenas |
| Spotted2 | SRR9914667 | hyenas |
| Striped1 | SRR5904109 | hyenas |
| Striped2 | SRR11430567 | hyenas |
| AlaskaResident16 | SRR1324596 | killerwhales |
| AlaskaResident17 | SRR1324597 | killerwhales |
| AlaskaResident19 | SRR1584023 | killerwhales |
| AlaskaResident20 | SRR1584027 | killerwhales |
| AlaskaResident24 | SRR1584031 | killerwhales |
| AlaskaTransient39 | SRR1324601 | killerwhales |
| AlaskaTransient42 | SRR1324603 | killerwhales |
| AlaskaTransient47 | SRR1584083 | killerwhales |
| AlaskaTransient48 | SRR1584089 | killerwhales |
| AlaskaTransient53 | SRR1584099 | killerwhales |
| BeringSea116 | SRR1584175 | killerwhales |
| BeringSea117 | SRR1324624 | killerwhales |
| BeringSea120 | SRR1324625 | killerwhales |
| BeringSea122 | SRR1584185 | killerwhales |
| BeringSea127 | SRR1584195 | killerwhales |
| CaliforniaTransient62 | SRR1324606 | killerwhales |

| | | |
|---|---|---|
| CaliforniaTransient63 | SRR1584121 | killerwhales |
| CaliforniaTransient66 | SRR1584127 | killerwhales |
| CaliforniaTransient68 | SRR1584131 | killerwhales |
| CaliforniaTransient75 | SRR1584141 | killerwhales |
| Iceland82 | SRR1584153 | killerwhales |
| Iceland86 | SRR1324611 | killerwhales |
| Iceland87 | SRR1324612 | killerwhales |
| Iceland91 | SRR1584157 | killerwhales |
| Iceland92 | SRR1324613 | killerwhales |
| MarionIsland131 | SRR1324629 | killerwhales |
| MarionIsland133 | SRR1324629 | killerwhales |
| MarionIsland134 | SRR1324630 | killerwhales |
| MarionIsland141 | SRR1324632 | killerwhales |
| MarionIsland143 | SRR1584239 | killerwhales |
| PacificOffshore102 | SRR1324618 | killerwhales |
| PacificOffshore26 | SRR1584039 | killerwhales |
| PacificOffshore96 | SRR1324615 | killerwhales |
| PacificOffshore97 | SRR1584159 | killerwhales |
| PacificOffshore99 | SRR1324617 | killerwhales |
| Russia105 | SRR1324619 | killerwhales |
| Russia109 | SRR1324620 | killerwhales |
| Russia110 | SRR1324621 | killerwhales |
| Russia111 | SRR1324622 | killerwhales |
| Russia113 | SRR1324623 | killerwhales |
| SouthernResident11 | SRR1584001 | killerwhales |
| SouthernResident12 | SRR1324595 | killerwhales |
| SouthernResident15 | SRR1584013 | killerwhales |
| SouthernResident2 | SRR1583985 | killerwhales |
| SouthernResident7 | SRR1583997 | killerwhales |
| Bobcat1 | ERR2737229 | lynx |
| Bobcat2 | SRR6071633 | lynx |
| Canadian1 | ERR2861644 | lynx |
| Canadian2 | SRR6071634 | lynx |
| Iberian1 | ERR5922475 | lynx |
| Iberian2 | ERR5922477 | lynx |
| Iberian3 | ERR5922483 | lynx |
| Norway1 | ERR5922462 | lynx |
| Norway2 | ERR5922464 | lynx |
| RussiaWest1 | ERR5922448 | lynx |
| RussiaWest2 | ERR5922450 | lynx |
| Yakutia1 | ERR2737237 | lynx |
| Yakutia2 | ERR2737233 | lynx |
| Odocoileus_hemionus | SRR18858846 | odoicoileus |
| Odocoileus_virgianus | SRR18858848 | odoicoileus |
| Odocoileus_virgianus | SRR18858863 | odoicoileus |
| Odocoileus_hemionus | SRR18858865 | odoicoileus |
| Odocoileus_hemionus | SRR18858869 | odoicoileus |
| Odocoileus_hemionus | SRR23126248 | odoicoileus |
| Odocoileus_virgianus | SRR23126253 | odoicoileus |
| Panthera tigris sumatrae | SAMN09080450 | panthera |
| Panthera tigris sumatrae | SAMN09080447 | panthera |
| Panthera tigris altaica | SAMN09770319 | panthera |
| Panthera tigris altaica | SAMN09770316 | panthera |
| Panthera tigris amoyensis | SAMN09770323 | panthera |

| | | |
|---|---|---|
| Panthera tigris amoyensis | SAMN09770321 | panthera |
| Panthera tigris virgata | SAMN27163016 | panthera |
| Panthera tigris virgata | SAMN27163014 | panthera |
| Panthera tigris tigris | SAMN12549969 | panthera |
| Panthera tigris tigris | SAMN17487598 | panthera |
| Panthera leo persica | SAMN06606823 | panthera |
| Panthera leo persica | SAMN14352183 | panthera |
| Panthera spelaea | SAMN14352179 | panthera |
| Panthera leo | SAMN14352190 | panthera |
| Western lion | SAMN14352197 | panthera |
| Western lion | SAMN14352198 | panthera |
| Eastern lion | SAMN14352186 | panthera |
| Eastern lion | SAMN14352187 | panthera |
| Southern lion | SAMN14352195 | panthera |
| Southern lion | SAMN14352192 | panthera |
| Panthera onca | SAMN29795438 | panthera |
| Panthera onca | SAMN29795432 | panthera |
| Panthera pardus (Southern | SAMEA8520932 | panthera |
| Panthera pardus (Southern | SAMEA8520941 | panthera |
| Panthera pardus (Amur) | SAMEA8520936 | panthera |
| Panthera pardus (Amur) | SAMN04361361 | panthera |
| Panthera uncia | SAMN13911153 | panthera |
| Panthera uncia | SAMN02086968 | panthera |
| BlackRhinoC1 | SRR25686487 | rhinos |
| BlackRhinoC2 | SRR25686488 | rhinos |
| BlackRhinoS1 | SRR25686483 | rhinos |
| Elasmotherium1 | SRR13333569 | rhinos |
| IndianRhino1 | DRR308100 | rhinos |
| IndianRhino2 | SRR13627513 | rhinos |
| JavanRhino1 | SRR13316980 | rhinos |
| MercksRhino1 | SRR13310806 | rhinos |
| SumatranRhino1 | ERR3677658 | rhinos |
| SumatranRhino2 | ERR3677654 | rhinos |
| WhiteRhinoNWR1 | SRR5852742 | rhinos |
| WhiteRhinoNWR2 | SRR5852743 | rhinos |
| WhiteRhinoSWR1 | SRR5852738 | rhinos |
| WhiteRhinoSWR2 | SRR20637441 | rhinos |
| WoollyRhino1 | SRR13309646 | rhinos |
| BighornSheep1 | SRR16036488 | sheep |
| BighornSheep2 | SRR16036486 | sheep |
| BighornSheep3 | SRR16036516 | sheep |
| ThinhornSheep1 | SRR16036497 | sheep |
| ThinhornSheep2 | SRR16036493 | sheep |
| ThinhornSheep3 | SRR16036492 | sheep |
| NearEast1 | SAMEA3497884 | wildboar |
| NearEast2 | SAMEA3497885 | wildboar |
| Netherlands1 | SAMEA3497870 | wildboar |
| Netherlands2 | SAMEA3497868 | wildboar |
| SouthChina1 | SAMEA3497815 | wildboar |
| SouthChina2 | SAMEA3497819 | wildboar |
| NorthChina1 | SAMEA3497820 | wildboar |
| NorthChina2 | SAMEA3497822 | wildboar |
| JavaWartyPig1 | SAMEA3497792 | wildboar |
| CanCo1 | SRR12075089 | wolves |

| | | |
|---|---|---|
| CanCo2 | SRR12075095 | wolves |
| CanEC1 | SRR12075085 | wolves |
| CanEC2 | SRR12075084 | wolves |
| CanGW12 | SRR8926748 | wolves |
| CanGW13 | SRR8926752 | wolves |
| CanGW14 | SRR8926751 | wolves |
| CanGW15 | SRR8926749 | wolves |
| CanGW16 | SRR8926750 | wolves |
| CanGW17 | SRR8926747 | wolves |
| CanGW1 | SRR8066608 | wolves |
| CanGW2 | SRR8066615 | wolves |
| CanGW5 | SRR8066612 | wolves |
| CanGW6 | SRR8066606 | wolves |
| CanGW7 | SRR8066607 | wolves |
| CanGW8 | SRR8066614 | wolves |
| CanGW9 | SRR8049197 | wolves |
| DAki1 | DRR336758 | wolves |
| DAki2 | DRR336759 | wolves |
| DAM1 | SRR7107992 | wolves |
| DAM2 | SRR12330325 | wolves |
| DBas1 | SRR2149861 | wolves |
| DBas2 | SRR7107885 | wolves |
| DCW1 | SRR20637403 | wolves |
| DCW2 | SRR20637404 | wolves |
| DFS1 | SRR16048637 | wolves |
| DFS2 | SRR16048635 | wolves |
| DSal1 | SRR2095503 | wolves |
| DSal2 | SRR2094408 | wolves |
| DSH1 | SRR2095539 | wolves |
| DSH2 | SRR2827574 | wolves |
| DSW1 | SRR16048618 | wolves |
| DSW2 | SRR16048617 | wolves |
| FinGW1 | ERR4317970 | wolves |
| FinGW2 | ERR4317955 | wolves |
| FinGW3 | ERR4317903 | wolves |
| FinGW4 | ERR4317975 | wolves |
| GreGW1 | SRR8049195 | wolves |
| OAGW1 | SRR8049196 | wolves |
| OGJ1 | SRR8049192 | wolves |
| RusGW1 | ERR4318094 | wolves |
| RusGW2 | ERR4318095 | wolves |
| RusGW3 | ERR4318096 | wolves |
| RusGW4 | ERR4318097 | wolves |
| RusGW5 | ERR4318101 | wolves |
| RusGW6 | ERR4318102 | wolves |
| RusGW9 | SRR7107647 | wolves |
| ScaGW1 | ERR5903368 | wolves |
| ScaGW2 | ERR5906614 | wolves |
| ScaGW3 | ERR5906617 | wolves |
| ScaGW4 | ERR5906616 | wolves |

**Table S2**. List of all lineage pairs. Numbers in the column 'index' correspond to supplementary figure 4A. Lineage pairs for which only female hybrids are fertile ('females_only' in the column 'hybridisation'), display Haldane's Rule, while 'genetic isolation' indicates the maintenance of separate sympatric gene pools, even though hybrid offspring is fertile regardless of sex. The column 'predict' indicates whether the taxonomic level of the lineage pair is correctly predicted based on a Dxy- and Fst-threshold of 0.225% and 0.26, respectively. TP = true positive, FP = false positive, TN = true negative, FN = false negative.

| Lineage2 | Lineage1 | hybridisation | dataset | level | predict | index |
|---|---|---|---|---|---|---|
| AfricanWolf | ArcticWolf | unknown | wolves | species | TP | 1 |
| AfricanWolf | EurasianWolf | unknown | wolves | species | TP | 2 |
| AfricanWolf | EasternCoyote | unknown | wolves | species | TP | 3 |
| AfricanWolf | Coyote | unknown | wolves | species | TP | 4 |
| AfricanWolf | Dog | unknown | wolves | species | TP | 5 |
| AmericanBison | RiverBuffalo | neither sex | bovini | genera | NA | 6 |
| AmurTiger | AmurLeopard | unknown | panthera | species | TP | 7 |
| AngolanGiraffe | KordofanGiraffe | unknown | giraffes | species | TP | 8 |
| AngolanGiraffe | MasaiGiraffe | unknown | giraffes | species | TP | 9 |
| AngolanGiraffe | ReticulatedGiraffe | unknown | giraffes | species | TP | 10 |
| AngolanGiraffe | NubianGiraffe | unknown | giraffes | species | TP | 11 |
| AngolanGiraffe | SouthAfricanGiraffe | both sexes | giraffes | sub | TN | 12 |
| AntarcticOrca | AlaskaResidentOrca | both sexes | killerwhales | sub | TN | 13 |
| AntarcticOrca | TransientOrca | both sexes | killerwhales | sub | TN | 14 |
| AntarcticOrca | NorthAtlanticOrca | both sexes | killerwhales | sub | TN | 15 |
| ArcticWolf | Coyote | genetic_isolation | wolves | species | TP | 16 |
| ArcticWolf | EasternCoyote | unknown | wolves | species | TP | 17 |
| AsianElephant | SavannaElephant | neither sex | elephants | genera | NA | 18 |
| AsiaticBlack | BrownBear | unknown | ursids | species | TP | 19 |
| AsiaticBlack | AmericanBlack | unknown | ursids | species | TP | 20 |
| AsiaticLion | AmurLeopard | unknown | panthera | species | TP | 21 |
| AsiaticLion | AmurTiger | females only | panthera | species | TP | 22 |
| AtlanticSpottedDolphin | TamanendBottlenose | unknown | dolphins | genera | NA | 23 |
| AustralasiaBottlenose | TamanendBottlenose | unknown | dolphins | species | TP | 24 |
| AustralasiaBottlenose | AtlanticSpottedDolphin | unknown | dolphins | genera | NA | 25 |
| BactrianCamel | Alpaca | neither sex | camelids | genera | NA | 26 |
| BactrianDeer | CaspianRedDeer | unknown | cervus | species | TP | 27 |
| BactrianDeer | TianshanWapiti | unknown | cervus | species | TP | 28 |
| BengalTiger | AsiaticLion | females only | panthera | species | TP | 29 |
| BengalTiger | AmurTiger | both sexes | panthera | sub | TN | 30 |
| BengalTiger | AmurLeopard | unknown | panthera | species | TP | 31 |
| BlackSeaBottlenose | AtlanticSpottedDolphin | unknown | dolphins | genera | NA | 32 |
| BlackSeaBottlenose | AustralasiaBottlenose | unknown | dolphins | species | TP | 33 |
| BlackSeaBottlenose | TamanendBottlenose | unknown | dolphins | species | FN | 34 |
| BorneanElephant | SavannaElephant | neither sex | elephants | genera | NA | 35 |
| BorneanElephant | AsianElephant | both sexes | elephants | sub | TN | 36 |
| BridledDolphin | TamanendBottlenose | unknown | dolphins | genera | NA | 37 |
| BridledDolphin | BlackSeaBottlenose | unknown | dolphins | genera | NA | 38 |
| BridledDolphin | AtlanticSpottedDolphin | unknown | dolphins | species | TP | 39 |
| BridledDolphin | AustralasiaBottlenose | unknown | dolphins | genera | NA | 40 |
| BrownBear | AmericanBlack | unknown | ursids | species | TP | 41 |
| BurrunanBottlenose | BlackSeaBottlenose | unknown | dolphins | species | TP | 42 |
| BurrunanBottlenose | TamanendBottlenose | unknown | dolphins | species | TP | 43 |
| BurrunanBottlenose | AustralasiaBottlenose | both sexes | dolphins | sub | TN | 44 |

| | | | | | | |
|---|---|---|---|---|---|---|
| BurrunanBottlenose | AtlanticSpottedDolphin | unknown | dolphins | genera | NA | 45 |
| BurrunanBottlenose | BridledDolphin | unknown | dolphins | genera | NA | 46 |
| CanadianLynx | Bobcat | genetic_isolation | lynx | species | TP | 47 |
| CapeBuffalo | RiverBuffalo | neither sex | bovini | genera | NA | 48 |
| CapeBuffalo | AmericanBison | neither sex | bovini | genera | NA | 49 |
| CapeBuffalo | EuropeanBison | neither sex | bovini | genera | NA | 50 |
| CaspianRedDeer | TianshanWapiti | both sexes | cervus | species | TP | 51 |
| Cattle | ForestBuffalo | neither sex | bovini | genera | NA | 52 |
| Cattle | EuropeanBison | females only | bovini | species | TP | 53 |
| Cattle | Gaur | females only | bovini | species | TP | 54 |
| Cattle | CapeBuffalo | neither sex | bovini | genera | NA | 55 |
| Cattle | Yak | females only | bovini | species | TP | 56 |
| Cattle | RiverBuffalo | neither sex | bovini | genera | NA | 57 |
| Cattle | Zebu | unknown | bovini | species | TP | 58 |
| Cattle | SwampBuffalo | neither sex | bovini | genera | NA | 59 |
| Cattle | SudanBuffalo | neither sex | bovini | genera | NA | 60 |
| Cattle | NileBuffalo | neither sex | bovini | genera | NA | 61 |
| Cattle | AmericanBison | females only | bovini | species | TP | 62 |
| Cattle | LowlandAnoa | neither sex | bovini | genera | NA | 63 |
| CaveBear | AmericanBlack | unknown | ursids | species | TP | 64 |
| CaveBear | BrownBear | unknown | ursids | species | TP | 65 |
| CaveBear | JapanBlack | unknown | ursids | species | TP | 66 |
| CaveBear | AsiaticBlack | unknown | ursids | species | TP | 67 |
| CentralEuropeanBoar | AnatolianBoar | both sexes | wildboar | sub | TN | 68 |
| CentralEuropeanRedDeer | BactrianDeer | unknown | cervus | species | TP | 69 |
| CentralEuropeanRedDeer | CaspianRedDeer | both sexes | cervus | sub | TN | 70 |
| CentralEuropeanRedDeer | TianshanWapiti | both sexes | cervus | species | TP | 71 |
| ChapmanZebra | AfricanWildass | neither sex | equus | species | TP | 72 |
| Chimpanzee | Bonobo | unknown | chimpanzees | species | TP | 73 |
| CommonDolphin | BlackSeaBottlenose | unknown | dolphins | genera | NA | 74 |
| CommonDolphin | BurrunanBottlenose | unknown | dolphins | genera | NA | 75 |
| CommonDolphin | TamanendBottlenose | unknown | dolphins | genera | NA | 76 |
| CommonDolphin | AustralasiaBottlenose | unknown | dolphins | genera | NA | 77 |
| CommonDolphin | BridledDolphin | unknown | dolphins | genera | NA | 78 |
| CommonDolphin | AtlanticSpottedDolphin | unknown | dolphins | genera | NA | 79 |
| CommonHippo | PygmyHippo | unknown | hippos | genera | NA | 80 |
| CrossRiverGorilla | BwindiGorilla | unknown | gorillas | species | TP | 81 |
| Dog | ArcticWolf | both sexes | wolves | sub | TN | 82 |
| Dog | EasternCoyote | unknown | wolves | species | TP | 83 |
| Dog | Coyote | unknown | wolves | species | TP | 84 |
| DomesticDonkey | AfricanWildass | both sexes | equus | sub | TN | 85 |
| DomesticDonkey | ChapmanZebra | neither sex | equus | species | TP | 86 |
| DomesticHorse | AfricanWildass | neither sex | equus | species | TP | 87 |
| DomesticHorse | ChapmanZebra | neither sex | equus | species | TP | 88 |
| DomesticHorse | DomesticDonkey | neither sex | equus | species | TP | 89 |
| Dromedary | BactrianCamel | genetic_isolation | camelids | species | TP | 90 |
| Dromedary | Alpaca | neither sex | camelids | genera | NA | 91 |
| EastAardwolf | BrownHyena | unknown | hyenas | genera | NA | 92 |
| EastBlueWildebeest | BlackWildebeest | unknown | gnous | species | TP | 93 |
| EasternCoyote | Coyote | both sexes | wolves | sub | TN | 94 |
| EasternLion | BengalTiger | females only | panthera | species | TP | 95 |
| EasternLion | AsiaticLion | both sexes | panthera | sub | TN | 96 |
| EasternLion | AmurTiger | females only | panthera | species | TP | 97 |
| EasternLion | AmurLeopard | unknown | panthera | species | TP | 98 |
| EasternLowlandGorilla | BwindiGorilla | both sexes | gorillas | sub | TN | 99 |
| EasternLowlandGorilla | CrossRiverGorilla | unknown | gorillas | species | TP | 100 |
| EurasianBear | KodiakBear | both sexes | brownbears | sub | TN | 101 |
| EurasianBear | KamtchatkaBear | both sexes | brownbears | sub | TN | 102 |
| EurasianBear | HokkaidoBear | both sexes | brownbears | sub | FP | 103 |
| EurasianBear | GrizzlyBear | both sexes | brownbears | sub | FP | 104 |
| EurasianBear | SyrianBear | both sexes | brownbears | sub | FP | 105 |

| | | | | | | |
|---|---|---|---|---|---|---|
| EurasianWolf | Coyote | unknown | wolves | species | TP | 106 |
| EurasianWolf | EasternCoyote | unknown | wolves | species | TP | 107 |
| EurasianWolf | ArcticWolf | both sexes | wolves | sub | TN | 108 |
| EurasianWolf | Dog | both sexes | wolves | sub | TN | 109 |
| EuropeanBison | RiverBuffalo | neither sex | bovini | genera | NA | 110 |
| EuropeanBison | AmericanBison | unknown | bovini | species | TP | 111 |
| FinWhale | BlueWhale | unknown | finwhales | species | TP | 112 |
| ForestBuffalo | CapeBuffalo | both sexes | bovini | sub | FP | 113 |
| ForestBuffalo | AmericanBison | neither sex | bovini | genera | NA | 114 |
| ForestBuffalo | RiverBuffalo | neither sex | bovini | genera | NA | 115 |
| ForestBuffalo | EuropeanBison | neither sex | bovini | genera | NA | 116 |
| ForestElephant | AsianElephant | neither sex | elephants | genera | NA | 117 |
| ForestElephant | BorneanElephant | neither sex | elephants | genera | NA | 118 |
| ForestElephant | SavannaElephant | both sexes | elephants | species | TP | 119 |
| FraserDolphin | BlackSeaBottlenose | unknown | dolphins | genera | NA | 120 |
| FraserDolphin | BridledDolphin | unknown | dolphins | genera | NA | 121 |
| FraserDolphin | AustralasiaBottlenose | unknown | dolphins | genera | NA | 122 |
| FraserDolphin | TamanendBottlenose | unknown | dolphins | genera | NA | 123 |
| FraserDolphin | BurrunanBottlenose | unknown | dolphins | genera | NA | 124 |
| FraserDolphin | CommonDolphin | unknown | dolphins | genera | NA | 125 |
| FraserDolphin | AtlanticSpottedDolphin | unknown | dolphins | genera | NA | 126 |
| Gaur | CapeBuffalo | neither sex | bovini | genera | NA | 127 |
| Gaur | ForestBuffalo | neither sex | bovini | genera | NA | 128 |
| Gaur | AmericanBison | unknown | bovini | species | TP | 129 |
| Gaur | RiverBuffalo | neither sex | bovini | genera | NA | 130 |
| Gaur | EuropeanBison | unknown | bovini | species | TP | 131 |
| GoldenJackal | EasternCoyote | unknown | wolves | species | TP | 132 |
| GoldenJackal | Dog | unknown | wolves | species | TP | 133 |
| GoldenJackal | AfricanWolf | unknown | wolves | species | TP | 134 |
| GoldenJackal | EurasianWolf | unknown | wolves | species | TP | 135 |
| GoldenJackal | ArcticWolf | unknown | wolves | species | TP | 136 |
| GoldenJackal | Coyote | unknown | wolves | species | TP | 137 |
| GrevyZebra | AfricanWildass | neither sex | equus | species | TP | 138 |
| GrevyZebra | ChapmanZebra | genetic_isolation | equus | species | TP | 139 |
| GrevyZebra | DomesticDonkey | neither sex | equus | species | TP | 140 |
| GrevyZebra | DomesticHorse | neither sex | equus | species | TP | 141 |
| GrizzlyBear | KodiakBear | both sexes | brownbears | sub | FP | 142 |
| Hartebeest | EastBlueWildebeest | neither sex | gnous | genera | NA | 143 |
| Hartebeest | BlackWildebeest | neither sex | gnous | genera | NA | 144 |
| HokkaidoBear | KodiakBear | both sexes | brownbears | sub | TN | 145 |
| HokkaidoBear | SyrianBear | both sexes | brownbears | sub | FP | 146 |
| HokkaidoBear | GrizzlyBear | both sexes | brownbears | sub | FP | 147 |
| IberianLynx | Bobcat | unknown | lynx | species | TP | 148 |
| IberianLynx | CanadianLynx | unknown | lynx | species | TP | 149 |
| IberianRedDeer | KansuRedDeer | unknown | cervus | species | TP | 150 |
| IberianRedDeer | NorthAmericanWapiti | unknown | cervus | species | TP | 151 |
| IberianRedDeer | SikaDeer | unknown | cervus | species | TP | 152 |
| IberianRedDeer | BactrianDeer | unknown | cervus | species | TP | 153 |
| IberianRedDeer | CentralEuropeanRedDeer | both sexes | cervus | sub | TN | 154 |
| IberianRedDeer | TianshanWapiti | unknown | cervus | species | TP | 155 |
| IberianRedDeer | CaspianRedDeer | both sexes | cervus | sub | TN | 156 |
| IberianRoeDeer | EuropeanRoeDeer | both sexes | capreolini | sub | TN | 157 |
| IberianRoeDeer | SiberianRoeDeer | females only | capreolini | species | TP | 158 |
| IberianRoeDeer | Moose | neither sex | capreolini | genera | NA | 159 |
| IndianOceanBottlenose | TamanendBottlenose | unknown | dolphins | species | TP | 160 |
| IndianOceanBottlenose | AtlanticSpottedDolphin | unknown | dolphins | genera | NA | 161 |
| IndianOceanBottlenose | BlackSeaBottlenose | unknown | dolphins | species | TP | 162 |
| IndianOceanBottlenose | BridledDolphin | unknown | dolphins | genera | NA | 163 |
| IndianOceanBottlenose | FraserDolphin | unknown | dolphins | genera | NA | 164 |
| IndianOceanBottlenose | AustralasiaBottlenose | both sexes | dolphins | sub | TN | 165 |
| IndianOceanBottlenose | BurrunanBottlenose | both sexes | dolphins | sub | TN | 166 |

| | | | | | | |
|---|---|---|---|---|---|---|
| IndianOceanBottlenose | CommonDolphin | unknown | dolphins | genera | NA | 167 |
| IndianRhino | BlackRhino | neither sex | rhinos | genera | NA | 168 |
| Jaguar | EasternLion | unknown | panthera | species | TP | 169 |
| Jaguar | AmurLeopard | unknown | panthera | species | TP | 170 |
| Jaguar | BengalTiger | unknown | panthera | species | TP | 171 |
| Jaguar | AsiaticLion | unknown | panthera | species | TP | 172 |
| Jaguar | AmurTiger | unknown | panthera | species | TP | 173 |
| JapanBlack | AmericanBlack | unknown | ursids | species | TP | 174 |
| JapanBlack | AsiaticBlack | unknown | ursids | sub | FP | 175 |
| JapanBlack | BrownBear | unknown | ursids | species | TP | 176 |
| JavanRhino | IndianRhino | unknown | rhinos | species | TP | 177 |
| JavanRhino | BlackRhino | neither sex | rhinos | genera | NA | 178 |
| KamtchatkaBear | GrizzlyBear | both sexes | brownbears | sub | FP | 179 |
| KamtchatkaBear | HokkaidoBear | both sexes | brownbears | sub | TN | 180 |
| KamtchatkaBear | KodiakBear | both sexes | brownbears | sub | TN | 181 |
| KamtchatkaBear | SyrianBear | both sexes | brownbears | sub | FP | 182 |
| KansuRedDeer | TianshanWapiti | both sexes | cervus | sub | TN | 183 |
| KansuRedDeer | BactrianDeer | unknown | cervus | species | TP | 184 |
| KansuRedDeer | CaspianRedDeer | unknown | cervus | species | TP | 185 |
| KansuRedDeer | CentralEuropeanRedDeer | unknown | cervus | species | TP | 186 |
| Kiang | ChapmanZebra | unknown | equus | species | TP | 187 |
| Kiang | GrevyZebra | unknown | equus | species | TP | 188 |
| Kiang | DomesticDonkey | unknown | equus | species | TP | 189 |
| Kiang | AfricanWildass | unknown | equus | species | TP | 190 |
| Kiang | DomesticHorse | neither sex | equus | species | TP | 191 |
| KordofanGiraffe | MasaiGiraffe | unknown | giraffes | species | TP | 192 |
| KordofanGiraffe | SouthAfricanGiraffe | unknown | giraffes | species | TP | 193 |
| KordofanGiraffe | NubianGiraffe | both sexes | giraffes | sub | TN | 194 |
| KordofanGiraffe | ReticulatedGiraffe | unknown | giraffes | species | TP | 195 |
| Lama | BactrianCamel | neither sex | camelids | genera | NA | 196 |
| Lama | Dromedary | neither sex | camelids | genera | NA | 197 |
| Lama | Alpaca | unknown | camelids | species | TP | 198 |
| Lama | WildGuanaco | both sexes | camelids | sub | FP | 199 |
| LowlandAnoa | AmericanBison | neither sex | bovini | genera | NA | 200 |
| LowlandAnoa | CapeBuffalo | neither sex | bovini | genera | NA | 201 |
| LowlandAnoa | Gaur | neither sex | bovini | genera | NA | 202 |
| LowlandAnoa | EuropeanBison | neither sex | bovini | genera | NA | 203 |
| LowlandAnoa | Zebu | neither sex | bovini | genera | NA | 204 |
| LowlandAnoa | RiverBuffalo | neither sex | bovini | species | TP | 205 |
| LowlandAnoa | ForestBuffalo | neither sex | bovini | genera | NA | 206 |
| LuangwaGiraffe | ReticulatedGiraffe | unknown | giraffes | species | TP | 207 |
| LuangwaGiraffe | NubianGiraffe | unknown | giraffes | species | TP | 208 |
| LuangwaGiraffe | KordofanGiraffe | unknown | giraffes | species | TP | 209 |
| LuangwaGiraffe | AngolanGiraffe | both sexes | giraffes | species | TP | 210 |
| LuangwaGiraffe | SouthAfricanGiraffe | both sexes | giraffes | species | TP | 211 |
| LuangwaGiraffe | MasaiGiraffe | both sexes | giraffes | sub | TN | 212 |
| MasaiGiraffe | SouthAfricanGiraffe | unknown | giraffes | species | TP | 213 |
| MasaiGiraffe | ReticulatedGiraffe | genetic_isolation | giraffes | species | TP | 214 |
| MediterraneanBottlenose | BridledDolphin | unknown | dolphins | genera | NA | 215 |
| MediterraneanBottlenose | BlackSeaBottlenose | both sexes | dolphins | sub | TN | 216 |
| MediterraneanBottlenose | AtlanticSpottedDolphin | unknown | dolphins | genera | NA | 217 |
| MediterraneanBottlenose | BurrunanBottlenose | unknown | dolphins | species | TP | 218 |
| MediterraneanBottlenose | CommonDolphin | unknown | dolphins | genera | NA | 219 |
| MediterraneanBottlenose | TamanendBottlenose | unknown | dolphins | species | FN | 220 |
| MediterraneanBottlenose | FraserDolphin | unknown | dolphins | genera | NA | 221 |
| MediterraneanBottlenose | AustralasiaBottlenose | unknown | dolphins | species | TP | 222 |
| MediterraneanBottlenose | IndianOceanBottlenose | unknown | dolphins | species | TP | 223 |
| Moose | EuropeanRoeDeer | neither sex | capreolini | genera | NA | 224 |
| Moose | SiberianRoeDeer | neither sex | capreolini | genera | NA | 225 |
| MuleDeer | SitkaDeer | both sexes | odocoileus | sub | FP | 226 |
| NileBuffalo | CapeBuffalo | both sexes | bovini | sub | TN | 227 |

| | | | | | | |
|---|---|---|---|---|---|---|
| NileBuffalo | RiverBuffalo | neither sex | bovini | genera | NA | 228 |
| NileBuffalo | EuropeanBison | neither sex | bovini | genera | NA | 229 |
| NileBuffalo | Gaur | neither sex | bovini | genera | NA | 230 |
| NileBuffalo | ForestBuffalo | both sexes | bovini | sub | TN | 231 |
| NileBuffalo | AmericanBison | neither sex | bovini | genera | NA | 232 |
| NileBuffalo | LowlandAnoa | neither sex | bovini | genera | NA | 233 |
| NileBuffalo | Zebu | neither sex | bovini | genera | NA | 234 |
| NileBuffalo | Yak | neither sex | bovini | genera | NA | 235 |
| NorthAfricanLion | AmurTiger | females only | panthera | species | TP | 236 |
| NorthAfricanLion | EasternLion | both sexes | panthera | sub | TN | 237 |
| NorthAfricanLion | AsiaticLion | both sexes | panthera | sub | TN | 238 |
| NorthAfricanLion | BengalTiger | females only | panthera | species | TP | 239 |
| NorthAfricanLion | AmurLeopard | unknown | panthera | species | TP | 240 |
| NorthAfricanLion | Jaguar | unknown | panthera | species | TP | 241 |
| NorthAmericanWapiti | BactrianDeer | unknown | cervus | species | TP | 242 |
| NorthAmericanWapiti | KansuRedDeer | both sexes | cervus | sub | TN | 243 |
| NorthAmericanWapiti | CaspianRedDeer | unknown | cervus | species | TP | 244 |
| NorthAmericanWapiti | TianshanWapiti | both sexes | cervus | sub | TN | 245 |
| NorthAmericanWapiti | CentralEuropeanRedDeer | genetic_isolation | cervus | species | TP | 246 |
| NorthAtlanticBlueWhale | SeiWhale | neither sex | bluewhales | species | TP | 247 |
| NorthAtlanticBlueWhale | PacificBlueWhale | both sexes | bluewhales | sub | TN | 248 |
| NorthAtlanticBlueWhale | PygmyBlueWhale | both sexes | bluewhales | sub | TN | 249 |
| NorthAtlanticOrca | AlaskaResidentOrca | both sexes | killerwhales | sub | TN | 250 |
| NorthAtlanticOrca | TransientOrca | both sexes | killerwhales | sub | TN | 251 |
| NorthChinaBoar | CentralEuropeanBoar | both sexes | wildboar | sub | FP | 252 |
| NorthChinaBoar | AnatolianBoar | both sexes | wildboar | sub | FP | 253 |
| NorthernLynx | Bobcat | unknown | lynx | species | TP | 254 |
| NorthernLynx | CanadianLynx | unknown | lynx | species | TP | 255 |
| NorthernLynx | IberianLynx | unknown | lynx | species | TP | 256 |
| NorthernWhiteRhino | IndianRhino | neither sex | rhinos | genera | NA | 257 |
| NorthernWhiteRhino | JavanRhino | neither sex | rhinos | genera | NA | 258 |
| NorthernWhiteRhino | BlackRhino | unknown | rhinos | genera | NA | 259 |
| NorthernWhiteRhino | SumatranRhino | neither sex | rhinos | genera | NA | 260 |
| NubianGiraffe | MasaiGiraffe | unknown | giraffes | species | TP | 261 |
| NubianGiraffe | SouthAfricanGiraffe | unknown | giraffes | species | TP | 262 |
| NubianGiraffe | ReticulatedGiraffe | both sexes | giraffes | species | TP | 263 |
| OffshoreBottlenose | BridledDolphin | unknown | dolphins | genera | NA | 264 |
| OffshoreBottlenose | MediterraneanBottlenose | both sexes | dolphins | sub | TN | 265 |
| OffshoreBottlenose | BurrunanBottlenose | unknown | dolphins | species | TP | 266 |
| OffshoreBottlenose | BlackSeaBottlenose | both sexes | dolphins | sub | TN | 267 |
| OffshoreBottlenose | CommonDolphin | unknown | dolphins | genera | NA | 268 |
| OffshoreBottlenose | TamanendBottlenose | genetic_isolation | dolphins | species | FN | 269 |
| OffshoreBottlenose | AtlanticSpottedDolphin | unknown | dolphins | genera | NA | 270 |
| OffshoreBottlenose | FraserDolphin | unknown | dolphins | genera | NA | 271 |
| OffshoreBottlenose | AustralasiaBottlenose | genetic_isolation | dolphins | species | TP | 272 |
| OffshoreBottlenose | IndianOceanBottlenose | unknown | dolphins | species | TP | 273 |
| Okapi | ReticulatedGiraffe | neither sex | giraffes | genera | NA | 274 |
| Okapi | LuangwaGiraffe | neither sex | giraffes | genera | NA | 275 |
| Okapi | KordofanGiraffe | neither sex | giraffes | genera | NA | 276 |
| Okapi | AngolanGiraffe | neither sex | giraffes | genera | NA | 277 |
| Okapi | SouthAfricanGiraffe | neither sex | giraffes | genera | NA | 278 |
| Okapi | MasaiGiraffe | neither sex | giraffes | genera | NA | 279 |
| Okapi | NubianGiraffe | neither sex | giraffes | genera | NA | 280 |
| Okapi | WestAfricanGiraffe | neither sex | giraffes | genera | NA | 281 |
| Onager | Kiang | unknown | equus | sub | FP | 282 |
| Onager | DomesticDonkey | unknown | equus | species | TP | 283 |
| Onager | GrevyZebra | unknown | equus | species | TP | 284 |
| Onager | AfricanWildass | unknown | equus | species | TP | 285 |
| Onager | DomesticHorse | neither sex | equus | species | TP | 286 |
| Onager | ChapmanZebra | unknown | equus | species | TP | 287 |
| PacificBlueWhale | SeiWhale | neither sex | bluewhales | species | TP | 288 |

| | | | | | | |
|---|---|---|---|---|---|---|
| PacificOffshoreOrca | AntarcticOrca | both sexes | killerwhales | sub | TN | 289 |
| PacificOffshoreOrca | AlaskaResidentOrca | both sexes | killerwhales | sub | TN | 290 |
| PacificOffshoreOrca | TransientOrca | both sexes | killerwhales | sub | TN | 291 |
| PacificOffshoreOrca | NorthAtlanticOrca | both sexes | killerwhales | sub | TN | 292 |
| PolarBear | AmericanBlack | unknown | ursids | species | TP | 293 |
| PolarBear | BrownBear | genetic_isolation | ursids | species | TP | 294 |
| PolarBear | CaveBear | unknown | ursids | species | TP | 295 |
| PolarBear | JapanBlack | unknown | ursids | species | TP | 296 |
| PolarBear | AsiaticBlack | unknown | ursids | species | TP | 297 |
| PrzewalskiHorse | GrevyZebra | neither sex | equus | species | TP | 298 |
| PrzewalskiHorse | ChapmanZebra | neither sex | equus | species | TP | 299 |
| PrzewalskiHorse | Kiang | neither sex | equus | species | TP | 300 |
| PrzewalskiHorse | Onager | neither sex | equus | species | TP | 301 |
| PrzewalskiHorse | DomesticDonkey | neither sex | equus | species | TP | 302 |
| PrzewalskiHorse | AfricanWildass | neither sex | equus | species | TP | 303 |
| PrzewalskiHorse | DomesticHorse | both sexes | equus | sub | TN | 304 |
| PygmyBlueWhale | SeiWhale | neither sex | bluewhales | species | TP | 305 |
| PygmyBlueWhale | PacificBlueWhale | both sexes | bluewhales | sub | TN | 306 |
| ReticulatedGiraffe | SouthAfricanGiraffe | unknown | giraffes | species | TP | 307 |
| SiberianLynx | IberianLynx | unknown | lynx | species | TP | 308 |
| SiberianLynx | NorthernLynx | both sexes | lynx | sub | TN | 309 |
| SiberianLynx | Bobcat | unknown | lynx | species | TP | 310 |
| SiberianLynx | CanadianLynx | unknown | lynx | species | TP | 311 |
| SiberianRoeDeer | EuropeanRoeDeer | females only | capreolini | species | TP | 312 |
| SikaDeer | BactrianDeer | unknown | cervus | species | TP | 313 |
| SikaDeer | TianshanWapiti | unknown | cervus | species | TP | 314 |
| SikaDeer | CaspianRedDeer | unknown | cervus | species | TP | 315 |
| SikaDeer | NorthAmericanWapiti | unknown | cervus | species | TP | 316 |
| SikaDeer | CentralEuropeanRedDeer | genetic_isolation | cervus | species | TP | 317 |
| SikaDeer | KansuRedDeer | unknown | cervus | species | TP | 318 |
| SlothBear | AmericanBlack | unknown | ursids | genera | NA | 319 |
| SlothBear | CaveBear | unknown | ursids | genera | NA | 320 |
| SlothBear | BrownBear | unknown | ursids | genera | NA | 321 |
| SlothBear | AsiaticBlack | unknown | ursids | genera | NA | 322 |
| SlothBear | PolarBear | unknown | ursids | genera | NA | 323 |
| SlothBear | JapanBlack | unknown | ursids | genera | NA | 324 |
| SnowLeopard | AmurLeopard | unknown | panthera | species | TP | 325 |
| SnowLeopard | AsiaticLion | unknown | panthera | species | TP | 326 |
| SnowLeopard | Jaguar | unknown | panthera | species | TP | 327 |
| SnowLeopard | BengalTiger | unknown | panthera | species | TP | 328 |
| SnowLeopard | AmurTiger | unknown | panthera | species | TP | 329 |
| SnowLeopard | EasternLion | unknown | panthera | species | TP | 330 |
| SnowLeopard | NorthAfricanLion | unknown | panthera | species | TP | 331 |
| SouthAardwolf | BrownHyena | unknown | hyenas | genera | NA | 332 |
| SouthAardwolf | EastAardwolf | unknown | hyenas | sub | FP | 333 |
| SouthafricanLeopard | Jaguar | unknown | panthera | species | TP | 334 |
| SouthafricanLeopard | AmurLeopard | unknown | panthera | sub | FP | 335 |
| SouthafricanLeopard | SnowLeopard | unknown | panthera | species | TP | 336 |
| SouthafricanLeopard | BengalTiger | unknown | panthera | species | TP | 337 |
| SouthafricanLeopard | EasternLion | unknown | panthera | species | TP | 338 |
| SouthafricanLeopard | AmurTiger | unknown | panthera | species | TP | 339 |
| SouthafricanLeopard | NorthAfricanLion | unknown | panthera | species | TP | 340 |
| SouthafricanLeopard | AsiaticLion | unknown | panthera | species | TP | 341 |
| SouthBlueWildebeest | EastBlueWildebeest | both sexes | gnous | sub | TN | 342 |
| SouthBlueWildebeest | Hartebeest | neither sex | gnous | genera | NA | 343 |
| SouthBlueWildebeest | BlackWildebeest | genetic_isolation | gnous | species | TP | 344 |
| SouthChinaBoar | NorthChinaBoar | both sexes | wildboar | sub | TN | 345 |
| SouthChinaBoar | CentralEuropeanBoar | both sexes | wildboar | sub | FP | 346 |
| SouthChinaBoar | AnatolianBoar | both sexes | wildboar | sub | FP | 347 |
| SouthChinaTiger | SnowLeopard | unknown | panthera | species | TP | 348 |
| SouthChinaTiger | AmurLeopard | unknown | panthera | species | TP | 349 |

| | | | | | | |
|---|---|---|---|---|---|---|
| SouthChinaTiger | SouthafricanLeopard | unknown | panthera | species | TP | 350 |
| SouthChinaTiger | EasternLion | females only | panthera | species | TP | 351 |
| SouthChinaTiger | AsiaticLion | females only | panthera | species | TP | 352 |
| SouthChinaTiger | Jaguar | unknown | panthera | species | TP | 353 |
| SouthChinaTiger | AmurTiger | both sexes | panthera | sub | TN | 354 |
| SouthChinaTiger | BengalTiger | both sexes | panthera | sub | TN | 355 |
| SouthChinaTiger | NorthAfricanLion | females only | panthera | species | TP | 356 |
| SouthernLion | AmurLeopard | unknown | panthera | species | TP | 357 |
| SouthernLion | SouthChinaTiger | females only | panthera | species | TP | 358 |
| SouthernLion | AmurTiger | females only | panthera | species | TP | 359 |
| SouthernLion | SnowLeopard | unknown | panthera | species | TP | 360 |
| SouthernLion | NorthAfricanLion | both sexes | panthera | sub | TN | 361 |
| SouthernLion | SouthafricanLeopard | unknown | panthera | species | TP | 362 |
| SouthernLion | EasternLion | both sexes | panthera | sub | TN | 363 |
| SouthernLion | AsiaticLion | both sexes | panthera | sub | TN | 364 |
| SouthernLion | Jaguar | unknown | panthera | species | TP | 365 |
| SouthernLion | BengalTiger | females only | panthera | species | TP | 366 |
| SouthernResidentOrca | AlaskaResidentOrca | both sexes | killerwhales | sub | TN | 367 |
| SouthernResidentOrca | PacificOffshoreOrca | both sexes | killerwhales | sub | TN | 368 |
| SouthernResidentOrca | TransientOrca | both sexes | killerwhales | sub | TN | 369 |
| SouthernResidentOrca | AntarcticOrca | both sexes | killerwhales | sub | TN | 370 |
| SouthernResidentOrca | NorthAtlanticOrca | both sexes | killerwhales | sub | TN | 371 |
| SouthernWhiteRhino | IndianRhino | neither sex | rhinos | genera | NA | 372 |
| SouthernWhiteRhino | SumatranRhino | neither sex | rhinos | genera | NA | 373 |
| SouthernWhiteRhino | JavanRhino | neither sex | rhinos | genera | NA | 374 |
| SouthernWhiteRhino | BlackRhino | unknown | rhinos | genera | NA | 375 |
| SouthernWhiteRhino | NorthernWhiteRhino | both sexes | rhinos | sub | TN | 376 |
| SpectacledBear | PolarBear | neither sex | ursids | genera | NA | 377 |
| SpectacledBear | BrownBear | neither sex | ursids | genera | NA | 378 |
| SpectacledBear | SlothBear | neither sex | ursids | genera | NA | 379 |
| SpectacledBear | AmericanBlack | neither sex | ursids | genera | NA | 380 |
| SpectacledBear | CaveBear | neither sex | ursids | genera | NA | 381 |
| SpectacledBear | AsiaticBlack | neither sex | ursids | genera | NA | 382 |
| SpectacledBear | JapanBlack | neither sex | ursids | genera | NA | 383 |
| SpinnerDolphin | AustralasiaBottlenose | unknown | dolphins | genera | NA | 384 |
| SpinnerDolphin | TamanendBottlenose | unknown | dolphins | genera | NA | 385 |
| SpinnerDolphin | BurrunanBottlenose | unknown | dolphins | genera | NA | 386 |
| SpinnerDolphin | AtlanticSpottedDolphin | unknown | dolphins | species | TP | 387 |
| SpinnerDolphin | CommonDolphin | unknown | dolphins | genera | NA | 388 |
| SpinnerDolphin | BridledDolphin | unknown | dolphins | species | TP | 389 |
| SpinnerDolphin | BlackSeaBottlenose | unknown | dolphins | genera | NA | 390 |
| SpinnerDolphin | MediterraneanBottlenose | unknown | dolphins | genera | NA | 391 |
| SpinnerDolphin | OffshoreBottlenose | unknown | dolphins | genera | NA | 392 |
| SpinnerDolphin | IndianOceanBottlenose | unknown | dolphins | genera | NA | 393 |
| SpinnerDolphin | FraserDolphin | unknown | dolphins | genera | NA | 394 |
| SpottedHyena | SouthAardwolf | neither sex | hyenas | genera | NA | 395 |
| SpottedHyena | EastAardwolf | neither sex | hyenas | genera | NA | 396 |
| SpottedHyena | BrownHyena | unknown | hyenas | genera | NA | 397 |
| SriLankanElephant | ForestElephant | neither sex | elephants | genera | NA | 398 |
| SriLankanElephant | BorneanElephant | both sexes | elephants | sub | TN | 399 |
| SriLankanElephant | AsianElephant | both sexes | elephants | sub | TN | 400 |
| SriLankanElephant | SavannaElephant | neither sex | elephants | genera | NA | 401 |
| StraightTuskedElephant | BorneanElephant | unknown | elephants | genera | NA | 402 |
| StraightTuskedElephant | ForestElephant | unknown | elephants | genera | NA | 403 |
| StraightTuskedElephant | AsianElephant | unknown | elephants | genera | NA | 404 |
| StraightTuskedElephant | SriLankanElephant | unknown | elephants | genera | NA | 405 |
| StraightTuskedElephant | SavannaElephant | unknown | elephants | genera | NA | 406 |
| StripedDolphin | BridledDolphin | unknown | dolphins | species | TP | 407 |
| StripedDolphin | BurrunanBottlenose | unknown | dolphins | genera | NA | 408 |
| StripedDolphin | AustralasiaBottlenose | unknown | dolphins | genera | NA | 409 |
| StripedDolphin | SpinnerDolphin | unknown | dolphins | species | TP | 410 |

| | | | | | | |
|---|---|---|---|---|---|---|
| StripedDolphin | CommonDolphin | unknown | dolphins | genera | NA | 411 |
| StripedDolphin | TamanendBottlenose | unknown | dolphins | genera | NA | 412 |
| StripedDolphin | BlackSeaBottlenose | unknown | dolphins | genera | NA | 413 |
| StripedDolphin | MediterraneanBottlenose | unknown | dolphins | genera | NA | 414 |
| StripedDolphin | FraserDolphin | unknown | dolphins | genera | NA | 415 |
| StripedDolphin | OffshoreBottlenose | unknown | dolphins | genera | NA | 416 |
| StripedDolphin | IndianOceanBottlenose | unknown | dolphins | genera | NA | 417 |
| StripedDolphin | AtlanticSpottedDolphin | unknown | dolphins | species | TP | 418 |
| StripedHyena | BrownHyena | unknown | hyenas | genera | NA | 419 |
| StripedHyena | EastAardwolf | neither sex | hyenas | genera | NA | 420 |
| StripedHyena | SouthAardwolf | neither sex | hyenas | genera | NA | 421 |
| StripedHyena | SpottedHyena | neither sex | hyenas | genera | NA | 422 |
| SudanBuffalo | ForestBuffalo | both sexes | bovini | sub | TN | 423 |
| SudanBuffalo | EuropeanBison | neither sex | bovini | genera | NA | 424 |
| SudanBuffalo | RiverBuffalo | neither sex | bovini | genera | NA | 425 |
| SudanBuffalo | NileBuffalo | both sexes | bovini | sub | TN | 426 |
| SudanBuffalo | CapeBuffalo | both sexes | bovini | sub | TN | 427 |
| SudanBuffalo | Gaur | neither sex | bovini | genera | NA | 428 |
| SudanBuffalo | AmericanBison | neither sex | bovini | genera | NA | 429 |
| SudanBuffalo | Yak | neither sex | bovini | genera | NA | 430 |
| SudanBuffalo | Zebu | neither sex | bovini | genera | NA | 431 |
| SudanBuffalo | LowlandAnoa | neither sex | bovini | genera | NA | 432 |
| SumatranElephant | SriLankanElephant | both sexes | elephants | sub | TN | 433 |
| SumatranElephant | ForestElephant | neither sex | elephants | genera | NA | 434 |
| SumatranElephant | StraightTuskedElephant | unknown | elephants | genera | NA | 435 |
| SumatranElephant | AsianElephant | both sexes | elephants | sub | TN | 436 |
| SumatranElephant | SavannaElephant | neither sex | elephants | genera | NA | 437 |
| SumatranElephant | BorneanElephant | both sexes | elephants | sub | TN | 438 |
| SumatranOrang | BorneanOrang | unknown | orangutan | species | TP | 439 |
| SumatranRhino | IndianRhino | neither sex | rhinos | genera | NA | 440 |
| SumatranRhino | BlackRhino | neither sex | rhinos | genera | NA | 441 |
| SumatranTiger | SouthafricanLeopard | unknown | panthera | species | TP | 442 |
| SumatranTiger | EasternLion | females only | panthera | species | TP | 443 |
| SumatranTiger | AmurTiger | both sexes | panthera | sub | TN | 444 |
| SumatranTiger | Jaguar | unknown | panthera | species | TP | 445 |
| SumatranTiger | SouthChinaTiger | both sexes | panthera | sub | TN | 446 |
| SumatranTiger | SouthernLion | females only | panthera | species | TP | 447 |
| SumatranTiger | BengalTiger | both sexes | panthera | sub | TN | 448 |
| SumatranTiger | AsiaticLion | females only | panthera | species | TP | 449 |
| SumatranTiger | AmurLeopard | unknown | panthera | species | TP | 450 |
| SumatranTiger | NorthAfricanLion | females only | panthera | species | TP | 451 |
| SumatranTiger | SnowLeopard | unknown | panthera | species | TP | 452 |
| SunBear | PolarBear | unknown | ursids | genera | NA | 453 |
| SunBear | SlothBear | unknown | ursids | genera | NA | 454 |
| SunBear | CaveBear | unknown | ursids | genera | NA | 455 |
| SunBear | AmericanBlack | unknown | ursids | genera | NA | 456 |
| SunBear | SpectacledBear | neither sex | ursids | genera | NA | 457 |
| SunBear | AsiaticBlack | unknown | ursids | genera | NA | 458 |
| SunBear | JapanBlack | unknown | ursids | genera | NA | 459 |
| SunBear | BrownBear | unknown | ursids | genera | NA | 460 |
| SwampBuffalo | RiverBuffalo | genetic_isolation | bovini | species | TP | 461 |
| SwampBuffalo | CapeBuffalo | neither sex | bovini | genera | NA | 462 |
| SwampBuffalo | Zebu | neither sex | bovini | genera | NA | 463 |
| SwampBuffalo | ForestBuffalo | neither sex | bovini | genera | NA | 464 |
| SwampBuffalo | Yak | neither sex | bovini | genera | NA | 465 |
| SwampBuffalo | EuropeanBison | neither sex | bovini | genera | NA | 466 |
| SwampBuffalo | Gaur | neither sex | bovini | genera | NA | 467 |
| SwampBuffalo | AmericanBison | neither sex | bovini | genera | NA | 468 |
| SwampBuffalo | SudanBuffalo | neither sex | bovini | genera | NA | 469 |
| SwampBuffalo | NileBuffalo | neither sex | bovini | genera | NA | 470 |
| SwampBuffalo | LowlandAnoa | neither sex | bovini | species | TP | 471 |

| | | | | | | |
|---|---|---|---|---|---|---|
| SyrianBear | KodiakBear | both sexes | brownbears | sub | FP | 472 |
| SyrianBear | GrizzlyBear | both sexes | brownbears | sub | FP | 473 |
| TapanuliOrang | BorneanOrang | unknown | orangutan | species | TP | 474 |
| TapanuliOrang | SumatranOrang | unknown | orangutan | species | FN | 475 |
| ThinhornSheep | BighornSheep | unknown | sheep | species | FN | 476 |
| ThoroldsDeer | BactrianDeer | unknown | cervus | species | TP | 477 |
| ThoroldsDeer | SikaDeer | unknown | cervus | species | TP | 478 |
| ThoroldsDeer | TianshanWapiti | unknown | cervus | species | TP | 479 |
| ThoroldsDeer | KansuRedDeer | unknown | cervus | species | TP | 480 |
| ThoroldsDeer | CaspianRedDeer | unknown | cervus | species | TP | 481 |
| ThoroldsDeer | NorthAmericanWapiti | unknown | cervus | species | TP | 482 |
| ThoroldsDeer | CentralEuropeanRedDeer | unknown | cervus | species | TP | 483 |
| ThoroldsDeer | IberianRedDeer | unknown | cervus | species | TP | 484 |
| ThoroldsDeer | YarkandDeer | unknown | cervus | species | TP | 485 |
| TianshanRoeDeer | IberianRoeDeer | females only | capreolini | species | TP | 486 |
| TianshanRoeDeer | EuropeanRoeDeer | females only | capreolini | species | TP | 487 |
| TianshanRoeDeer | Moose | neither sex | capreolini | genera | NA | 488 |
| TianshanRoeDeer | SiberianRoeDeer | both sexes | capreolini | sub | TN | 489 |
| TransientOrca | AlaskaResidentOrca | genetic_isolation | killerwhales | sub | TN | 490 |
| VirungaGorilla | EasternLowlandGorilla | both sexes | gorillas | sub | TN | 491 |
| VirungaGorilla | BwindiGorilla | both sexes | gorillas | sub | TN | 492 |
| VirungaGorilla | CrossRiverGorilla | unknown | gorillas | species | TP | 493 |
| Waterdeer | TianshanRoeDeer | unknown | capreolini | genera | NA | 494 |
| Waterdeer | SiberianRoeDeer | unknown | capreolini | genera | NA | 495 |
| Waterdeer | Moose | neither sex | capreolini | genera | NA | 496 |
| Waterdeer | EuropeanRoeDeer | unknown | capreolini | genera | NA | 497 |
| Waterdeer | IberianRoeDeer | unknown | capreolini | genera | NA | 498 |
| WestAfricanGiraffe | KordofanGiraffe | both sexes | giraffes | sub | TN | 499 |
| WestAfricanGiraffe | SouthAfricanGiraffe | unknown | giraffes | species | TP | 500 |
| WestAfricanGiraffe | AngolanGiraffe | unknown | giraffes | species | TP | 501 |
| WestAfricanGiraffe | LuangwaGiraffe | unknown | giraffes | species | TP | 502 |
| WestAfricanGiraffe | NubianGiraffe | both sexes | giraffes | sub | TN | 503 |
| WestAfricanGiraffe | ReticulatedGiraffe | unknown | giraffes | species | TP | 504 |
| WestAfricanGiraffe | MasaiGiraffe | unknown | giraffes | species | TP | 505 |
| WestBlueWildebeest | EastBlueWildebeest | both sexes | gnous | sub | TN | 506 |
| WestBlueWildebeest | BlackWildebeest | unknown | gnous | species | TP | 507 |
| WestBlueWildebeest | Hartebeest | neither sex | gnous | genera | NA | 508 |
| WestBlueWildebeest | SouthBlueWildebeest | both sexes | gnous | sub | TN | 509 |
| WesternLion | AmurTiger | females only | panthera | species | TP | 510 |
| WesternLion | SouthernLion | both sexes | panthera | sub | TN | 511 |
| WesternLion | NorthAfricanLion | both sexes | panthera | sub | TN | 512 |
| WesternLion | SouthafricanLeopard | unknown | panthera | species | TP | 513 |
| WesternLion | AsiaticLion | both sexes | panthera | sub | TN | 514 |
| WesternLion | Jaguar | unknown | panthera | species | TP | 515 |
| WesternLion | EasternLion | both sexes | panthera | sub | TN | 516 |
| WesternLion | SouthChinaTiger | females only | panthera | species | TP | 517 |
| WesternLion | SnowLeopard | unknown | panthera | species | TP | 518 |
| WesternLion | AmurLeopard | unknown | panthera | species | TP | 519 |
| WesternLion | SumatranTiger | females only | panthera | species | TP | 520 |
| WesternLion | BengalTiger | females only | panthera | species | TP | 521 |
| WesternLowlandGorilla | VirungaGorilla | unknown | gorillas | species | TP | 522 |
| WesternLowlandGorilla | EasternLowlandGorilla | unknown | gorillas | species | TP | 523 |
| WesternLowlandGorilla | CrossRiverGorilla | both sexes | gorillas | sub | TN | 524 |
| WesternLowlandGorilla | BwindiGorilla | unknown | gorillas | species | TP | 525 |
| WhitetailedDeer | SitkaDeer | females only | odocoileus | species | TP | 526 |
| WhitetailedDeer | MuleDeer | females only | odocoileus | species | TP | 527 |
| WildCamel | BactrianCamel | both sexes | camelids | sub | TN | 528 |
| WildCamel | Lama | neither sex | camelids | genera | NA | 529 |
| WildCamel | Dromedary | genetic_isolation | camelids | species | TP | 530 |
| WildCamel | WildVicuna | neither sex | camelids | genera | NA | 531 |
| WildCamel | Alpaca | neither sex | camelids | genera | NA | 532 |

| | | | | | | |
|---|---|---|---|---|---|---|
| WildCamel | WildGuanaco | neither sex | camelids | genera | NA | 533 |
| WildGuanaco | BactrianCamel | neither sex | camelids | genera | NA | 534 |
| WildGuanaco | Alpaca | unknown | camelids | sub | FP | 535 |
| WildGuanaco | Dromedary | neither sex | camelids | genera | NA | 536 |
| WildVicuna | BactrianCamel | neither sex | camelids | genera | NA | 537 |
| WildVicuna | Lama | unknown | camelids | species | TP | 538 |
| WildVicuna | Dromedary | neither sex | camelids | genera | NA | 539 |
| WildVicuna | WildGuanaco | genetic_isolation | camelids | species | TP | 540 |
| WildVicuna | Alpaca | unknown | camelids | sub | FP | 541 |
| WoollyMammoth | AsianElephant | unknown | elephants | genera | NA | 542 |
| WoollyMammoth | ForestElephant | unknown | elephants | genera | NA | 543 |
| WoollyMammoth | StraightTuskedElephant | unknown | elephants | genera | NA | 544 |
| WoollyMammoth | SumatranElephant | unknown | elephants | genera | NA | 545 |
| WoollyMammoth | SriLankanElephant | unknown | elephants | genera | NA | 546 |
| WoollyMammoth | BorneanElephant | unknown | elephants | genera | NA | 547 |
| WoollyMammoth | SavannaElephant | unknown | elephants | genera | NA | 548 |
| Yak | RiverBuffalo | neither sex | bovini | genera | NA | 549 |
| Yak | Zebu | females only | bovini | species | TP | 550 |
| Yak | LowlandAnoa | neither sex | bovini | genera | NA | 551 |
| Yak | EuropeanBison | unknown | bovini | species | TP | 552 |
| Yak | AmericanBison | females only | bovini | species | TP | 553 |
| Yak | Gaur | unknown | bovini | species | TP | 554 |
| Yak | ForestBuffalo | neither sex | bovini | genera | NA | 555 |
| Yak | CapeBuffalo | neither sex | bovini | genera | NA | 556 |
| YarkandDeer | TianshanWapiti | unknown | cervus | species | TP | 557 |
| YarkandDeer | BactrianDeer | both sexes | cervus | sub | TN | 558 |
| YarkandDeer | CentralEuropeanRedDeer | unknown | cervus | species | TP | 559 |
| YarkandDeer | NorthAmericanWapiti | unknown | cervus | species | TP | 560 |
| YarkandDeer | IberianRedDeer | unknown | cervus | species | TP | 561 |
| YarkandDeer | SikaDeer | unknown | cervus | species | TP | 562 |
| YarkandDeer | KansuRedDeer | unknown | cervus | species | TP | 563 |
| YarkandDeer | CaspianRedDeer | unknown | cervus | species | TP | 564 |
| Zebu | RiverBuffalo | neither sex | bovini | genera | NA | 565 |
| Zebu | Gaur | females only | bovini | species | TP | 566 |
| Zebu | AmericanBison | females only | bovini | species | TP | 567 |
| Zebu | CapeBuffalo | neither sex | bovini | genera | NA | 568 |
| Zebu | ForestBuffalo | neither sex | bovini | genera | NA | 569 |
| Zebu | EuropeanBison | females only | bovini | species | TP | 570 |

**Table S3. Conversion between *Fst* and the genealogical divergence index (*GDI*)**

| **Genealogical Divergence Index (*GDI*)** | **Probability of concordance (*P*)** | **Coalescent units ($\tau$ = T/2Ne)** | ***Fst*** |
|---|---|---|---|
| $GDI = (3P-1)/2$ | $P = 1 - (2/3)e^{-\tau}$ | $\tau = ln(1-Fst_1)$**, or<br>$\tau = Fst_2/(1-Fst_2)$*** | $Fst = (D_{xy}-\pi_{xy})/D_{xy}$ |
| 0.2* | 0.47 | 0.23 | $Fst_1 = 0.205$<br>$Fst_2 = 0.19$ |
| 0.7# | 0.8 | 1.2 | $Fst_1 = 0.7$<br>$Fst_2 = 0.55$ |

*GDI = 0.2 and 0.7 are values suggested in a previous study as putative thresholds for intraspecific and interspecific comparisons.

**The relationship between $\tau$ and $Fst_1$ holds in a demographic scenario in which the population split occurred so recently that mutations are negligible, such that differentiation is caused solely by loss of genetic variation owing to a reduction in population size.

***The relationship between $\tau$ and $Fst_2$ holds in a demographic scenario in which population sizes remain constant, such that differentiation is caused solely accumulation of novel mutations.

**Table S4. Comparison between study findings.** Comparison between distance estimates (*Dxy*, in percentages) reported by Lebedev et al 2025 (exome data) and by this study (genome-wide data). This data underlies supplementary figure 4B.

| lineage1 | lineage2 | latin1 | latin2 | lebedev_2025 | this_study |
|---|---|---|---|---|---|
| wolf | dog | lupus | familiaris | 0.095 | 0.18 |
| river_type | swamp_type | bubalis | carabanensis | 0.21 | 0.33 |
| river_type | lowland_anoa | bubalis | depressicornis | 0.27 | 0.39 |
| swamp_type | lowland_anoa | carabanensis | depressicornis | 0.23 | 0.35 |
| wapiti | asian_red | canadensis | hanglu | 0.13 | 0.24 |
| european_red | wapiti | elaphus | canadensis | 0.16 | 0.31 |
| european_red | asian_red | elaphus | hanglu | 0.16 | 0.28 |
| european_roe | siberian_roe | capreolus | pygargus | 0.37 | 0.55 |
| brown_bear | polar_bear | arctos | maritimus | 0.16 | 0.3 |
| brown_bear | american_bear | arctos | americanus | 0.265 | 0.51 |
| polar_bear | american_bear | maritimus | americanus | 0.265 | 0.51 |
| bactrian_camel | wild_camel | bactrianus | ferus | 0.09 | 0.12 |
| bactrian_camel | dromedary | bactrianus | dromedarius | 0.245 | 0.25 |

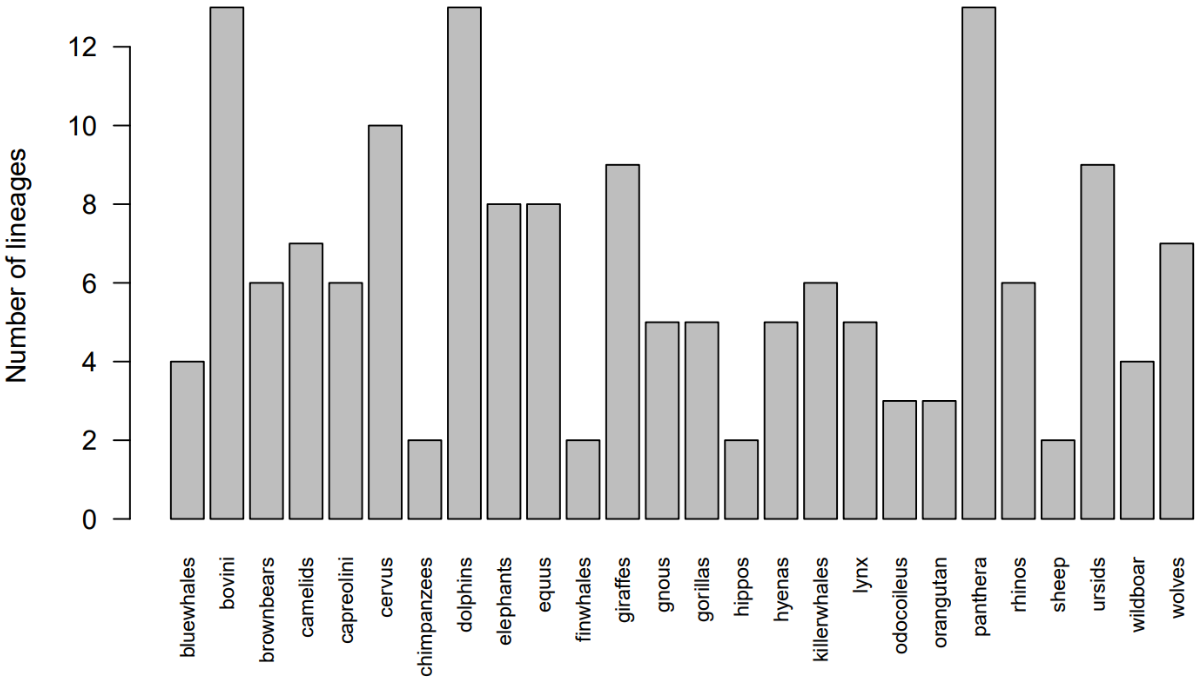


**Supplementary Figure 1A**. ***Overview of datasets***. Number of lineages per dataset, with each lineage being represented by at minimum two individuals. Genetic distances were calculated between all pairs of individuals within each of the datasets.

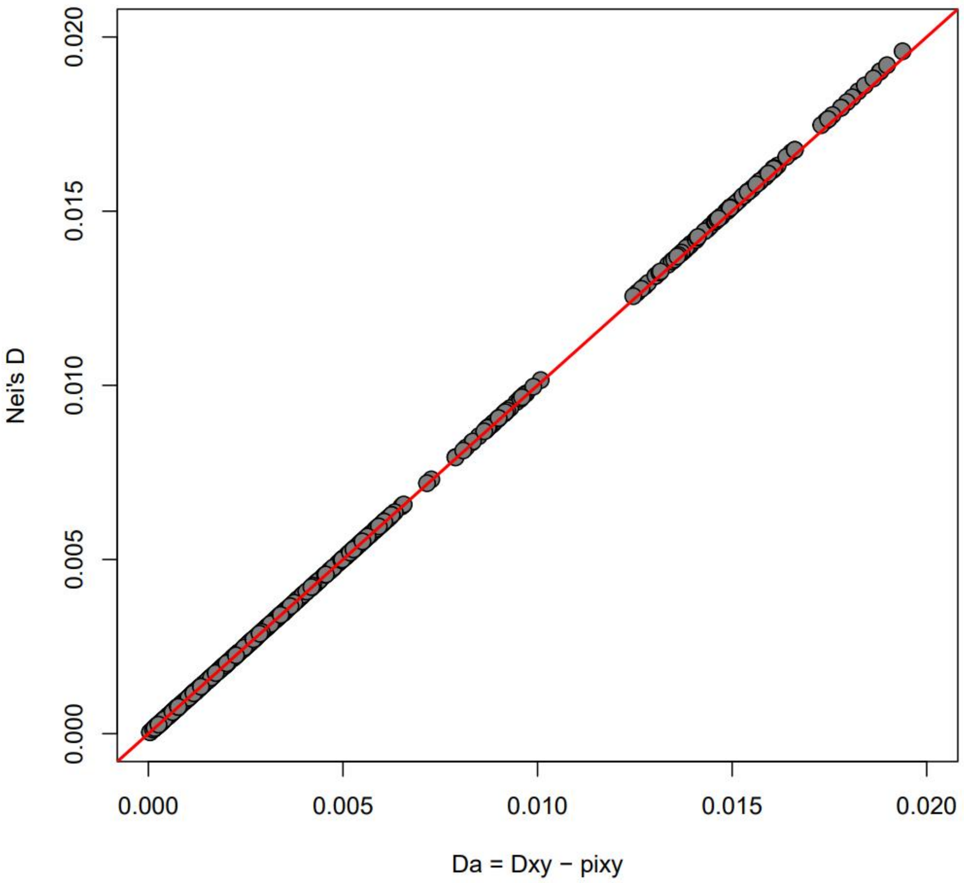


**Supplementary Figure 1B**. ***Net divergence.*** When including polymorphic and monomorphic sites, such that values are relatively small, Nei's D approximates net divergence ($Da = Dxy - \frac{1}{2}(\pi 1+\pi 2)$), here illustrated for a dataset of nearly 500 lineage pairs. Nei's D was calculated as: $D = -ln(Dxy/\sqrt{((1-\pi_1)*(1-\pi_2))})$

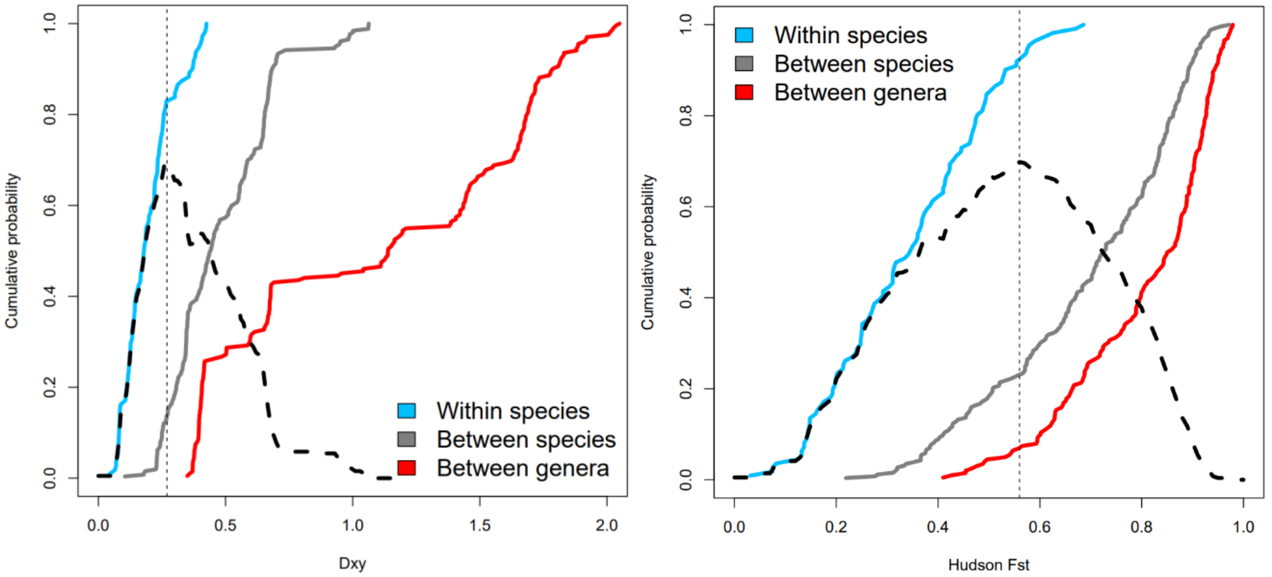


**Supplementary Figure 2A**. ***Cumulative distribution of Dxy and Hudson Fst-estimates across three categories***. These plots are made after figure 6 of Hey & Pinto (2012). The dashed curve depicts the difference in cumulative probability of *Dxy*-estimates (left) and Hudson *Fst*-estimates (right) among intraspecific ('within species') and interspecific ('between species') lineage pairs. The dashed vertical lines indicate the maximum difference, observed at *Dxy* = 0.26 (left) and *Fst* = 0.56 (right).

| | within species | between species | between genera |
|---|---|---|---|
| both sexes | 111 | 92 | 0 |
| females only | 0 | 35 | 0 |
| neither sex | 0 | 21 | 114 |

**Supplementary Figure 2B**. ***Contingency table listing the overlap between the two categorical variables (taxonomic level and hybrid fertility)***. Table listing the number of lineage pairs per category for two categorical variables. Not included are lineage pairs for which the viability or fertility of hybrids is unknown.

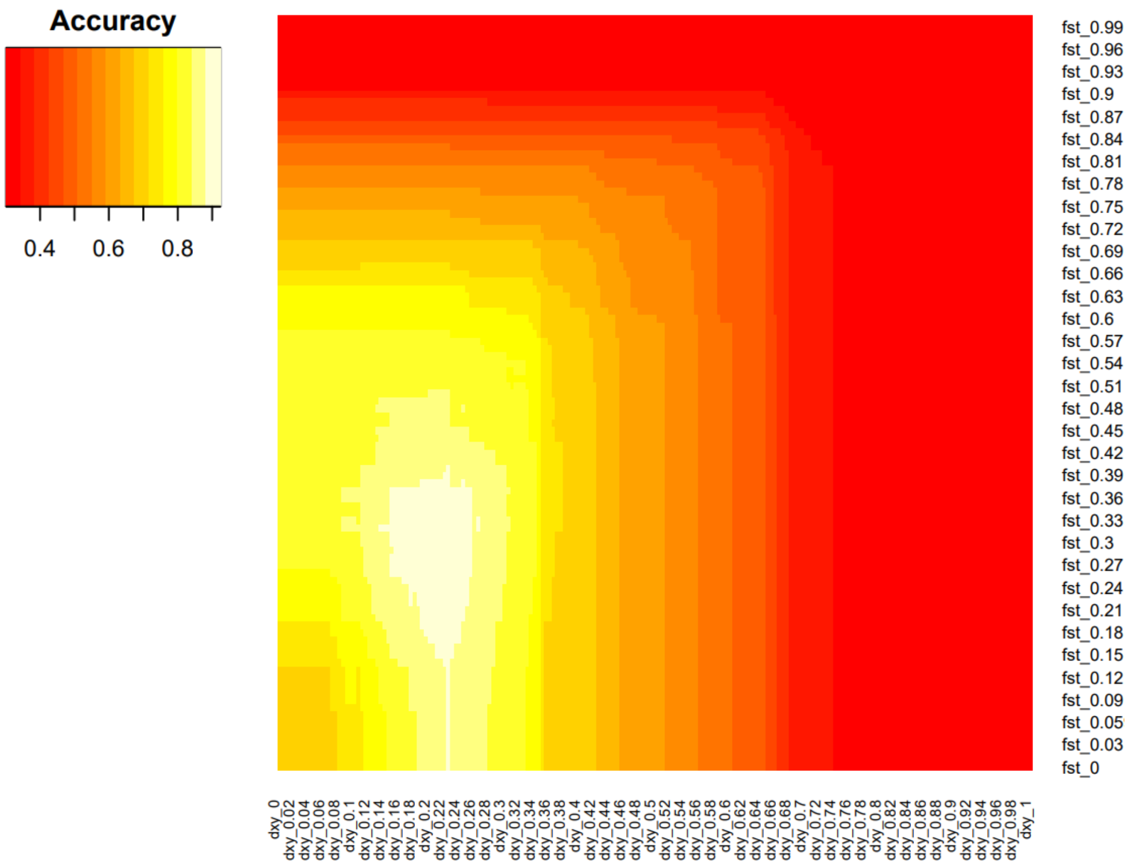


**Supplementary Figure 3A.** ***Accuracy heatmap for a range of combinations of Dxy and Fst-thresholds.*** The highest accuracy (brightest colour) is observed for Dxy = 0.225% and Fst = 0.26. This heatmap has been generated using the following R script, expecting Table S2 as input:

```
findthresholds<-function(inputdf=dxydf,fstvec=seq(0.5,0.1,-0.005),dxyvec=seq(0.1,0.5,0.005),doplot=TRUE)
	{
	# CALCULATE PERFORMANCE OF ALL CLASSIFICATION MODELS:
	alldf				<- inputdf[(inputdf$level!="intergeneric"),,drop=TRUE]
	accuracymat			<- matrix(NA,nrow=length(fstvec),ncol=length(dxyvec))
	rownames(accuracymat)		<- paste("fst",fstvec,sep="_")
	colnames(accuracymat)		<- paste("dxy",dxyvec,sep="_")
	for(i in c(1:length(fstvec)))
		{
		myfst			<- fstvec[i]
		for(j in c(1:length(dxyvec)))
			{
			mydxy	<- dxyvec[j]
			FP<- nrow(alldf[(alldf$HudsonFst<myfst|alldf$Dxy<mydxy)&alldf$level=="interspecific",])
			TP<- nrow(alldf[(alldf$HudsonFst>=myfst&alldf$Dxy>=mydxy)&alldf$level=="interspecific",])
			FN<- nrow(alldf[(alldf$HudsonFst>=myfst&alldf$Dxy>=mydxy)&alldf$level=="intraspecific",])
			TN<- nrow(alldf[(alldf$HudsonFst<myfst|alldf$Dxy<mydxy)&alldf$level=="intraspecific",])
			accuracymat[i,j]<- (TP+TN)/(TP+TN+FP+FN)
			}
		}
	library("gplots")
	heatmap.2(accuracymat,density.info="none",trace="none",dendrogram='none',Rowv=FALSE,Colv=FALSE)
	}
```

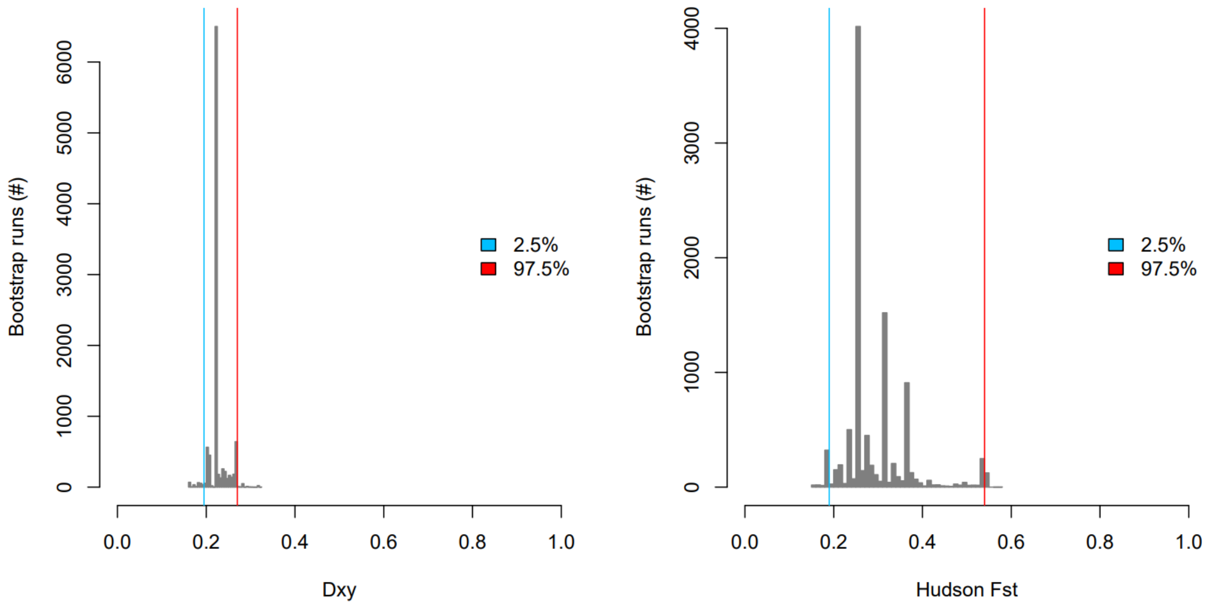


**Supplementary Figure 3B**. ***Outcome of bootstrap analyses.*** Histograms depicting the best performing Dxy and Fst thresholds after resampling of genera with replacement, a procedure which was repeated 10.000 times. The blue and red lines indicate the 95% confidence intervals: *Dxy* = 0.2% – 0.27%, and *Fst* = 0.19 – 0.55. These plots have been generated using the following R script:

```
bootstraprun<-function(nruns=1000,printverbose=10,fst_vec=seq(0.1,0.7,0.01),dxy_vec=seq(0.15,0.4,0.005))
	{
	fulldf		<- dxydf[dxydf$level!="intergeneric",,drop=TRUE]
	allgenera	<- unique(fulldf$genus1)
	ngenera		<- length(allgenera)
	cvdf		<<- data.frame("ndata"=rep(NA,nruns),"ngenera"=rep(NA,nruns),"fst"=rep(NA,nruns), "dxy"=rep(NA,nruns),
			"FP"=rep(NA,nruns),"TP"=rep(NA,nruns),"FN"=rep(NA,nruns),"TN"=rep(NA,nruns))
	for(k in c(1:nruns))
		{
		if(k%%printverbose==0){cat(paste("Run ",k,sep=""),sep="\n")}
		samplevec		<- sample(c(1:ngenera),ngenera,replace=TRUE)
		nunique			<- length(unique(samplevec))
		for(j in c(1:ngenera))
			{
			genusnr		<- samplevec[j]
			mygenus		<- allgenera[genusnr]
			interdf		<- fulldf[fulldf$genus1%in%mygenus&fulldf$level=="interspecific",]
			intradf		<- fulldf[fulldf$genus1%in%mygenus&fulldf$level=="intraspecific",]
			if(j==1)
				{
				bootstrapdf<- rbind(interdf,intradf)
				}else{
				bootstrapdf<- rbind(bootstrapdf,interdf,intradf)
				}
			}
		ndata			<- nrow(bootstrapdf)
		findthresholds(inputdf=bootstrapdf,fstvec=fst_vec,dxyvec=dxy_vec,doplot=FALSE)
		cvdf[k,]	<<- c(ndata,nunique,as.vector(unlist(outdf[1,])))
		}
	cvdf$accuracy		<<- round((cvdf$TP+cvdf$TN)/(cvdf$TP+cvdf$TN+cvdf$FP+cvdf$FN),4)
	cvdf$oob_error		<<- round((cvdf$FP+cvdf$FN)/(cvdf$TP+cvdf$TN+cvdf$FP+cvdf$FN),4)
	cvdf$sensitivity	<<- round(cvdf$TP/(cvdf$TP+cvdf$FN),4)
	cvdf$specificity	<<- round(cvdf$TN/(cvdf$TN+cvdf$FP),4)
	quantile(cvdf$dxy,c(0.025,0.975))
	quantile(cvdf$fst,c(0.025,0.975))
	hist(cvdf$dxy,breaks=40,col="grey50",border="grey50",xlim=c(0,1),xlab="Dxy",main=NULL,ylab="Bootstrap runs (#)")
	hist(cvdf$fst,breaks=40,col="grey50",border="grey50",xlim=c(0,1),xlab="Hudson Fst",main=NULL,ylab="Bootstrap runs(#)")
	}
```

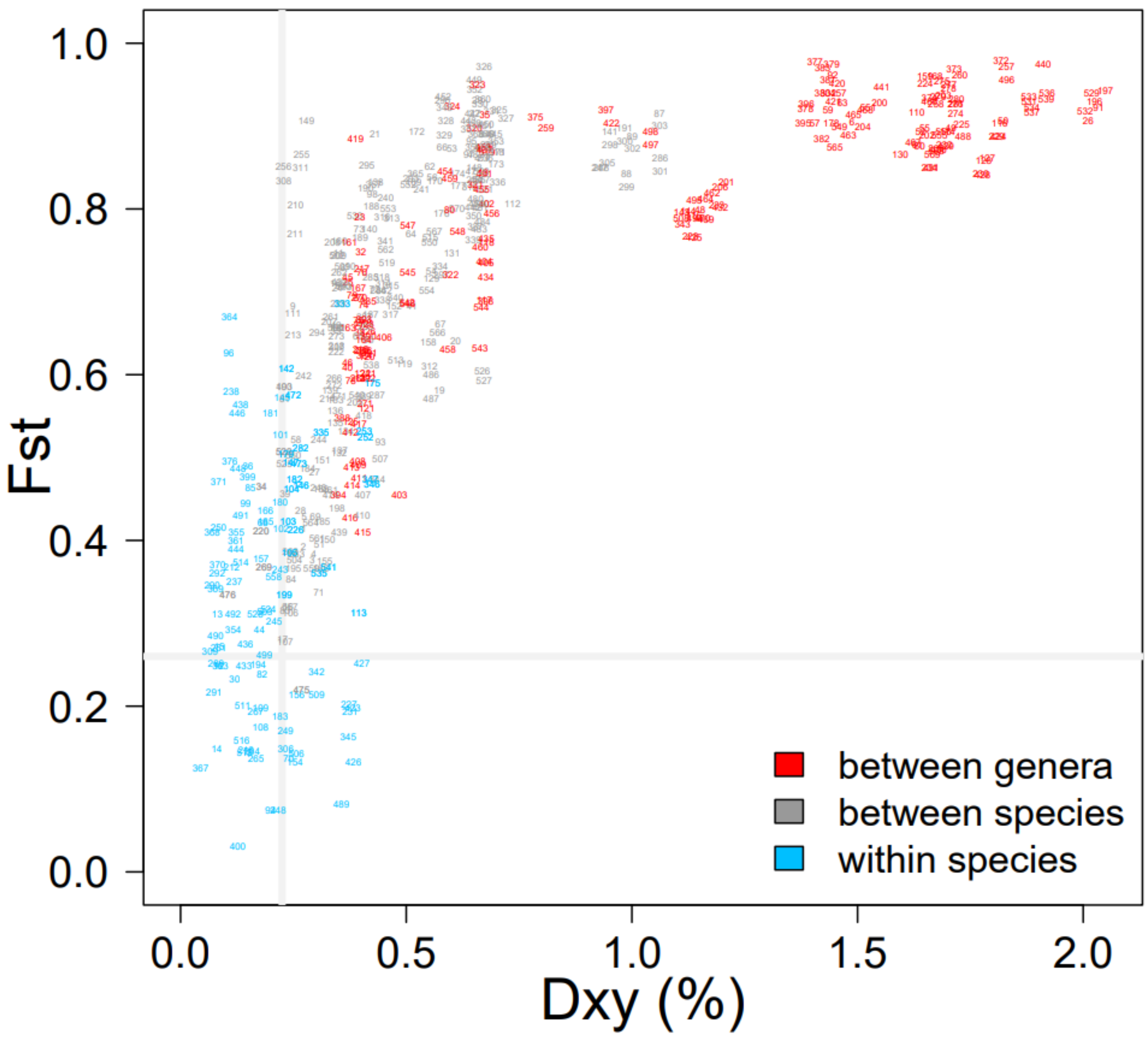


**Supplementary Figure 4A.** ***Fst-Dxy scatterplot with labels***. Idem as Fig, 2C, but with labels, which correspond to the index column in Table S2. Notable lineage pairs are Sumatran and Tapanuli orangutans (475), eastern coyote and Eurasian/Arctic gray wolves (107, 17), straight-tusked elephants and forest/savanna elephants (403, 406), Amur leopards and South African leopards (335), Asian versus European wild boars (252, 253, 346, 347), Japanese black bears and mainland Asiatic black bears (175), eastern and southeastern aardwolves (333), woolly mammoths versus Asian elephants (542, 543, 545, 547), common hippos and pygmy hippos (80), and brown hyenas and striped hyenas (419). The large cluster of intergeneric lineage pairs at Dxy ≈ 0.4% contains comparisons among dolphin lineages. The large cluster of intergeneric lineage pairs at Dxy ≈ 0.7% contains comparisons among bear lineages, as well as among Asian and African elephants. The cluster of intrageneric lineage pairs at Dxy ≈ 1.0% contains comparisons between assess/zebras and caballine horses.

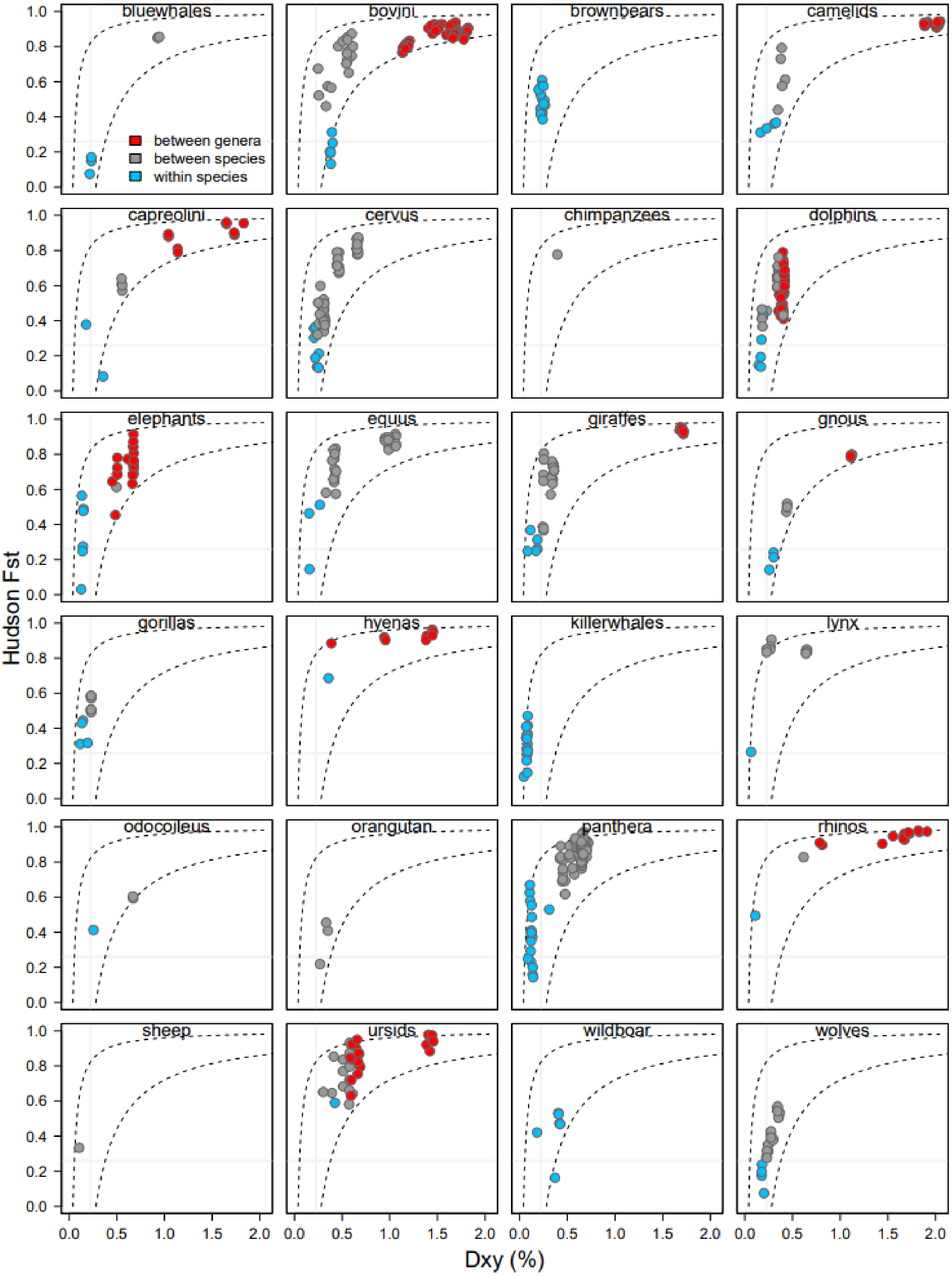


**Supplementary Figure 4B**. ***Fst-Dxy scatterplot per dataset***. Idem as Fig, 2C, but split out per dataset. Not shown are the datasets of fin whales and hippos (only one lineage pair).

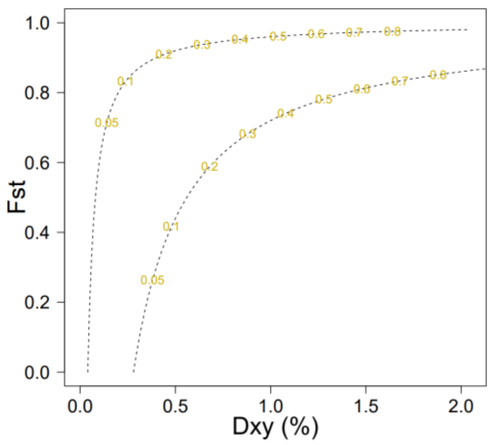


**Supplementary Figure 4C. *Predicted Fst vs Dxy curves.*** Expected *Dxy*- and *Fst*-estimates given a constant *Ne* of 10.000 (left) or 70.000 (right) and given a split time (in millions of generations) depicted in yellow. These curves roughly delineate the 'envelope' of expected empirical values, and can be reproduced using the following R commands:

```
u            <- 10^-8        # mutation rate per site per generation
Ne1          <- 10000        # low Ne
Ne2          <- 70000        # high Ne
t1           <- 5000         # t1, unit of time steps, in generations
tmax         <- 1000000      # maximum number of generations
xmax         <- 2*u*tmax
tvec         <- seq(0,tmax,t1)
pixy1        <- 4*Ne1*u
dxyvec1      <- seq(0,xmax,2*u*t1)+pixy1
fstvec1      <- (dxyvec1-pixy1)/dxyvec1
pixy2        <- 4*Ne3*u
dxyvec2      <- seq(0,xmax,2*u*t1)+pixy2
fstvec2      <- (dxyvec2-pixy2)/dxyvec2
pdf("Fst_vs_Dxy_prediction.pdf")
plot(dxydf$Dxy,dxydf$HudsonFst,ylim=c(0,1),xlim=c(0,max(dxydf$Dxy,na.rm=TRUE)),pch=16,col="white",xla
b="",ylab="",cex=1.5,cex.axis=1.5,las=1)
lines(dxyvec1*100,fstvec1,col="black",lty=2)
lines(dxyvec2*100,fstvec2,col="black",lty=2)
myvec    <- c(50000,seq(100000,800000,100000))
for(k in c(1:length(myvec)))
        {
        mytime  <-myvec[k]
        text(x=dxyvec1[tvec==mytime]*100,y=fstvec1[tvec==mytime],mytime/1000000,cex=1,col="gold3")
        text(x=dxyvec2[tvec==mytime]*100,y=fstvec2[tvec==mytime],mytime/1000000,cex=1,col="gold3")
        }
mtext(side=1,"Dxy (%)",cex=2,line=2.5)
mtext(side=2,"Fst",cex=2,line=2.5)
dev.off()
```

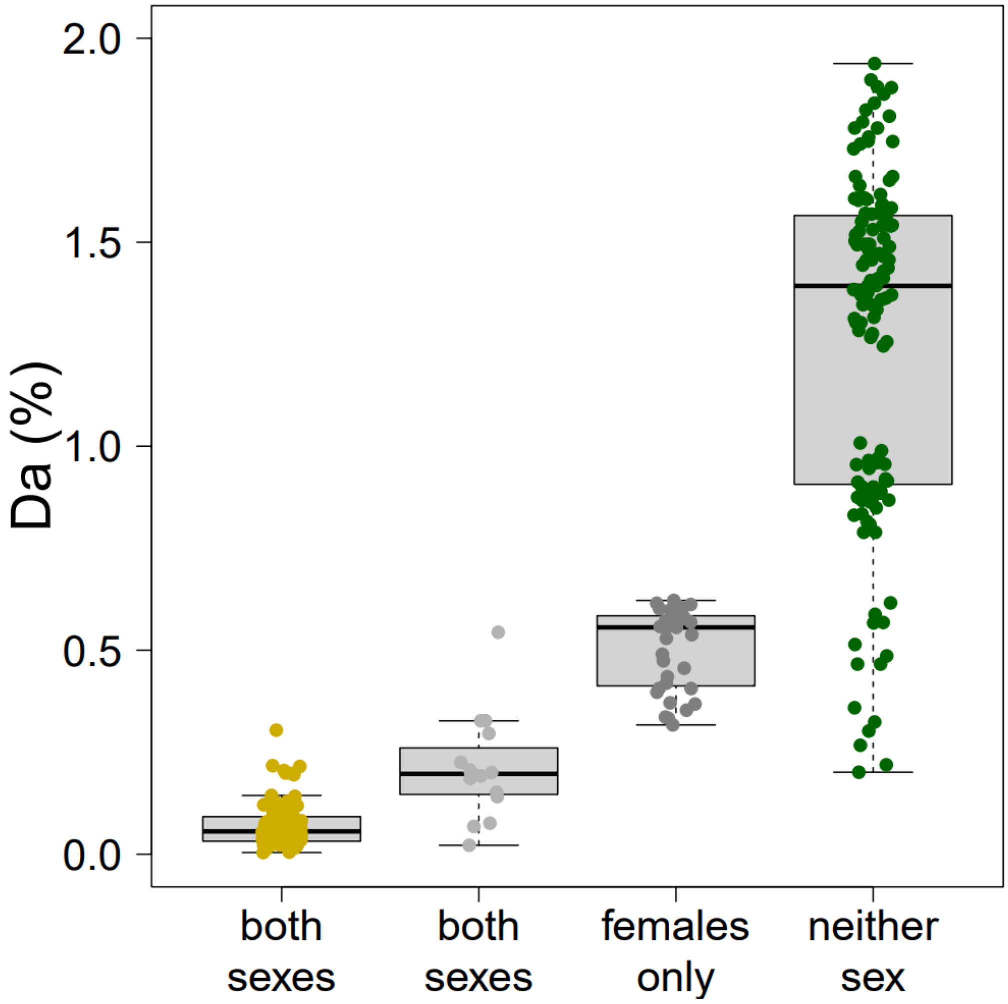


**Supplementary Figure 5.** ***Da per hybridisation class***. Boxplot depicting absolute genetic distances between lineage pairs, with lineage pairs categorized based on hybrid viability and fertility, with either both hybrid sexes being fertile (gold), genetic isolation at range overlap despite either hybrid sex being fertile (light grey), only female hybrids being fertile (grey) or neither hybrid sex being fertile or even viable (green). In other words, this plot is the same as main figure 2D, except for the addition of the extra category of genetic isolation. Lineage pairs included in this category are: coyote vs grey wolf, black vs blue wildebeest, brown bear vs polar bear, bobcat vs Canadian Lynx, red deer vs sika deer, red deer vs wapiti, reticulated giraffe vs Masai giraffe, Chapman zebra vs Grevy zebra, river-type vs swamp-type buffalo, guanaco vs vicuna, common vs Indo-Pacific bottlenose dolphin, and dromedary vs camel.

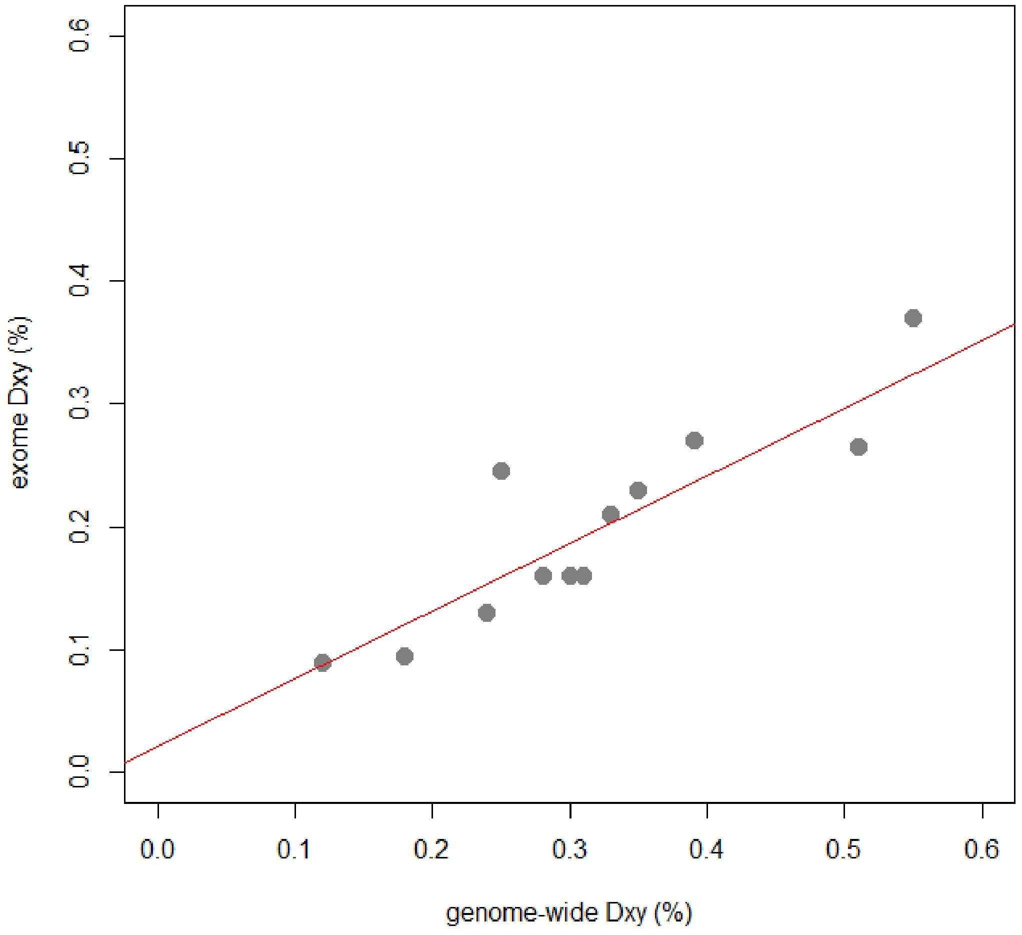


**Supplementary Figure 6.** Comparison between distance estimates of Lebedev et al. 2025 (exome data, y-axis) and distance estimates of this study (genome-wide data, x-axis) for overlapping lineage pairs. The intercept and coefficient are 0.02 and 0.55, respectively.

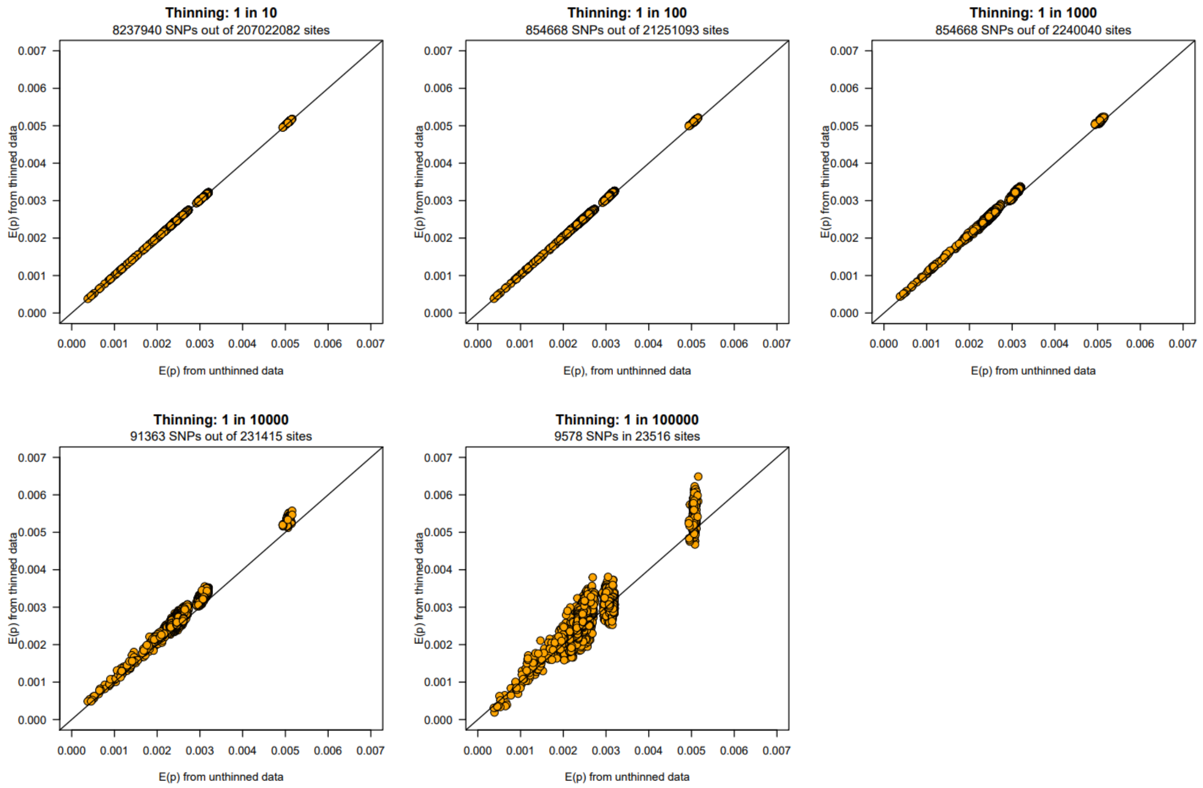


**Supplementary Figure 7A. *Thinning sensitivity analyses*.** Scatterplots depicting the sensitivity of *E(p)*-estimates to the thinning parameter, evaluated for a dataset of red deer, *Cervus elaphus* (De Jong et al. 2025). Each datapoint represents E(p)-estimates for a pair of individuals, inferred either from unthinned (x-axis) or thinned data (y-axis). In order to speed up calculations, we thinned datasets optionally by a factor of 1 in 100.

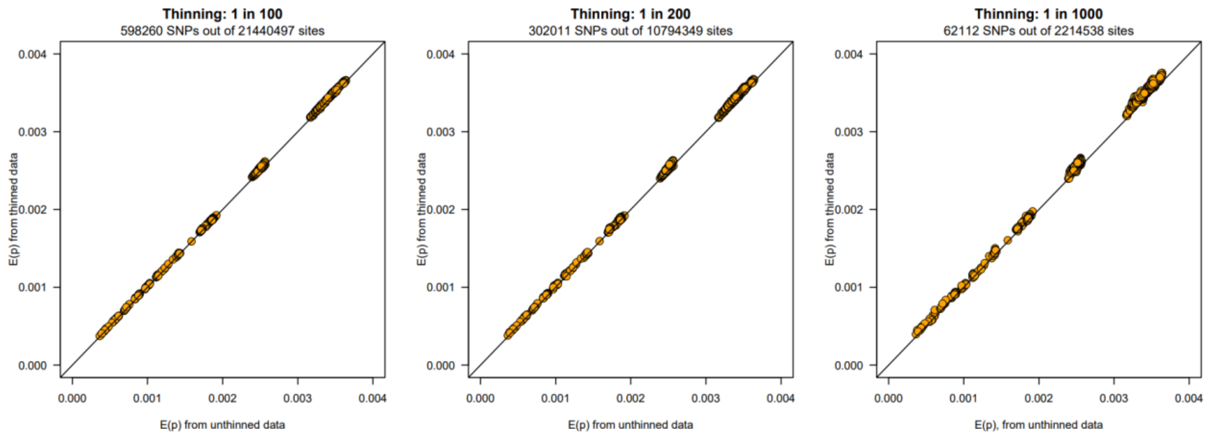


**Supplementary Figure 7B**. ***Thinning sensitivity analyses***. Scatterplots depicting the sensitivity of *E(p)*-estimates to thinning parameter, evaluated for a dataset of giraffes (Coimbra et al. 2023). Each datapoint represents *E(p)*-estimates for a pair of individuals, inferred either from unthinned (x-axis) or thinned data (y-axis).

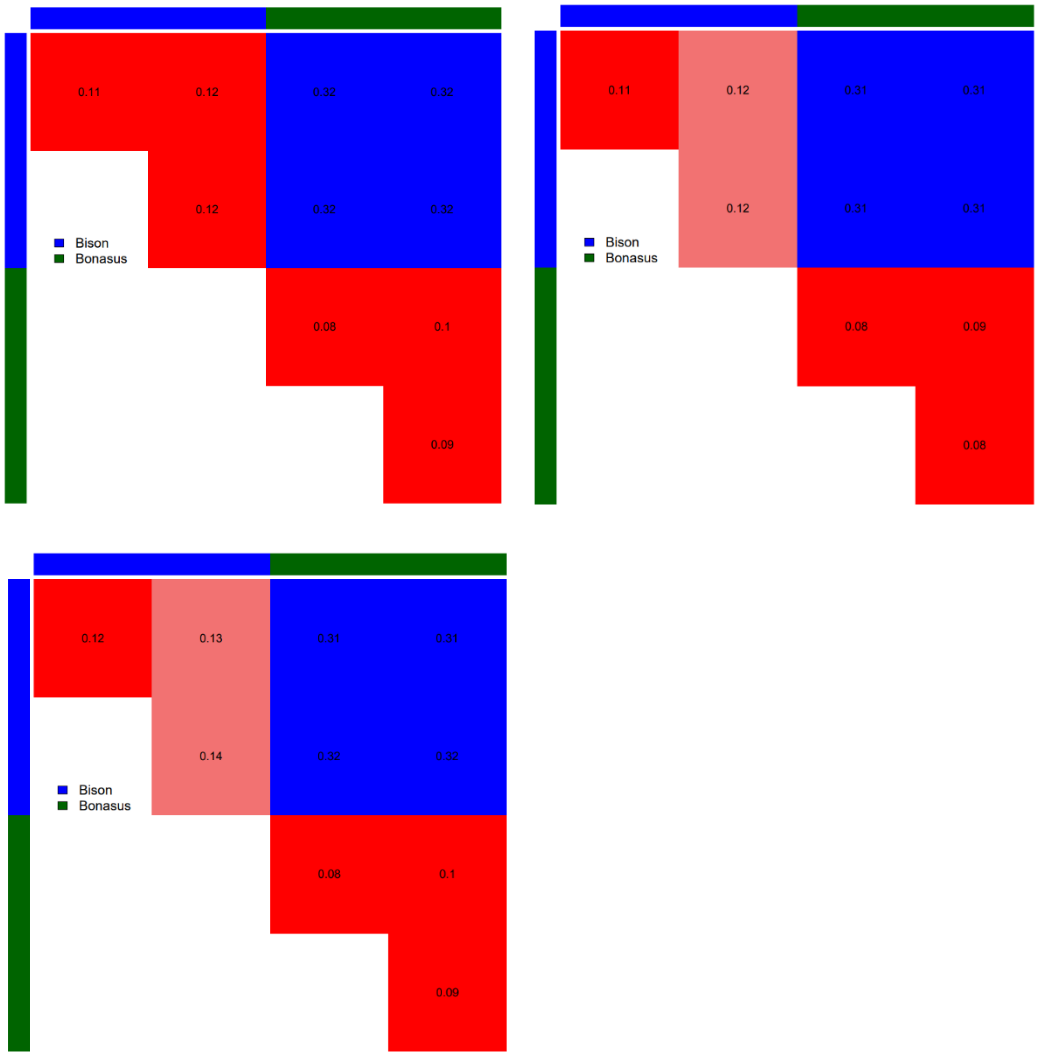


**Figure 7C. *Reference genome sensitivity analyses*.** *Dxy*- and *π*-estimates for American bison (*Bos bison*) and wisent (*Bos bonasus*) inferred from mapping sequencing reads against three different reference genomes, namely *B. bison* (top-left), *B. bonasus* (top-right) and *B. taurus* (bottom-left). Note the consistency of the obtained estimates.

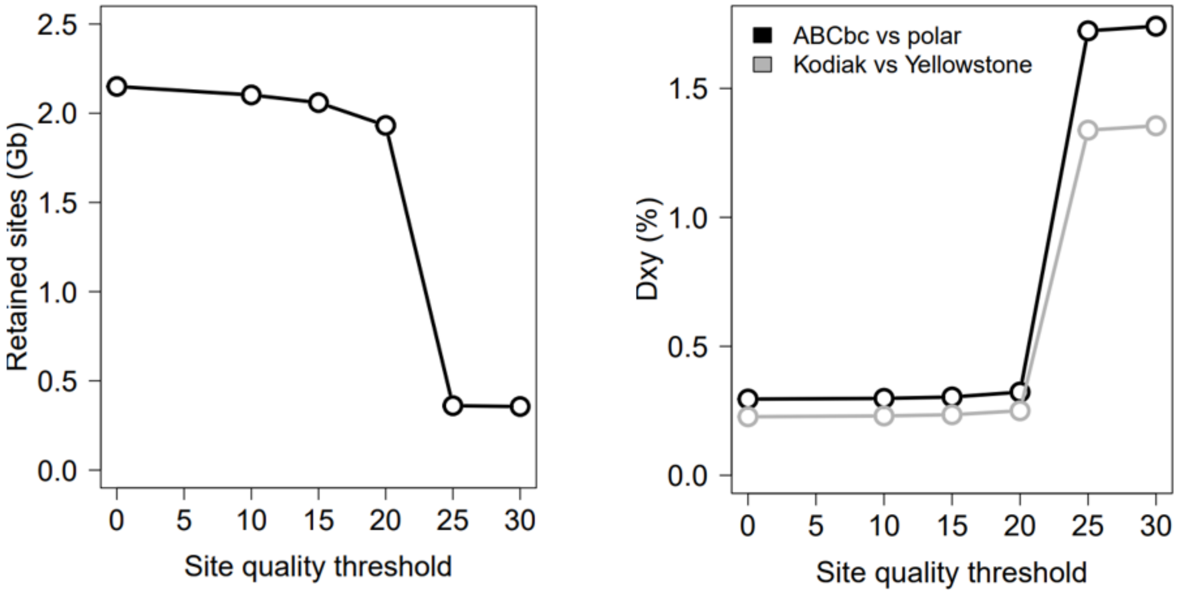


**Supplementary Figure 7D**. ***Site quality sensitivity analyses***. Scatterplots depicting the sensitivity of *E(p)*-estimates to the site quality filter, evaluated for a dataset of brown bears and polar bears (De Jong et al. 2025). Note that a site quality filter causes an upward bias of the E(p)-estimates, particularly for when the threshold is above 20. We decided to not apply a site quality filter.

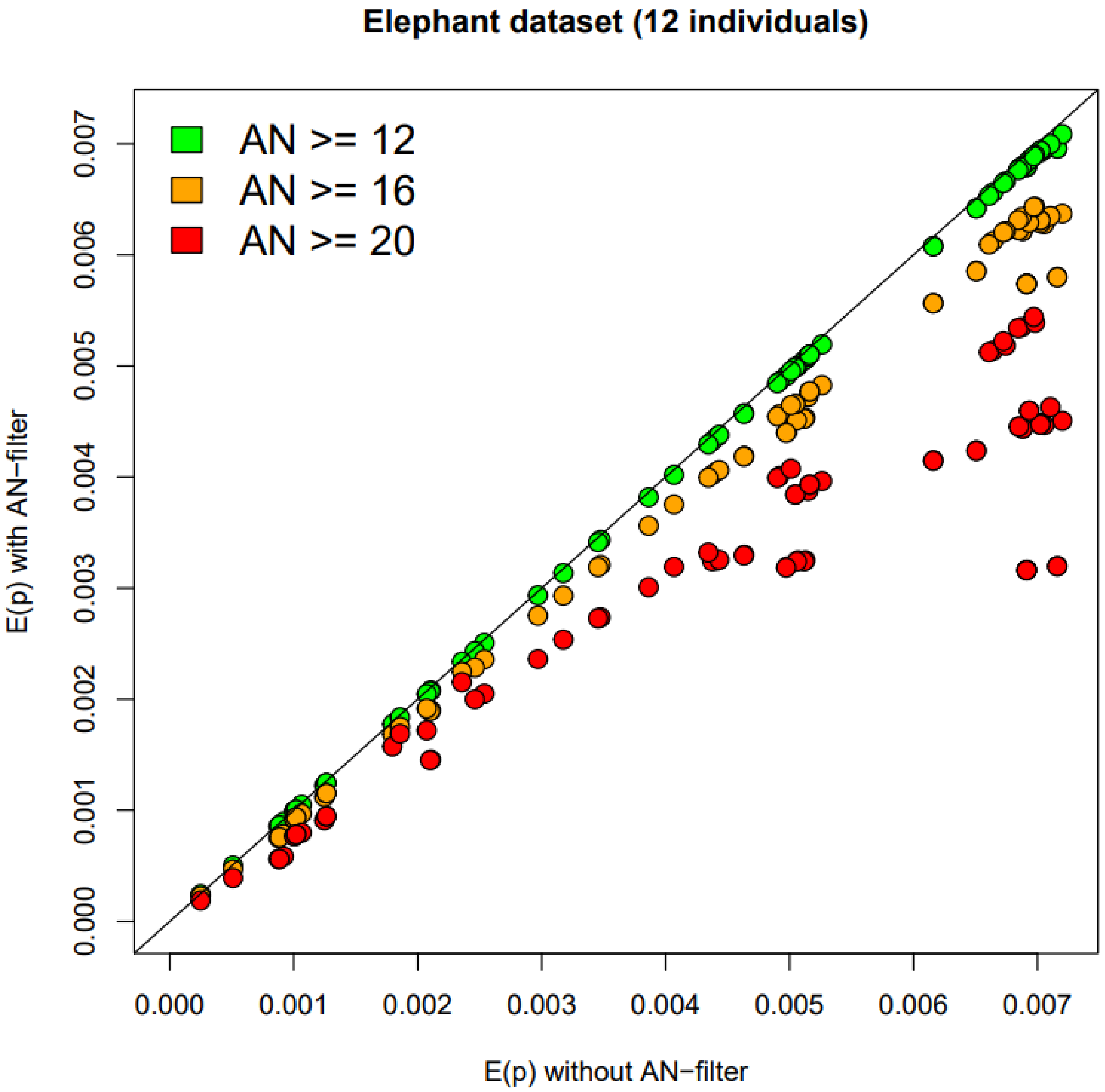


**Supplementary Figure 7E.** ***Effect of AN filter***. Scatterplots depicting the sensitivity of *E(p)*-estimates to a filter on per-site allelic number (AN), here evaluated for a dataset of 12 elephants (of which 3 ancient samples). Each datapoint represents *E(p)*-estimates for a pair of individuals, inferred either from unfiltered data (x-axis) or filtered data (y-axis), with colour coding denoting the exact filter setting. Note that a strict AN-filter causes *E(p)*-estimates to decrease, with the magnitude of decrease varying across pairs of individuals. We decided to not apply an AN-filter, and instead only filter sites on global depth.

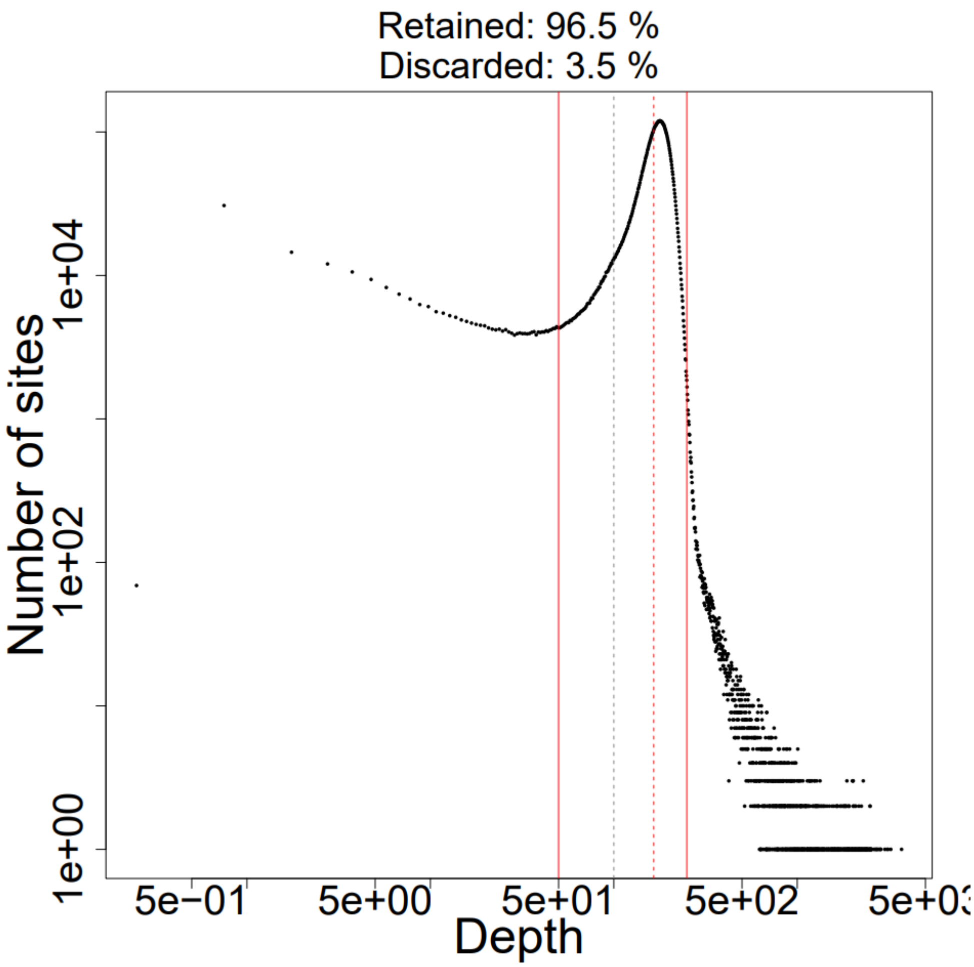


**Supplementary Figure 7F**. ***Depth filter***. Sites were removed based on a combined filter, based on visual examination of a histogram depicting the depth distribution across sites, here shown for a dataset of 10 gnous. The red solid lines denote the minimum and maximum depth thresholds.

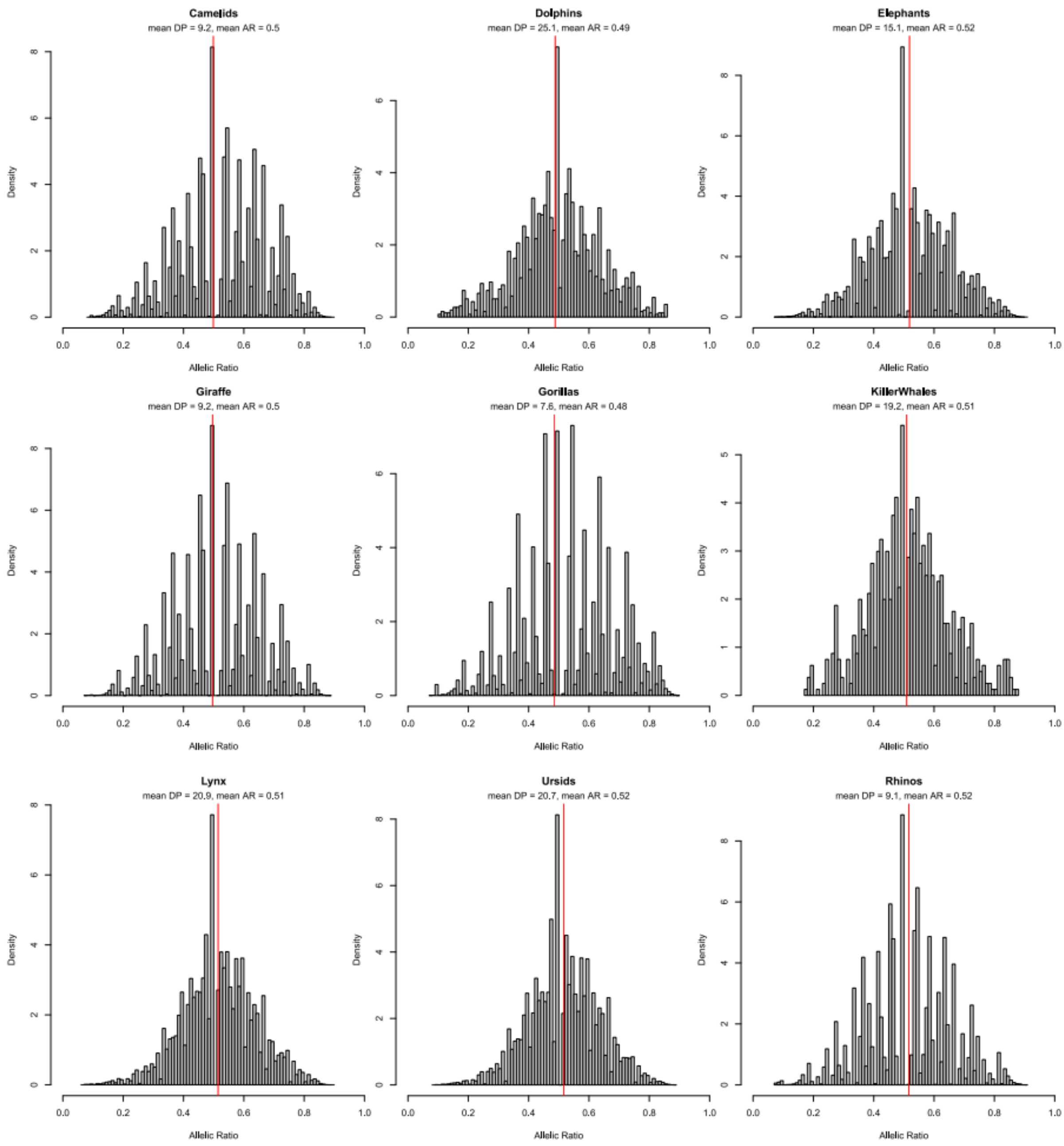


**Supplementary Figure 8**. ***Allelic ratio***. Histograms depicting allelic ratios observed across heterozygous genotypes for a set of nine datasets (all individuals combined). The allelic ratio was calculated as the proportion of reads per heterozygous genotype supporting one allele over the other. The mean expected value is 0.5.

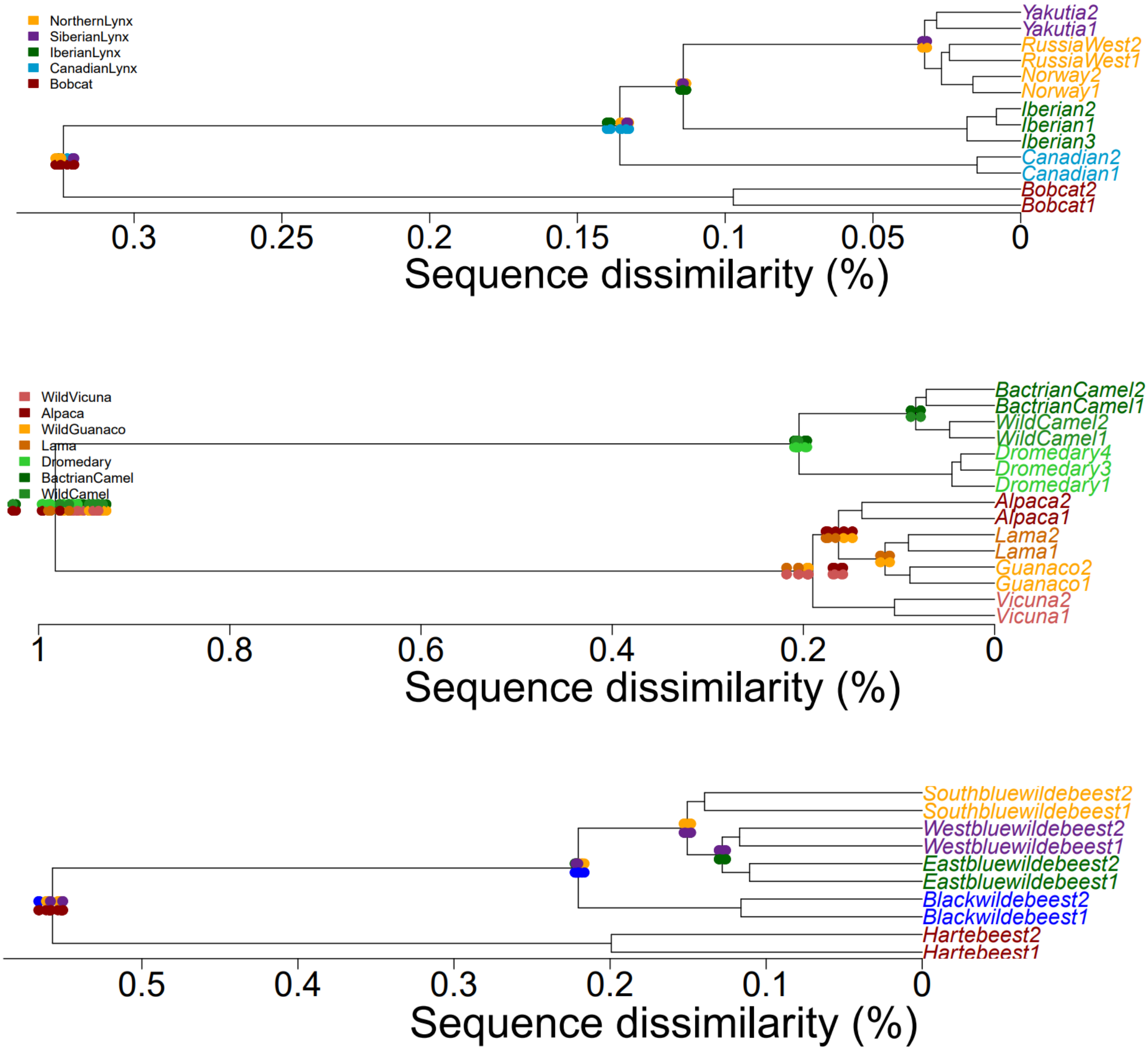


**Supplementary Figure 9.** ***UPGMA-dendrogram depicting genetic distances***. Genetic distances (sequence dissimilarity, in percentages) between pairs of individuals, superimposed on a UPGMA-dendrogram, for an arbitrary selection of data sets. Top: lynx; middle: camelids; bottom: gnus). Each pair of individuals is represented by a vertically aligned pair of dots, with the upper and lower dot according to the position in the dendrogram. Note that the values on the x-axis denote branch lengths, and hence are half the actual genetic distances between a pair of individuals. In the absence of gene flow and the absence of data artefacts (such as sequencing errors, genotype calling errors and incorrect distance calculations), genetic distances for any node are expected to be equidistant regardless of sample selection.

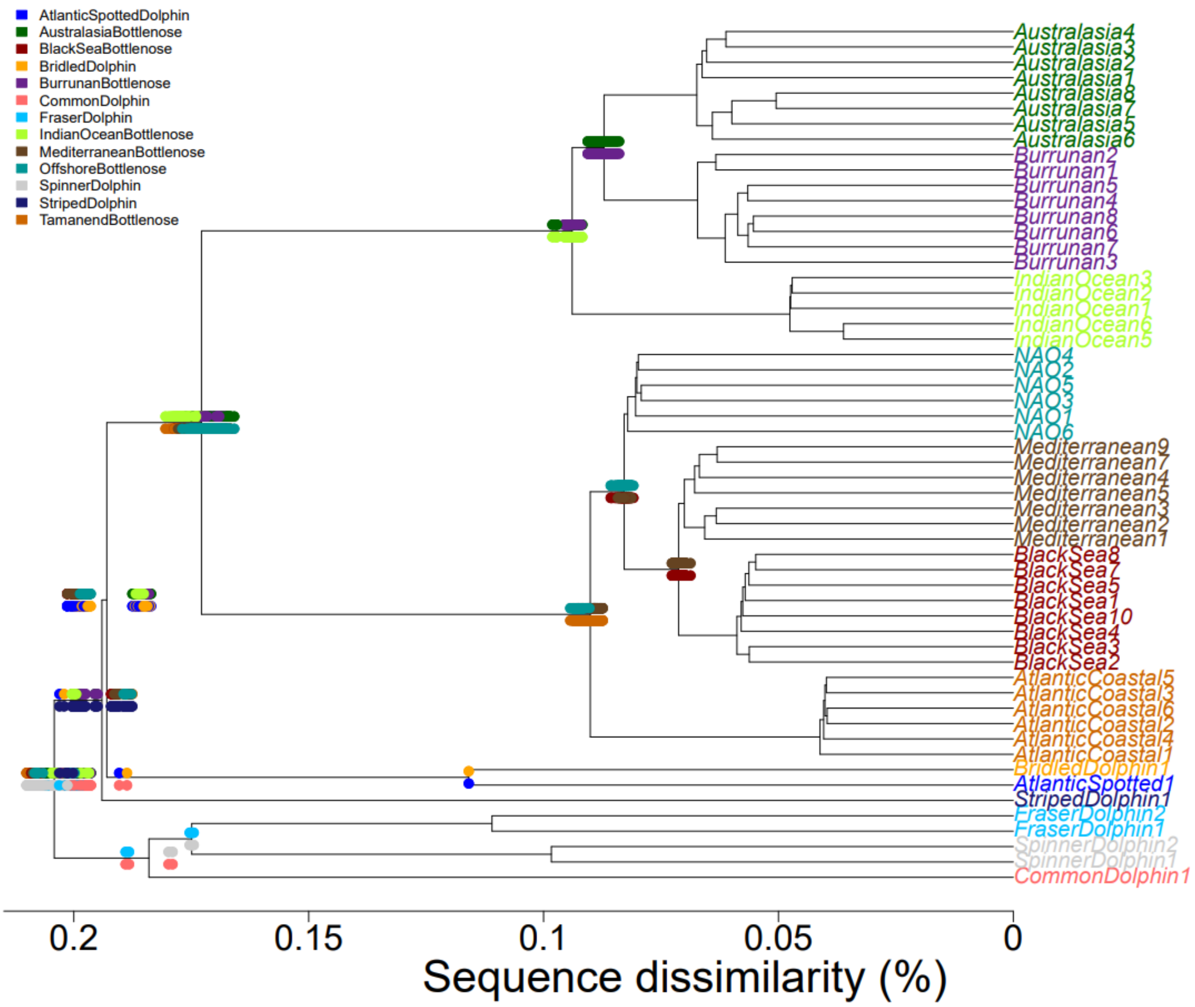


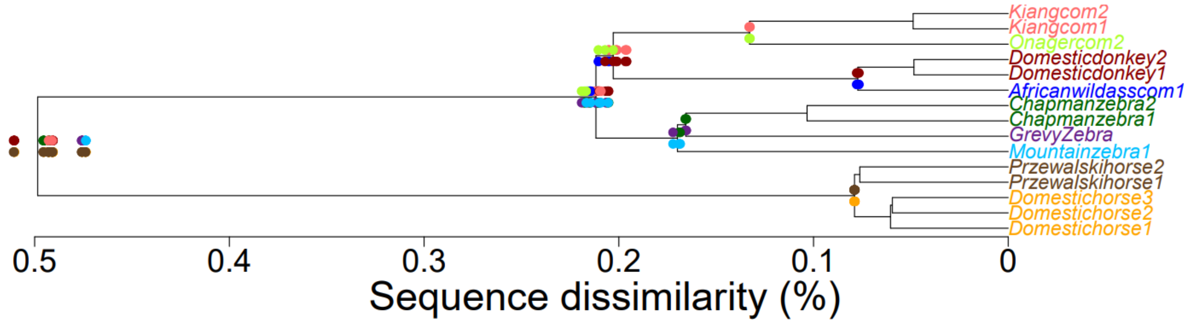


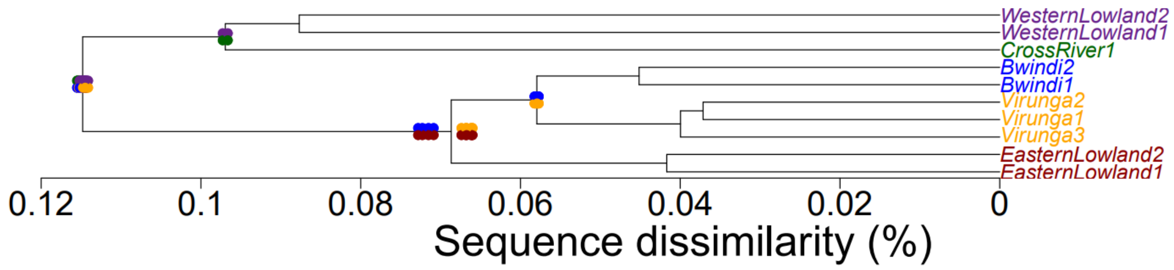


**Supplementary Figure 9 continued.** *Idem, for the datasets of dolphins (top), equids (middle), and gorillas (bottom).*

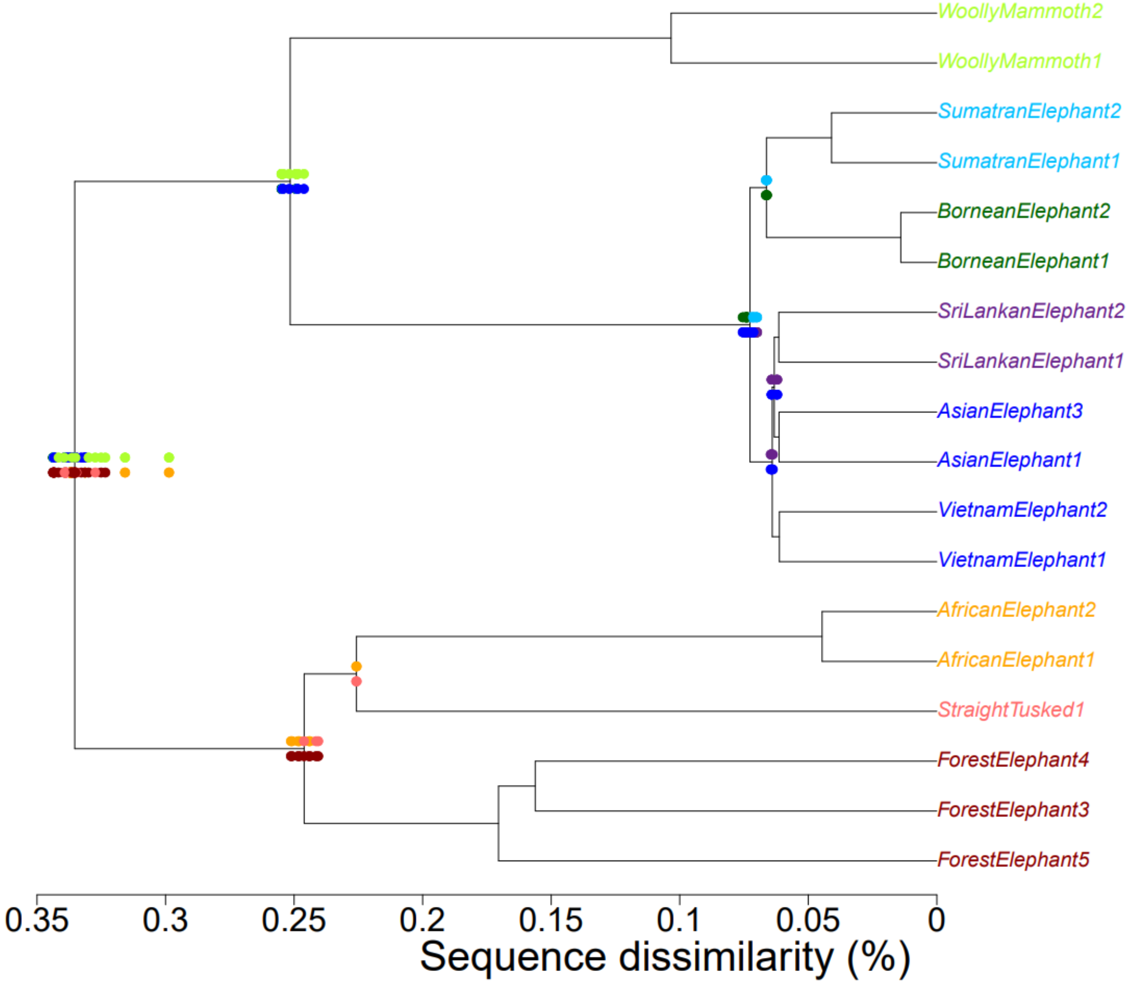


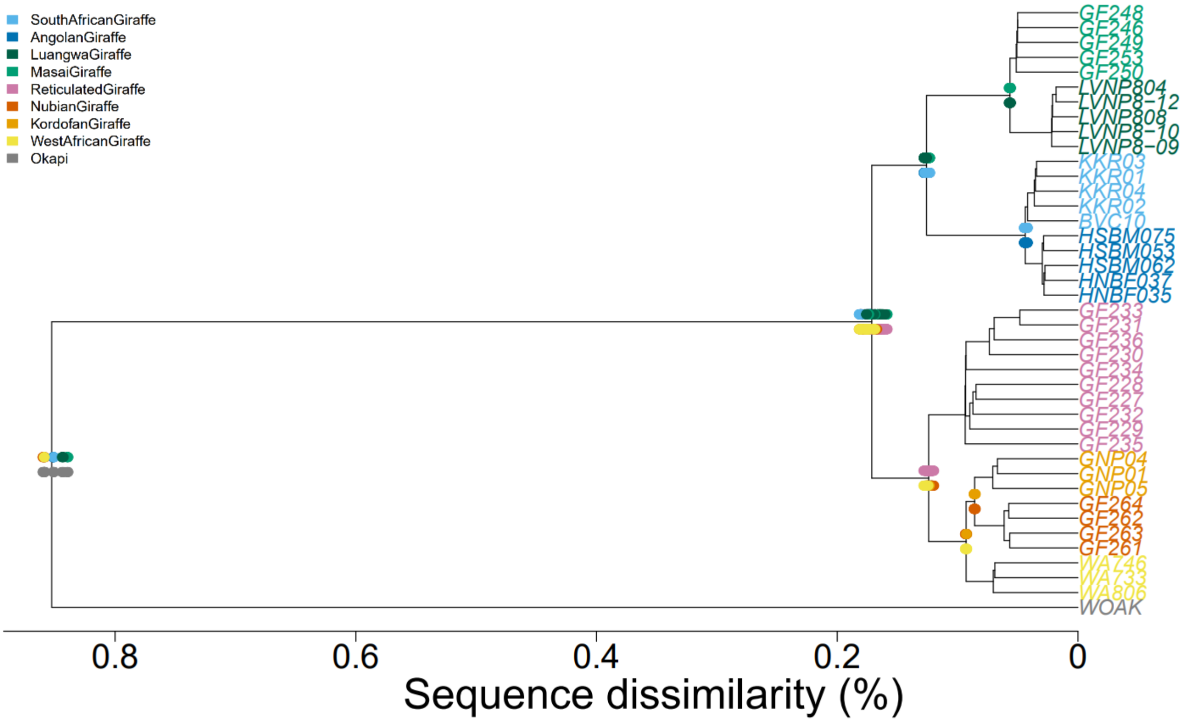


**Supplementary Figure 9 continued**. *Idem, for datasets of elephants (top) and giraffes.*

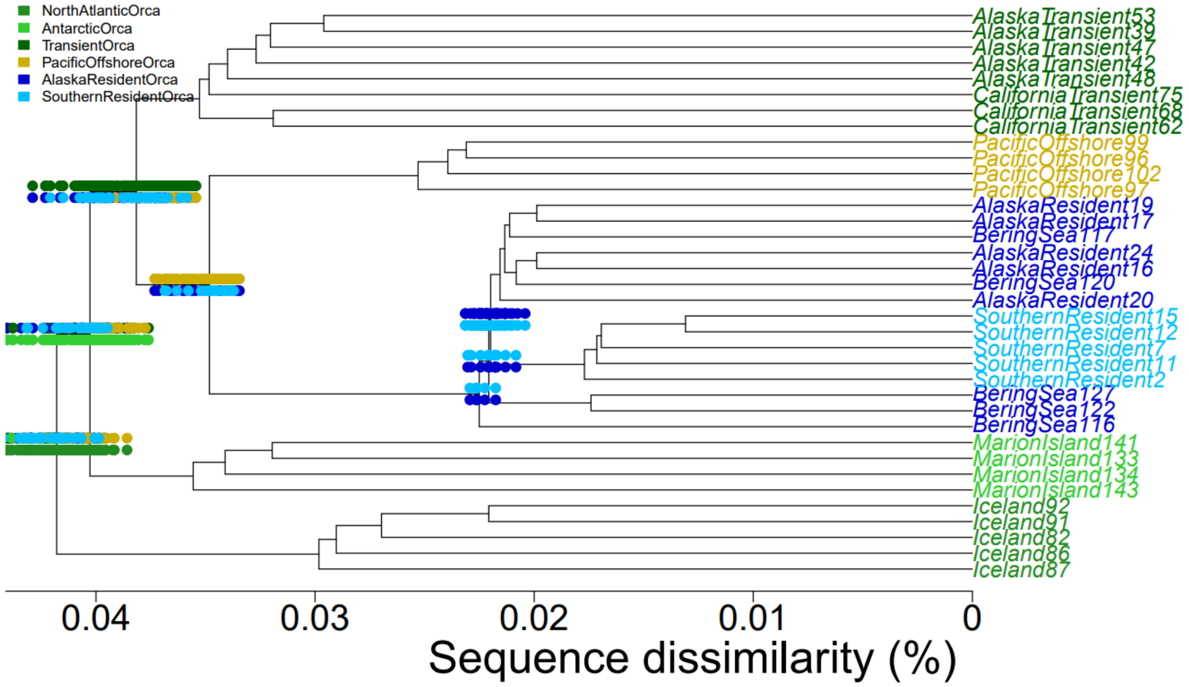


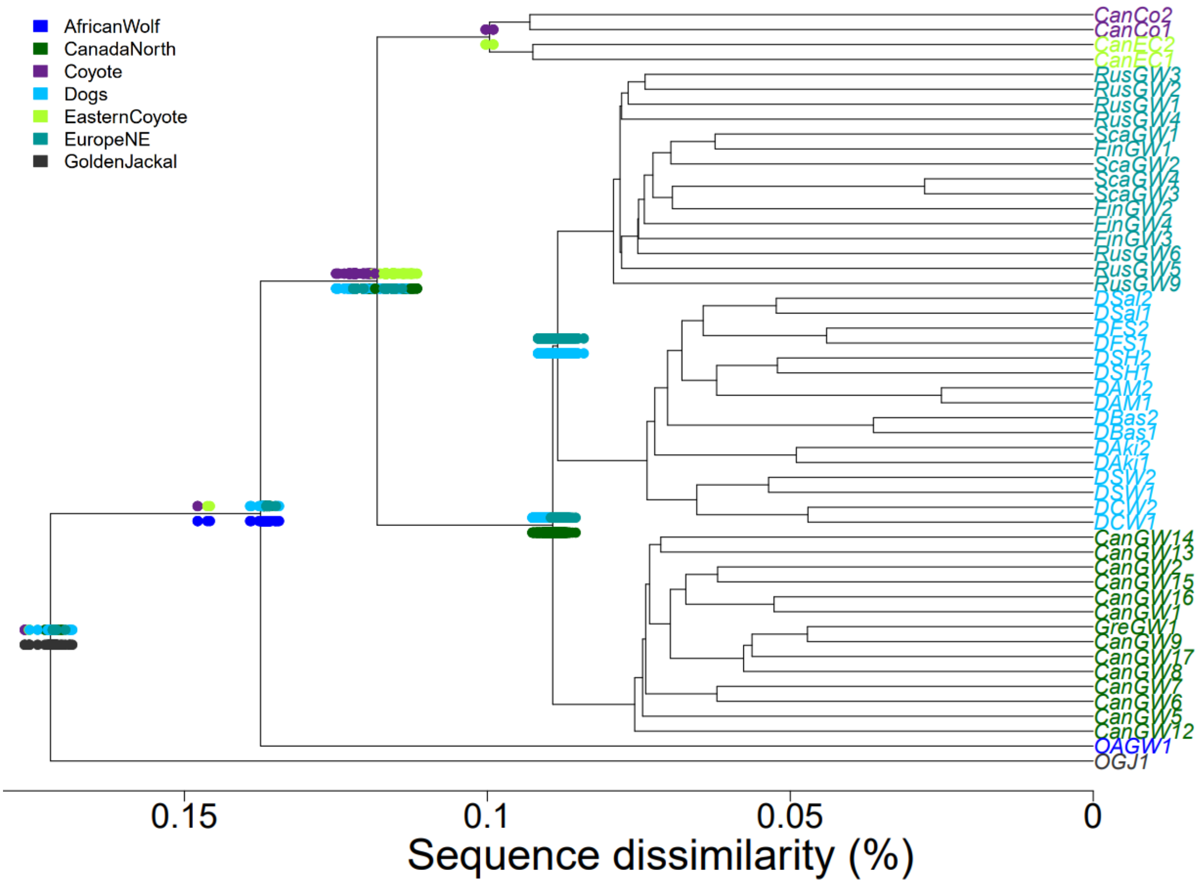


**Supplementary Figure 9 continued.** *Idem, for the datasets of killer whales (top) and wolves (bottom).*

299
